# The Morphology of α-Synuclein Fibrils Changes during Formation, Storage, and upon Exposure to Ligands

**DOI:** 10.1101/2025.08.29.672948

**Authors:** Timothy S. Chisholm

**Affiliations:** Yusuf Hamied Department of Chemistry, University of Cambridge, Cambridge, CB2 1EW, UK

## Abstract

Protein fibrils are pathological hallmarks of many neurodegenerative diseases, including Parkinson’s disease. The preparation of α-synuclein (αSyn) fibrils in vitro is widely used in research relating to these conditions. However, αSyn fibrils exhibit substantial structural polymorphism. How fibril morphology evolves during formation, storage, or in the presence of small molecule ligands is poorly understood. Here, the evolution of αSyn fibril morphology was investigated using fluorescence assays, circular dichroism, transmission electron microscopy, and ligand profiling. Fibril morphology was found to evolve continuously during both aggregation of αSyn and subsequent storage, even at -79 °C. The inclusion of ligands, such as Thioflavin X, during aggregation affected both the reaction kinetics and the fibril morphologies formed. The addition of ligands to pre-formed fibrils also produced changes in fibril morphology. These results highlight that αSyn fibrils are in equilibrium with their chemical environment and transition through a series of transient morphologies. Furthermore, these findings suggest the potential to use ligands to remodel fibril structure, possibly allowing for pathological morphologies to be converted to less pathological forms.

## INTRODUCTION

Neurodegenerative diseases, such as Alzheimer’s disease and Parkinson’s disease (PD), are one of the most significant health challenges faced by society.^1–4^ These conditions are characterized by the formation of fibrillar protein aggregates in the brain. The proteins that most commonly form pathological protein fibrils are amyloid-β and tau in AD,^5^ and α-synuclein (αSyn) in PD.^6,7^ αSyn is a 140 residue, intrinsically disordered peptide that appears to have a range of native functions that are not yet fully understood.^8–10^ Aggregates of αSyn are also found in other neurodegenerative conditions, most notably dementia with Lewy bodies and multiple system atrophy, that are collectively termed synucleinopathies.^11,12^ However, the precise morphology of fibrils appears to be disease-specific.^13–19^

The properties of these aggregates, and the mechanism by which they form, have been the focus of intensive research efforts. The preparation of protein fibrils *in vitro* is therefore routinely performed, although notably the physical, chemical, and biological properties of fibrils depends on their morphology.^20–23^ The aggregation of amyloidogenic proteins is a complex process involving multiple reaction steps including primary nucleation, secondary nucleation, and elongation.^24–27^ Kinetic models have been developed that can be used to study these processes,^27–29^ but our understanding of how specific morphologies arise during aggregation is limited.

Evidence also suggests that the structure of fibrils, once formed, may continue to change. Aβ fibrils formed *in vitro*, and in the brains of transgenic mice, undergo a maturation process that affects their structure.^30–33^ Tau fibrils appear to aggregate via a range of different morphologies.^34,35^

For αSyn, microscopy has suggested that changes to fibril morphology can occur once formed.^36,37^ Changes to fibrils can also occur during storage, with disaggregation sometimes observed during storage at 4 °C,^20,36,38^ whereas the effects of storage at -79 °C are not well studied.^39^

Small molecule ligands that bind protein fibrils are used as a research tool to study aggregates.^40,41^ Thioflavin T (ThT) is a commonly used ligand that exhibits an increase in fluorescence when bound to protein fibrils, and is frequently employed to monitor aggregation kinetics.^42^ Small molecule ligands are also a useful tool for characterizing fibril morphology. Determining the structure of fibrils is challenging and most accurately done using cryo-electron microscopy.^43^ Ligand profiling is a comparatively accessible method to characterize fibril morphologies based on their ligand-binding properties.^44,45^

Protein aggregates have a collection of ligand binding sites that are morphology-dependent. Different ligands and binding assays report on different binding sites. These binding sites, and the underlying morphology, can therefore be characterized with ligand binding assays. This ligand profiling approach is summarized in Figure 1 for a protein fibril with two types of binding sites (blue and orange circles). The solvatochromic reporting ligand R is added first and binds to both sites, with a resultant increase in fluorescence. The non-fluorescent competing ligand C is then added which binds only orange sites, leading to a partial decrease in fluorescence emission. The binding of R, and subsequent displacement by C, is dependent on the nature and abundance of the blue and orange sites. Performing assays with a combination of ligands can therefore report on multiple different binding sites and characterize the fibril morphology.

**Figure 1.**
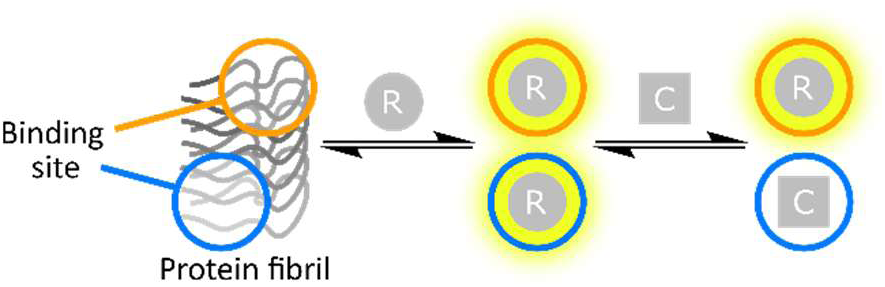
Competition binding assay performed using a protein fibril with two binding sites (blue and orange circles). A solvatochromic reporting ligand, R, binds to both sites leading to an increase in fluorescence. A competing ligand, C, is added which binds only blue sites, leading to a partial decrease in fluorescence.

Given the widespread use of fibrils formed *in vitro*, and the different properties that distinct polymorphs can possess, the formation of different morphologies during aggregation and storage could have a significant effect on the consistency and interpretation of research in this area. Additionally, as fluorescent ligands are routinely used to monitor aggregation, there is a need to better understand the effect of ligands on the aggregation process.

To gain a deeper understanding of the relationship between aggregation and morphology, this work aims to study three key questions. First, how does fibril morphology change during aggregation? Second, how does fibril morphology change during storage? Third, how do small-molecule ligands affect the aggregation and morphology of fibrils? This work employs a combination of fluorescence assays, circular dichroism (CD), transmission electron microscopy (TEM), and ligand profiling, to characterize aggregate morphology and answer these questions. Ultimately the morphology of αSyn fibrils was found to be constantly changing during aggregation and storage, and is sensitive to the presence of ligands. Rather than direct conversion of monomeric to a single polymorph, αSyn passes through a series of transient polymorphs and does not form a final stable polymorph under the conditions studied.

## RESULTS AND DISCUSSION

### Selection of a Panel of Ligands

A panel of ligands was first desired that could differentiate a wide range of fibril polymorphs by ligand profiling. Ligand profiling is most effective when structurally diverse ligands are used to sample a range of binding sites. Prior work has already identified several ligands that target distinct sites on αSyn fibrils.^44,45^ To create a larger panel of ligands, a molecular fingerprinting approach was used to select structurally diverse αSyn ligands. Molecular fingerprints are cheminformatics tools that encode molecular structures into a numerical, machine-readable format.^46,47^ These fingerprints can be compared to identify molecules with similar or distinct structures. Morgan fingerprints are the most commonly used, and describe the substructures surrounding each atom in a molecule.^48^

Using a database of amyloid-binding ligands,^40^ Morgan fingerprints were calculated for αSyn ligands with nanomolar dissociation constants. These ligands were visualized using t-distributed stochastic neighbor embedding (t-SNE) to map the chemical space (Figure 2a). The ligands ThX, CR, BTA, ThR, OXI, and S5H were selected based on prior evidence they target different binding sites (Figure 3).^40,44,45,49–51^ ThX is an analogue of ThT that exhibits a superior enhancement in fluorescence upon binding.^51^ Next, AAR, TPHN-1, IND, BF, FER, and XIA were selected to improve coverage of the chemical space (Figure 3).^52–56^ The coordinates of the selected ligands are annotated in the chemical space representation shown in Figure 2a. The resultant panel has five solvatochromic reporting ligands (in red) and seven non-solvatochromic competing ligands (in black, Figure 3). Pairwise Tanimoto similarity coefficients calculated using Morgan fingerprints confirmed that the ligands in this panel are structurally distinct, with a maximum coefficient of 0.42 between CR and ThR (Figure 2b).^57,58^ These ligands were prepared based on literature methods (see SI).^44,51–56,59–61^

**Figure 2.**
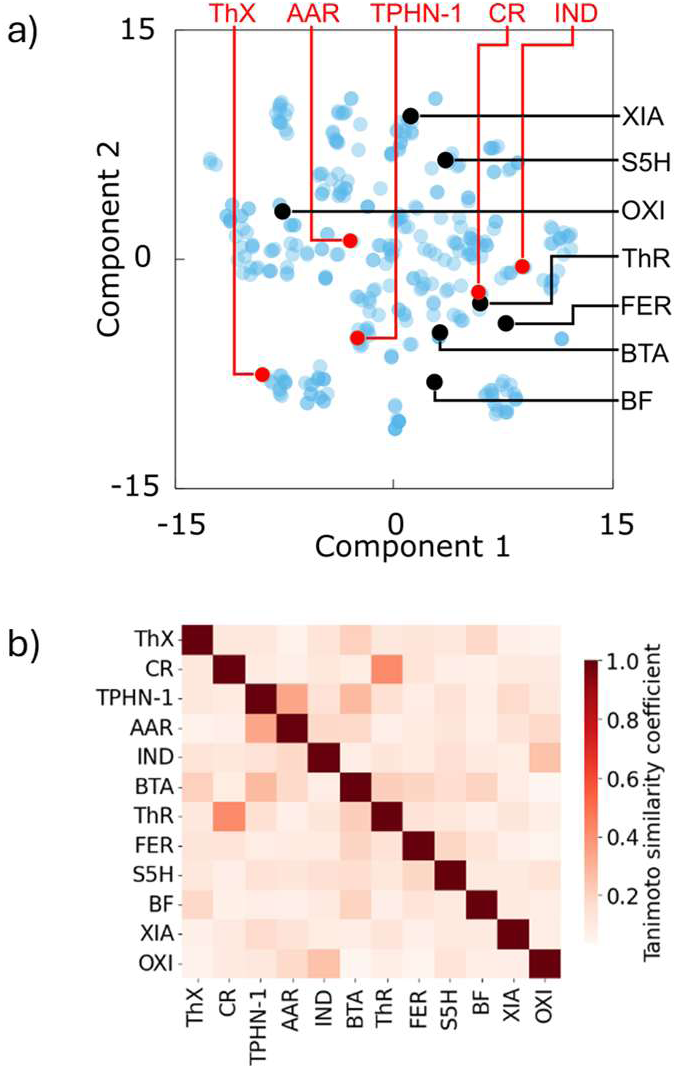
(a) t-SNE plot of Morgan fingerprints (radius 2) calculated for the database of αSyn ligands. (b) Heat map of pairwise Tanimoto similarity coefficients of ligands calculated using Morgan fingerprints (radius 2).

**Figure 3.**
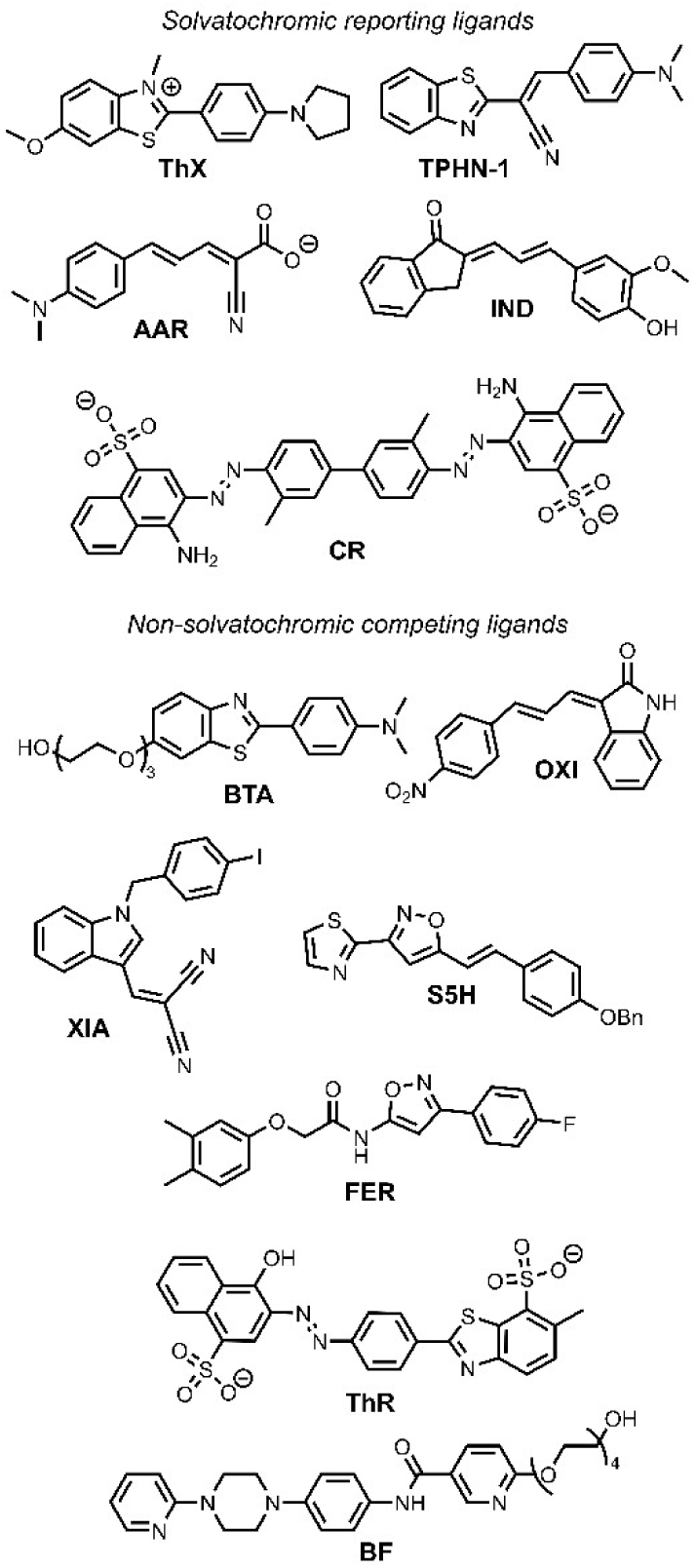
Structures of the αSyn ligands used in this study.

### Aggregation of αSyn

A standard aggregation protocol was followed to generate αSyn fibrils to study. A solution of αSyn monomer (50 µM) in 50 mM Tris with 1.5 mM NaN3 (pH 7.4) was prepared in a 1.5 mL Protein LoBind Eppendorf microtubule containing a PTFE magnetic stirrer bar (8 mm x 1.5 mm). The tube was sealed with parafilm and the solution was stirred at 50 rpm (37 °C). A single batch of monomer was used to ensure consistency across this experiment. Variation arising from different aggregation preparations is common and may obscure the changes in structural features that this study is aiming to measure.

For these experiments, no ligands were present in the aggregation mixture. Aggregation was performed with monomeric αSyn alone, and ligands were only added to aliquots taken at different time points for fluorescence measurements.

### Monitoring αSyn Morphology During Aggregation

The evolution of αSyn fibril morphology during aggregation was first studied. Aliquots of the aggregation mixture were taken after 24 h, 48 h, 72 h, 96 h, 1 week, 2 weeks, and 4 weeks, and subjected to analysis using fluorescent ligands, TEM, CD, and ligand profiling. Smaller aliquots were taken at more frequent intervals and analyzed only with fluorescent ligands to monitor aggregation kinetics. As no ligand was included in the aggregation mixture, reporting ligands were combined with the reaction aliquot.

The measured aggregation kinetics were dependent on the fluorescent ligand added. ThX and CR gave similar kinetic curves with a lag time (*tlag*: time to 10% of maximum signal) of 30 h, and a growth halftime (*t1/2*: time to 50% of maximum signal) of 40 h (Figure 4a). The ligands IND, AAR, and TPHN-1 gave a different kinetic profile with a *tlag* of 40 h and *t1/2* of 100 h (Figure 4b). This result indicates the formation of at least two distinct types of binding sites. Sites recognized by ThX and CR form earlier during aggregation, while sites recognized by AAR, TPHN-1, and IND form more slowly. This difference suggests that the fibril morphology, or ensemble of morphologies, changes over time. The fluorescence of all ligands decreased at later timepoints, indicating further changes in the binding sites.

**Figure 4.**
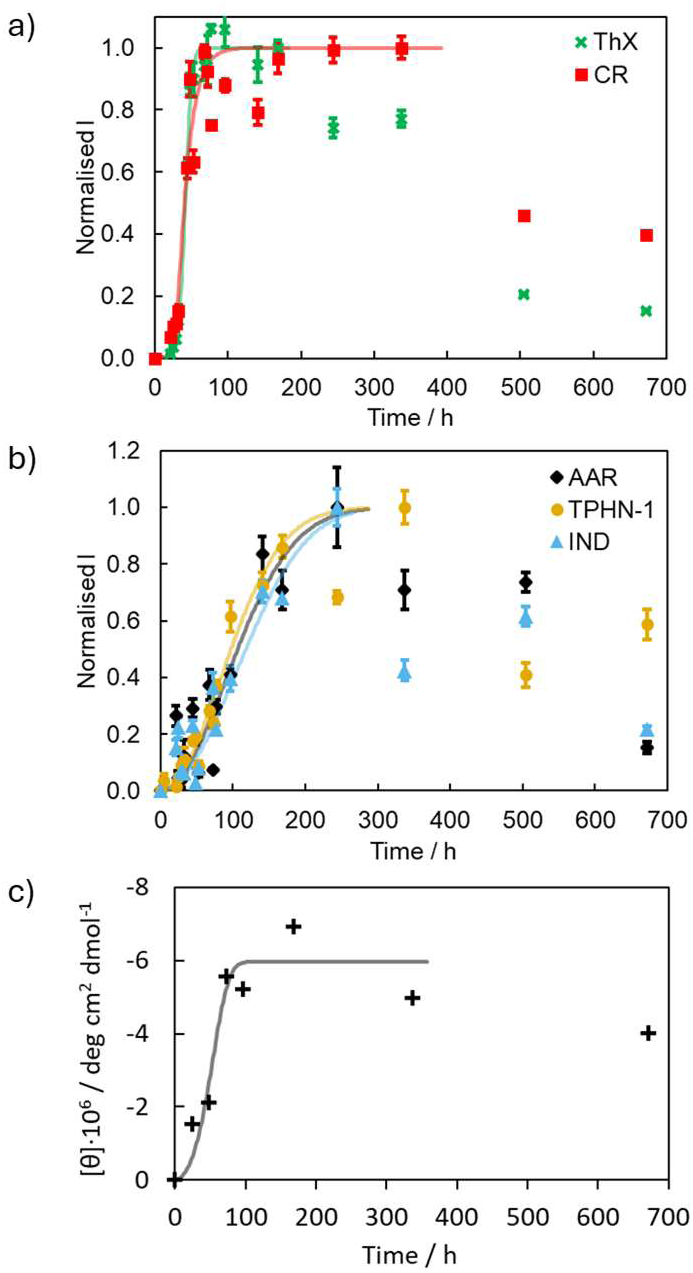
Aggregation kinetics of 50 µM αSyn in 50 mM Tris with 1.5 mM NaN3 (pH 7.4, 37 °C). Monitoring the aggregation process using (a) ThX (**×**), CR (▪), (b) AAR (◆), TPHN1 (⬤), or IND (▲) fluorescence. Aliquots were diluted to 1.0 µM αSyn in 1xPBS (pH 7.4) in the presence of fluorescent ligand (2.0 µM) and the fluorescence of the reporting ligand was measured. Data points are the average of at least three measurements with 95% confidence intervals shown. (c) Monitoring the aggregation process using CD. High molecular weight αSyn species were isolated from the aggregation reaction by centrifugal filtration (nominal molecular weight limit = 100 kDa) and spectra were recorded in 10 mM sodium phosphate buffer (pH 7.4). The signal measured at 220 nm is shown. Plotted lines are intended only as guides and were generated using AmyloFit.^28^

Aggregation kinetics were also followed by monitoring the CD band at 220 nm which is characteristic of β-sheet secondary structure (Figure 4c). The maximum concentration of β-sheets occurs after approximately 100 h, with a gradual decrease at later timepoints. By TEM, fibrils were observed as early as 24 h (Figure 5a-f). Over time these fibrils fragmented and became wider with a more heterogeneous width distribution (Figure 5g-i, Figure S50). From 72 h to 2 weeks the number of fibrils decreased and small, non-fibrillar aggregates appeared.

**Figure 5.**
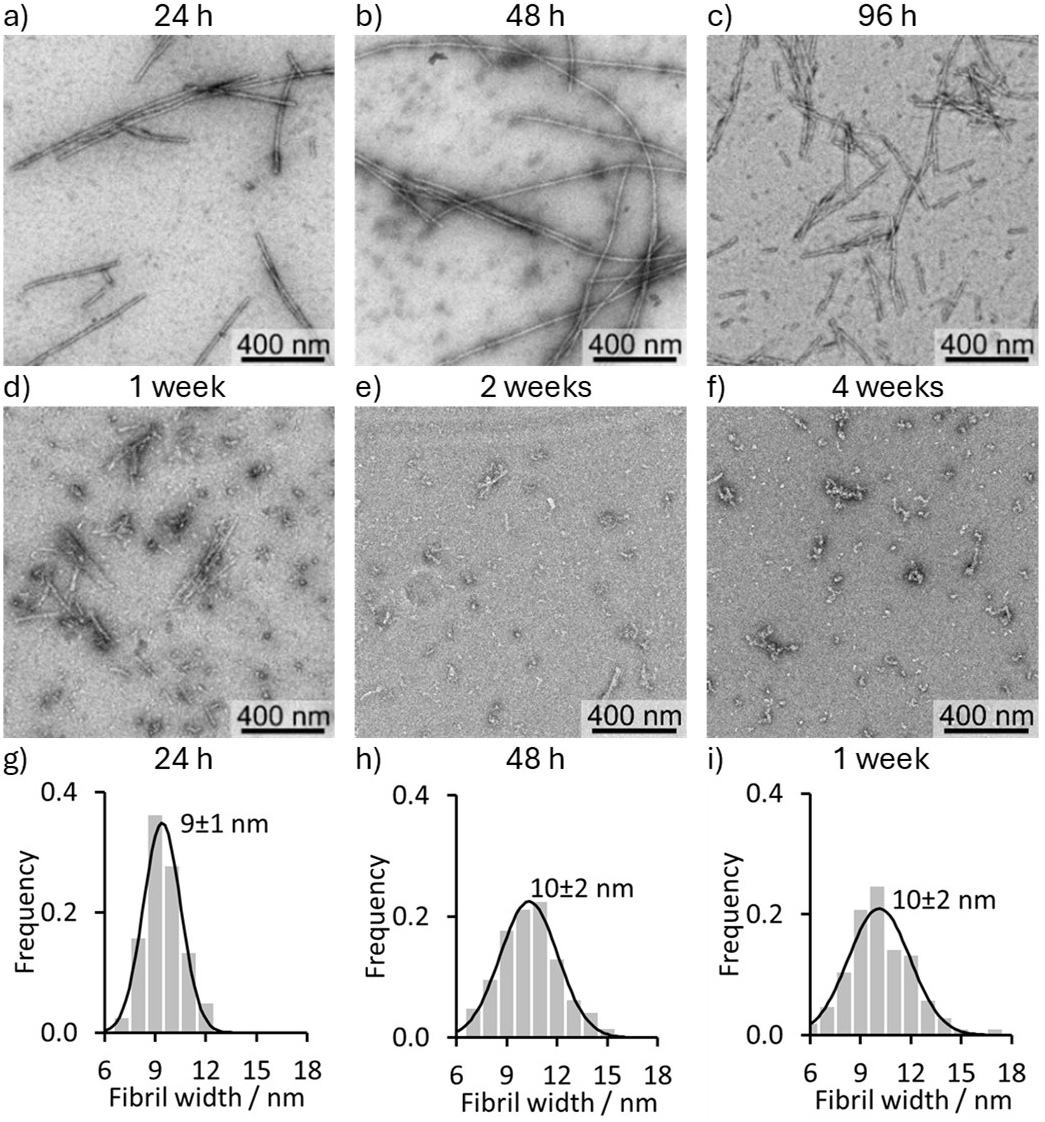
Representative TEM micrographs of αSyn aggregates obtained after (a) 24 h, (b) 48 h, (c) 96 h, (d) 1 week (168 h), (e) 2 weeks (336 h), and (f) 4 weeks (672 h) aggregation. The distribution of αSyn fibril widths of (g) 24 h, (h) 48 h, and (i) 1 week (168 h) aggregation timepoints. Fibril widths were measured as the full width half maximum (FWHM) in ImageJ,^62^ and at least 100 measurements from multiple representative TEM micrographs were taken for each timepoint. Plotted black lines are the Gaussian distribution corresponding to the mean ± standard deviation shown on each graph. Aggregation was performed with 50 µM αSyn in 50 mM Tris and 1.5 mM NaN3 (pH 7.4, 37 °C).

Ligand profiling was used to characterize the binding sites in more detail. Samples of the aggregation mixture were combined with 2 µM fluorescent ligand then 5 µM of competing ligand.

When combined with the 96 h αSyn aggregation timepoint, each reporting ligand had a unique response to the competing ligands (Figure 6). For example, ThX was displaced by ThR, BTA, and BF, whereas CR was only displaced by BTA. The responses of AAR, TPHN-1, and IND were similar, with smaller differences between the competing ligands BTA, S5H and BF. These differences in responses indicate the reporting ligands target different sets of binding sites. The competing ligands then target a subset of these sites.

**Figure 6.**
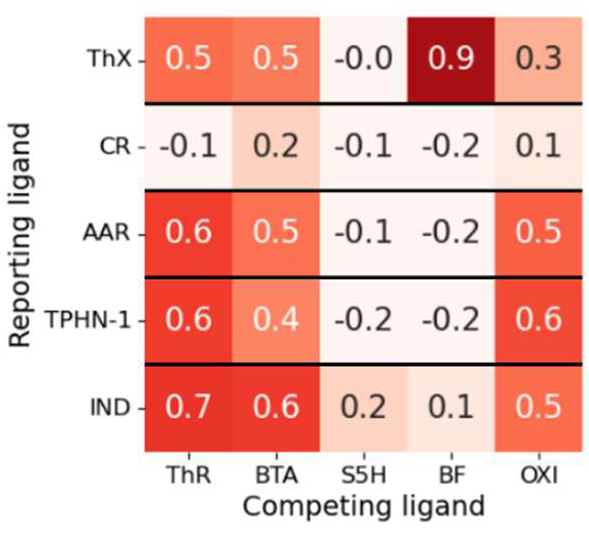
Heatmap showing the proportion of reporting ligand that is displaced by competing ligands from 96 h αSyn aggregates.

When these assays were repeated across aggregation timepoints the proportion of ThX and IND displaced by BTA, ThR, and BF changed (Figure 7). These changes in ligand binding reflect changes in the binding sites, and therefore fibril morphology, that occur during aggregation.

**Figure 7.**
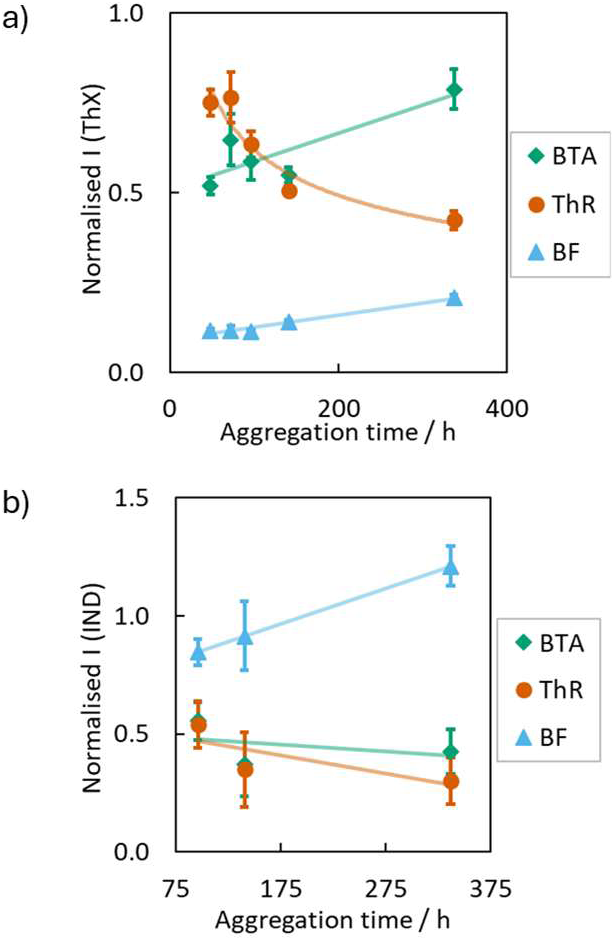
Fluorescence measurements for the displacement of (a) ThX (λex = 440 nm, λem = 488- 492 nm) and (b) IND (λex = 430 nm, λem = 565-570 nm) by competing ligands from αSyn fibrils at different aggregation timepoints. Aggregation was performed using 50 µM αSyn in 50 mM Tris with 1.5 mM NaN3 (pH 7.4, 37 °C). The reaction was monitored by diluting aliquots to 1.0 µM αSyn in 1xPBS (pH 7.4) and adding 2.0 µM reporting ligand and 5.0 µM competing ligand (BTA: ; ThR: ⬤; BF: ▴). Data points are the average of at least three measurements with 95% confidence intervals shown. Plotted lines are intended only as guides.

In summary, fluorescence assays, CD, TEM, and ligand profiling each provide distinct and complementary measurements of morphological changes that occur during αSyn aggregation. Fibrils were observed after only 24 h aggregation by TEM, whereas the lag time by fluorescence and CD was approximately 30 h. ThX and CR appear to report on binding sites that form earlier during aggregation than those of AAR, TPHN-1, and IND, and give a signal that correlates to the formation of β-sheets as determined by CD.

From 72 h to 2 weeks the number of fibrils decreased by TEM and small, non-fibrillar aggregates appeared, despite the fluorescent signal of ThX and CR remaining constant. The fluorescence emission of AAR, TPHN-1, and IND continued to increase, suggesting these ligands may report primarily on smaller aggregates or more mature fibril structures. The binding site distribution, assessed using ligand profiling, also continued to change. After 2 weeks few fibrils remained, coinciding with a decrease in fluorescence from all reporting ligands, although CD measurements still suggested a relatively high concentration of β-sheets.

These data highlight that a range of polymorphs exist in equilibrium and continue to change during both aggregation and maturation. These structural changes are not detectable by every analytical technique used. A comprehensive assessment of morphology therefore requires the use of multiple analytical methods.

### Storage of αSyn Fibrils Under Different Conditions

The effects of storage on αSyn fibril morphology were then assessed, as fibrils are often stored for weeks or months prior to use. Aliquots of αSyn fibrils after 96 h of aggregation were stored at -79 °C, 4 °C, and 37 °C for 25 days. These samples were then analysed using fluorescent ligands, TEM, CD, and ligand profiling.

Fluorescence measurements revealed clear changes after storage (Figure 8). Samples of fibril stored at -79 °C, 4 °C, and 37 °C were combined with each reporting ligand. The resulting fluorescence was compared with measurements taken using 96 h αSyn fibrils before storage. After storage at -79 °C the fluorescence of added ThX and CR was unchanged, whereas the fluorescence of added AAR and TPHN-1 decreased. After storage at 4 °C fluorescence from all ligands was reduced, with ThX showing less than 10% of its original fluorescence. After storage at 37 °C ThX fluorescence was also the most affected, CR and AAR fluorescence was halved, and TPHN-1 fluorescence did not change. The fluorescence of IND did not increase in the presence of any of the stored fibrils.

**Figure 8.**
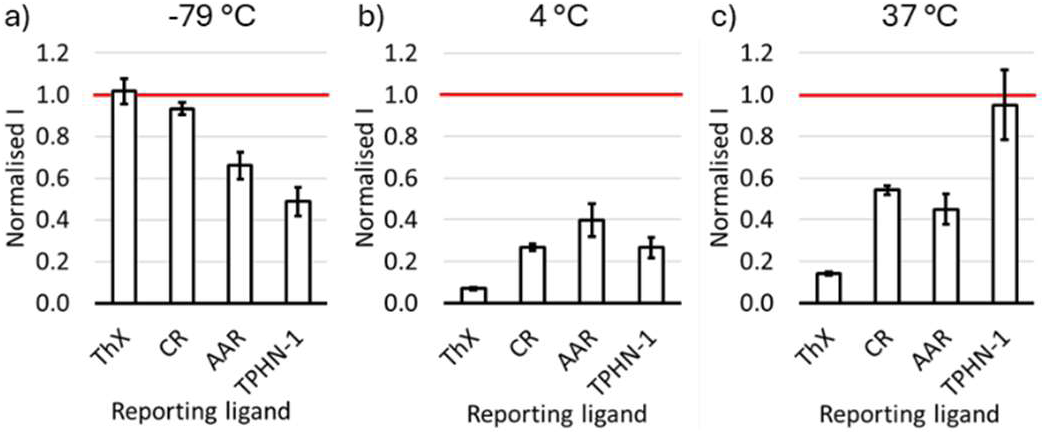
Fluorescence emission of reporting ligands (2.0 µM) bound to aliquots of 96 h αSyn aggregates stored for 25 days at (a) -79 °C, (b) 4 °C, and (c) 37 °C. Fluorescence data are normalised to the signal measured for the reporting ligands bound to 96 h αSyn aggregates pre-storage, represented by the plotted red line. Datapoints are the average of at least three measurements, with 95% confidence intervals shown.

TEM micrographs also revealed changes in fibril morphology (Figure 9). For storage at 37 °C only small, non-fibrillar aggregates were observed (Figure 5a). Fibrils were still present after storage at both 4 °C and 37 °C, and were wider (12±2 nm, 14±2 nm) than the initial 96 h αSyn aggregates (10±2 nm) (Figure 9a-b). However, the CD spectra of fibrils stored at 4 °C showed a weaker negative band around 220 nm and no positive band around 200 nm, suggesting a decrease in β-sheet character and a possible increase in random coil structures (Figure S47).

**Figure 9.**
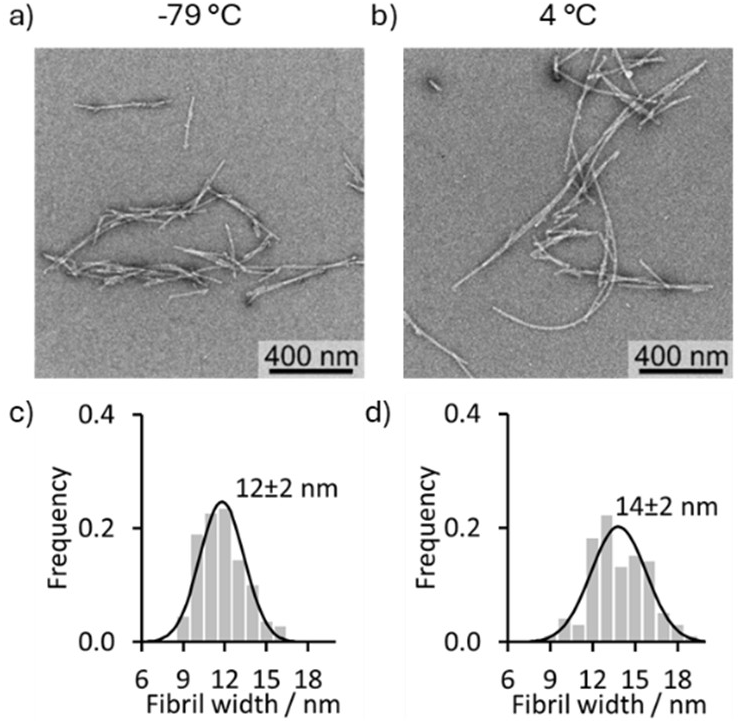
Representative TEM micrographs of 96 h αSyn aggregates after storage for 25 days at (a) -79 °C and (b) 4 °C. The distribution of fibril widths of 96 h αSyn aggregates after storage for 25 days at (c) -79 °C and (d) -4 °C. Fibril widths were measured as the full width half maximum (FWHM) in ImageJ,^62^ and at least 100 measurements from multiple representative TEM micrographs were taken for each timepoint. Plotted black lines are the Gaussian distribution corresponding to the mean ± standard deviation shown on each graph.

Ligand profiling supported that a change in fibril morphology had occurred during storage (Figure 10). Binding site changes were quantified using ΔNorm, defined as the change in the proportion of reporting ligand displaced by a competing ligand between one sample containing 96 h αSyn fibrils that had been stored, and a second sample containing 96 h αSyn fibrils pre-storage. For fibrils stored at -79 °C, significant changes in displacement were only observed for the reporting ligands AAR and TPHN-1. Less AAR was displaced by ThR, BTA, and OXI from the stored fibrils, while more TPHN-1 was displaced by FER and less by OXI.

**Figure 10.**
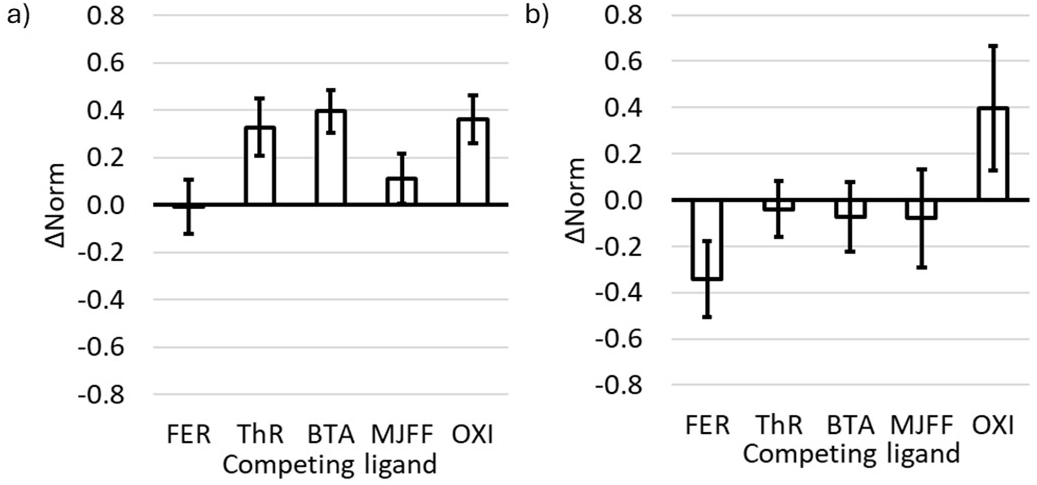
The change in the amount of (a) AAR and (b) TPHN-1 displaced by competing ligands from 96 h αSyn aggregates after storage at -79 °C for 25 days. ΔNorm is calculated as the difference in proportion of reporting ligand displaced between one sample containing 96 h αSyn fibrils that had been stored, and a second sample containing 96 h αSyn pre-storage. Datapoints are the average of at least three measurements, with 95% confidence intervals shown.

These data demonstrate that αSyn fibril morphology changed under each storage condition explored. These changes were not apparent using every assay format. For example, similar fluorescence was measured when ThX was added to fibrils before and after storage at -79 °C, yet TEM and other ligand assays revealed changes in fibril structure.

To assess the effects of long-term storage a second sample of αSyn fibrils was analysed. These fibrils were prepared under similar conditions (50 mM Tris, pH 7.4, 37 °C), but in the absence of NaN3. Samples were stored at -79 °C for 450 days (1 year and 3 months) and 606 days (1 year and 8 months). When these samples were combined with ThX, the measured fluorescence was lower for the fibrils that had been stored longer (Figure 11a), and changes in binding sites were observed by ligand profiling (Figures S70-S73). TEM analysis revealed a high number of thin fibrils was present in the sample stored for 1 year and 8 months (Figure 11b-c, Figure S49). While fibrils can therefore be stored for long time periods without disaggregation or conversion into non-fibrillar aggregates, their morphology appears to continue to change.

**Figure 11.**
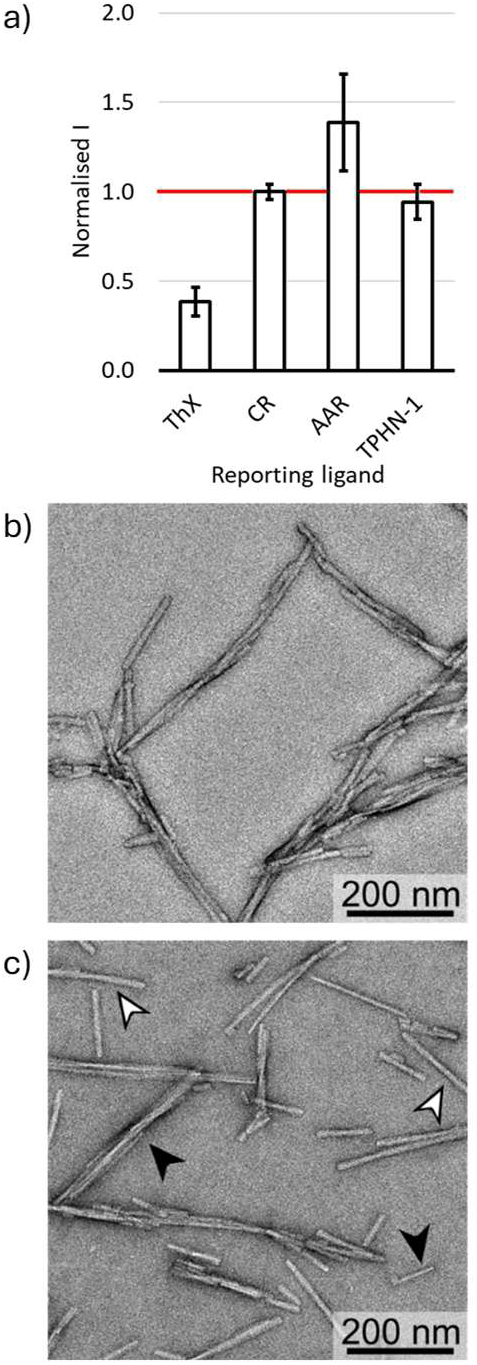
a) Fluorescence emission of reporting ligands (2.0 µM) bound to αSyn aggregates after storage at -79 °C for 1 year and 8 months, normalised to the fluorescence measured when bound to αSyn aggregates after storage at -79 °C for 1 year and 3 months, represented by the plotted red line. Datapoints are the average of at least three measurements, with 95% confidence intervals shown. Representative TEM micrographs of αSyn aggregates after storage at -79 °C for (b) 1 year and 3 months, and (c) 1 year and 8 months. Black arrowheads identify thin fibrils, and white arrowheads identify thick fibrils.

The impact of freeze-thaw cycles was also studied. Samples of αSyn aggregates were repeatedly thawed in room temperature water then refrozen at -79 °C in an acetone bath containing dry ice. When these aggregates were combined with reporting ligands, the observed fluorescence emission depended on the number of freeze-thaw cycles the sample had been subjected to (Figure 12).

**Figure 12.**
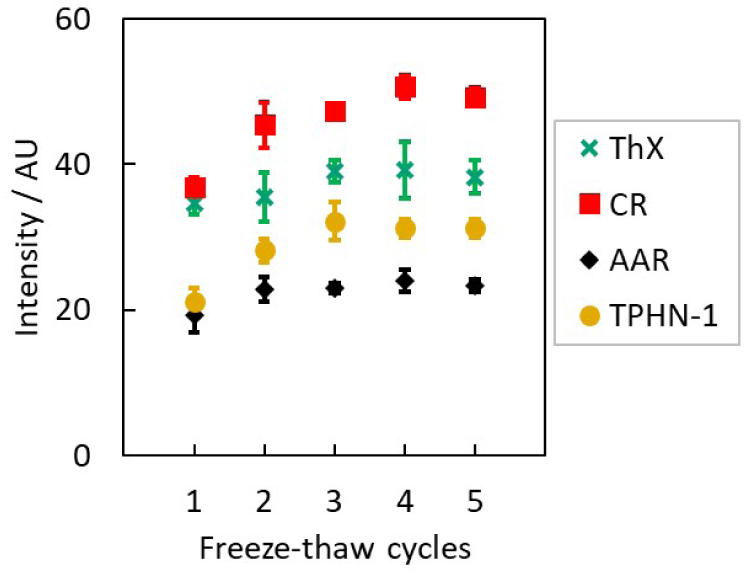
The fluorescence intensity of ThX (**×**), CR (▪), AAR (◆), or TPHN-1 (⬤) (2.0 µM) bound to αSyn aggregates after storage at -79 °C for 1 year and 8 months years and different numbers of freeze-thaw cycles. Data points are the average of at least three measurements with 95% confidence intervals shown.

Reliable storage of αSyn fibrils is therefore a challenge. Changes to fibril morphology occur under common storage conditions, although samples stored at -79 °C produced the smallest changes. Repeated freeze-thaw cycles also had a relatively minor impact on fibril morphology. In all cases these changes introduce a source of variability in experiments using stored fibrils, particularly if aliquots are taken and used at different storage times. Fibrils should ideally be used without large differences in storage time, and with a consistent number of freeze thaw cycles.

### Aggregation in the Presence of ThX

Monitoring αSyn aggregation is commonly achieved through the addition of ThT into the aggregation mixture. The results above demonstrate that using a single fluorescent ligand has limitations for monitoring the aggregation process. Additionally, whether inclusion of a ligand during the aggregation process affects the outcome is not well understood. The binding of a ligand to aggregates or intermediates will affect the energy landscape of the aggregation process and could influence reaction kinetics and the polymorphs formed. To determine the impact of a ligand on the aggregation process, the αSyn aggregation protocol was repeated in the presence of 20 µM ThX. Timepoints were analysed using ThX fluorescence, TEM, and ligand profiling.

The inclusion of ThX produced both a shorter lag time and more rapid conversion to aggregated product, with *tlag* of 18 h and *t1/2* of 22 h (Figure 13). Compared to the fibrils formed in the absence of ThX during aggregation, a higher ThX fluorescence was measured followed by a more rapid decrease in fluorescence with time. TEM micrographs also demonstrate that the inclusion of ThX leads to fibrils that fragment less and retain greater fibrillar character over time at 37 °C (Figure 14a-f). Ligand profiling nevertheless demonstrated that binding sites continued to change, with the amount of ThX displaced by OXI and ThR decreasing over time even as overall ThX fluorescence remained constant (Figure 14g). The ligand profile observed for αSyn aggregates after 96 h was also different compared to that for αSyn aggregates formed without ThX (Figure 14h).

**Figure 13.**
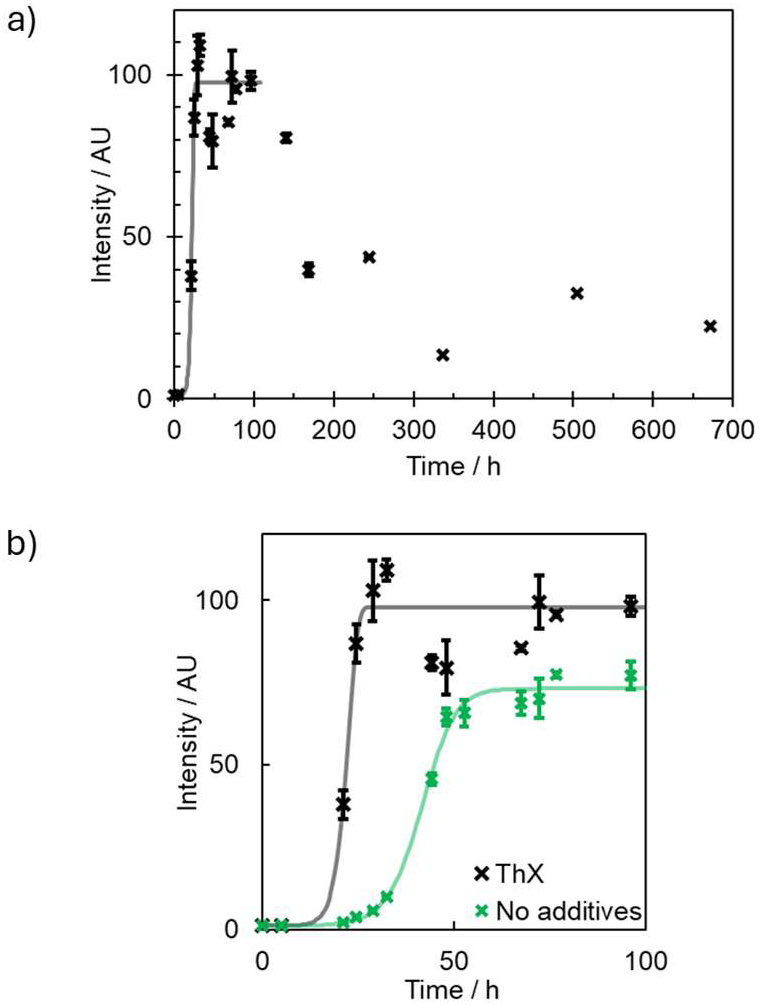
(a) Aggregation kinetics of 50 µM αSyn in 50 mM Tris with 20 µM ThX and 1.5 mM NaN3 (pH 7.4, 37 °C). The reaction was monitored by diluting aliquots to 1.0 µM αSyn in 1xPBS (pH 7.4) and adding ThX to a final concentration of 2.0 µM (λex = 440 nm, λem = 488-492 nm). Data points are the average of at least three measurements with 95% confidence intervals shown. (b) Comparison of the aggregation kinetics in the presence (**×**) and absence of ThX (**×**) from Figure 4.

**Figure 14.**
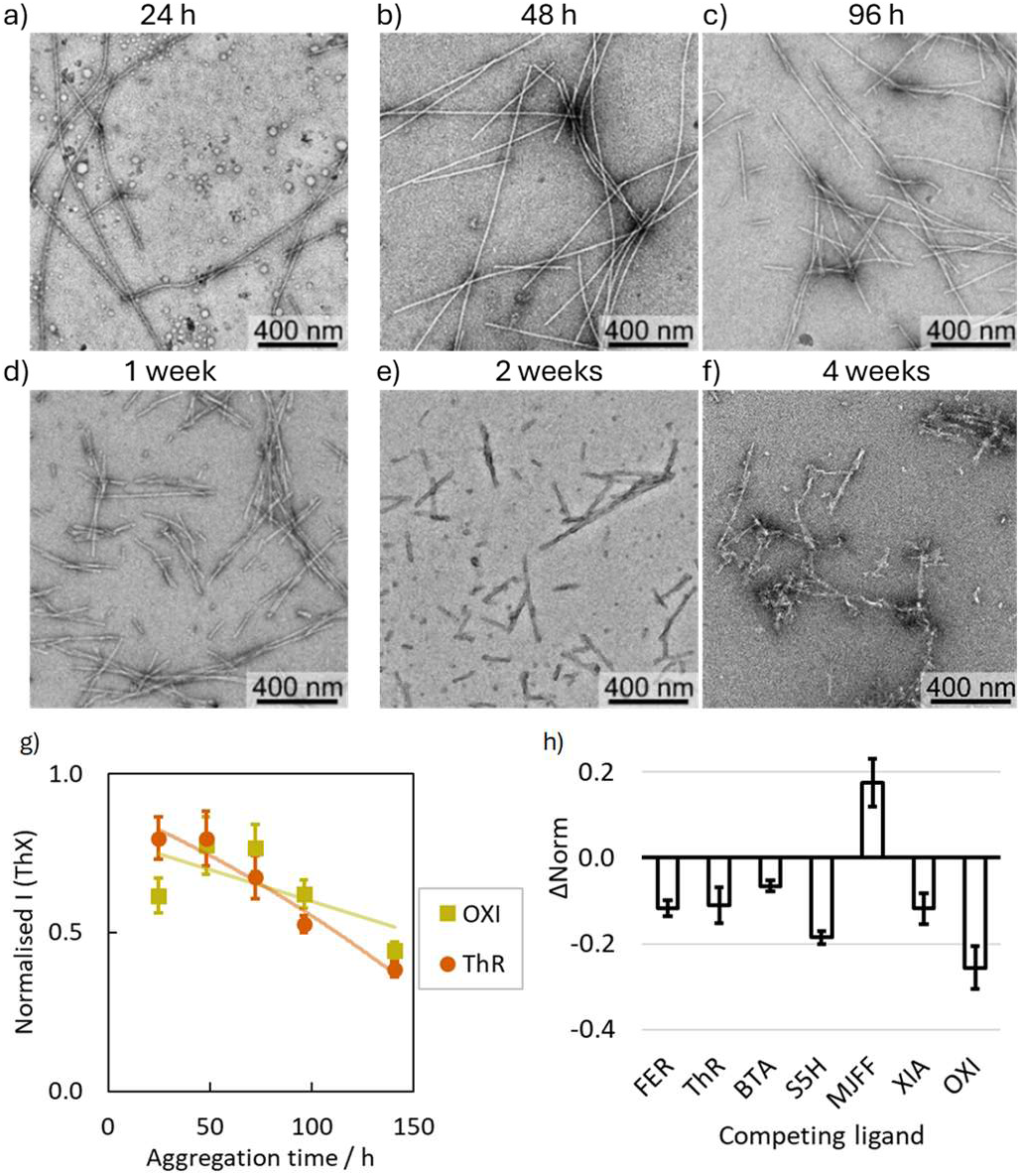
Representative TEM micrographs of αSyn aggregates obtained after (a) 24 h, (b) 48 h, (c) 96 h, (d) 1 week (168 h), (e) 2 weeks (336 h), and (f) 4 weeks (672 h) aggregation. Aggregation was performed using 50 µM αSyn in 50 mM Tris with 20 µM ThX and 1.5 mM NaN3 (pH 7.4, 37 °C). (g) Fluorescence measurements for the displacement of ThX (λex = 440 nm, λem = 488-492 nm) from αSyn fibrils formed in the presence of ThX. Aggregation was performed using 50 µM αSyn in 50 mM Tris with 20 µM ThX and 1.5 mM NaN3 (pH 7.4, 37 °C). The reaction was monitored by diluting aliquots to 1.0 µM αSyn in 1xPBS (pH 7.4) and adding 2.0 µM ThX and 5.0 µM competing ligand (OXI: ▪; ThR: ⬤). Data points are the average of at least three measurements with 95% confidence intervals shown. Plotted lines are intended only as guides. (h) The difference in the proportion of ThX displaced by competing ligands from αSyn aggregates prepared in the presence and absence of ThX. ΔNorm is calculated as the difference in proportion of ThX displaced between one sample containing 96 h αSyn fibrils that had been aggregated in the presence of ThX, and a second sample containing 96 h αSyn fibrils that had been aggregated in the absence of ThX. Datapoints are the average of at least three measurements, with 95% confidence intervals shown.

The 96 h αSyn fibrils formed in the presence of ThX were also more stable in cooled storage. While ThX fluorescence decreased after the aggregates were stored at 4 °C, the intensity did not changed after the aggregates were stored at -79 °C (Figure 15a), and long, homogeneous fibrils were observed in both cases (Figure 15b-c). Changes to the ligand profile were also observed after storage at 4 °C and 37 °C, but not after storage at -79 °C (Figures S59, S61).

**Figure 15.**
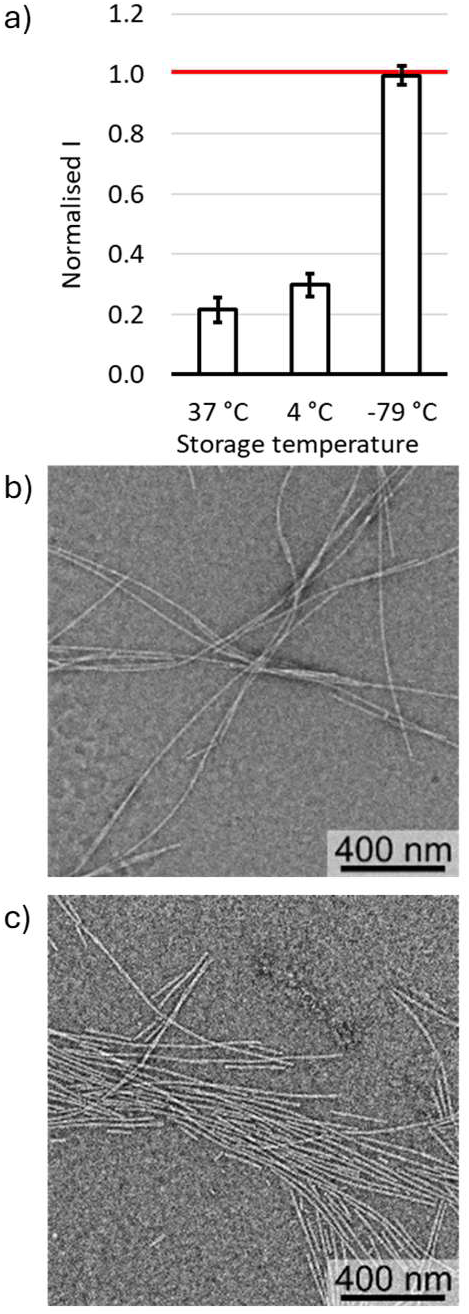
Characterisation of 96 h αSyn aggregates formed in the presence of ThX after the aggregates were stored for 25 days at different temperatures. (a) Fluorescence emission of ThX (2.0 µM) and 96 h αSyn aggregates after the aggregates were stored at 37 °C, 4 °C, or -79 °C for 25 days. Fluorescence data are normalised to the signal measured for ThX bound to αSyn aggregates pre-storage, represented by the plotted red line. Datapoints are the average of at least three measurements, with 95% confidence intervals shown. Representative TEM micrographs of the 96 h αSyn aggregates after storage for 25 days at (b) 4 °C, and (c) -79 °C.

These results demonstrate that the inclusion of ThX during aggregation affects both aggregation kinetics and the morphology of the fibrils formed. The fibrils formed in the presence of ThX were more stable than those formed in its absence. Some fibrillar material still remained after storage at 37 °C, and numerous large fibrils were observed after storage at 4 °C or -79 °C. Notably, no changes to the ligand-binding properties of these αSyn aggregates formed in the presence of ThX were observed when stored at -79 °C.

### Incubation of αSyn Fibrils with Fluorescent Ligands

Given that ThX can affect the formation of αSyn fibrils, the effect of adding ligands to preformed fibrils was studied. Aliquots of 96 h αSyn aggregates were treated with 20 µM ThX, CR, AAR, TPHN-1, or IND, and incubated for 25 days at 37 °C. The fluorescent signal and ligand profile for each sample was then measured.

The fluorescence emission of reporting ligands was compared when added to αSyn fibrils before incubation, and after incubation. Following incubation, ThX and IND exhibited decreased fluorescence suggesting a change in either fibril concentration or the binding sites present (Figure 16a). No change was measured for CR after incubation, whereas the fluorescence of both AAR and TPHN-1 increased. Ligand profiling revealed that the ThX binding sites had changed following incubation (Figure 16b), whereas the ligand profiles for CR, AAR, TPHN-1, and IND were unaffected (Figures S65-S69).

**Figure 16.**
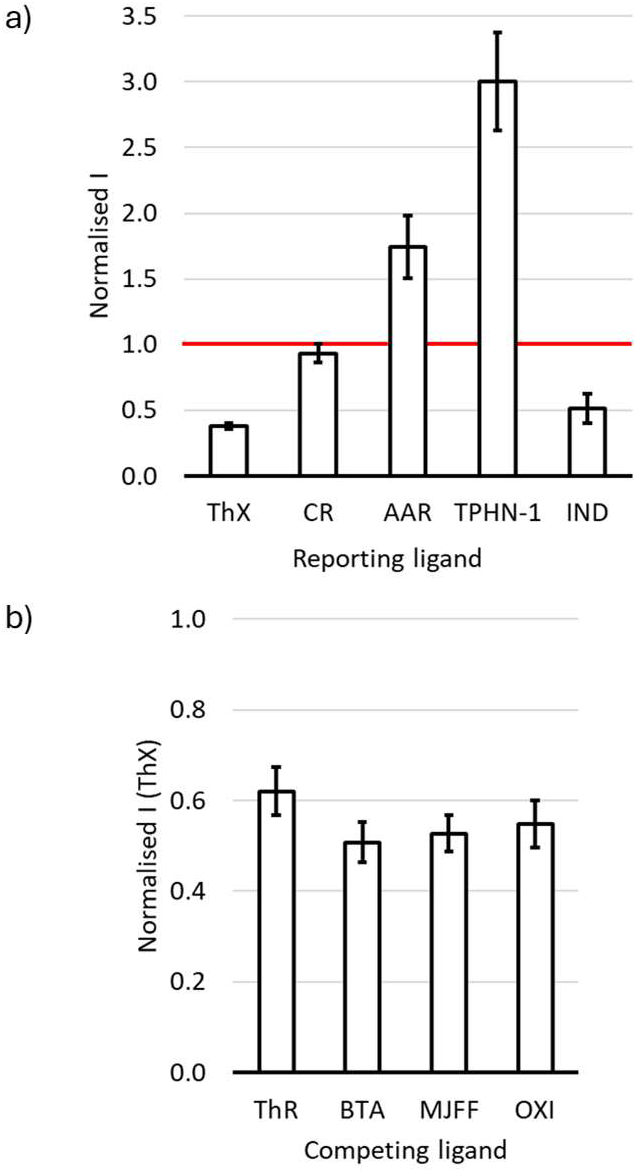
(a) Fluorescence emission of different reporting ligands after incubation with 96 h αSyn aggregates for 25 days at 37 °C. Fluorescence data are normalised to the signal measured for the reporting ligands bound to 96 h αSyn aggregates pre-incubation, represented by the plotted red line. (b) The proportion of ThX displaced by competing ligands from αSyn aggregates that have been incubated with 20 µM ThX for 25 days at 37 °C. Datapoints are the average of at least three measurements, with 95% confidence intervals shown.

The morphology of αSyn fibrils can therefore be influenced by the introduction of ligands. This observation is in line with αSyn fibrils existing in equilibrium with, and being affected by, their surrounding chemical environment.

## CONCLUSIONS

This work demonstrates that the morphology of αSyn fibrils continually changes during aggregation and under common storage conditions. Fibril morphology is also affected by the presence of small molecule ligands, added either during aggregation or once complete aggregation has already occurred. αSyn fibrils therefore appear to transition through a series of polymorphs over time. This process required the use of several analytical techniques in tandem to detect. Fluorescence assays, TEM, CD, and ligand profiling each reported on different aspects of fibril structure. A combination of these techniques provided the most accurate assessment of fibril morphology.

This approach first revealed that the morphology of αSyn fibrils evolved during aggregation. ThX and CR gave a distinct kinetic profile to AAR, TPHN-1, and IND, indicating that different binding sites formed at different rates. Ligand profiling also revealed that binding sites were changing even when the total fluorescence of a specific reporting ligand was constant. Different ligands are therefore sensitive to different polymorphs, and differences in the binding sites between polymorphs can exist even when the overall fluorescence intensity is the same.

Once formed, the morphology of αSyn fibrils continued to change under common storage conditions. Substantial structural changes were observed after storage of αSyn fibrils at 4 °C, and after storage at 37 °C only non-fibrillar aggregates were observed. Storage at -79 °C produced the least significant changes in morphology, and fibrils were relatively stable through repeated freeze-thaw cycles. These changes indicate that fibril morphology remains in equilibrium with the local chemical environment under common storage conditions. The effectiveness of a given storage condition is likely dependent on the fibril morphology present and the subsequent kinetic accessibility of other polymorphs.

Fibril morphology was also shown to be sensitive to the presence of exogenous ligands. Typically aggregation is monitored in real-time using the fluorescence of ThT, an analogue of ThX. The addition of ThX to the aggregation reaction was shown to affect aggregation kinetics and produced a more stable fibril morphology. When ThT is included in the aggregation reaction, a common observation in the literature is that ThT fluorescence can “overshoot” during aggregation, reaching an initial maximum before decreasing slightly.^63–66^ This decrease in fluorescence may arise from the structure of αSyn fibrils continuing to change, affecting ThT binding sites.

The addition of ThX and other ligands to preformed fibrils also produced changes in fibril morphology over time. This result is consistent with αSyn fibrils remaining in equilibrium with their environment. This result also raises the possibility of using ligands to guide the formation of specific morphologies, or to convert pathological morphologies to less pathological morphologies. Polymorphism is a well-established feature of protein fibrils and has been extensively studied in the context of crystallography. During aggregation or crystallisation the initial polymorph that nucleates is the most kinetically accessible, but not necessarily the most thermodynamically stable.^67^ These results show that the αSyn fibrils studied appear to transition through a sequence of transient, intermediate morphologies. These transitions occur not only during initial aggregation, but also after complete aggregation has occurred, and did not terminate at a final stable polymorph. Changes in fibril morphology were observed even after over a year in storage at -79 °C. In contrast, *in vivo* fibrils observed in neurodegenerative conditions appear to achieve a final, disease-specific morphology.^41^ These pathological morphologies may be the endpoint of a prolonged series of polymorphs that requires long timescales, or specific pathological conditions, to achieve.

The molecular mechanism producing changes in fibril morphology is unclear. Two possible pathways are that fibrils disaggregate and reaggregate into different morphologies, or structural changes arise on fibrils and propagate down their length. Each new set of polymorphs can then affect subsequent transitions in morphology by secondary nucleation or changing the chemical environment in solution. The energy landscape governing polymorph formation is therefore dynamic and evolves in response to the emergence and disappearance of different morphologies. A feedback loop is established where the presence of one polymorph alters both the relative stability of accessible polymorphs, and the kinetic pathway between them. This energy landscape is also affected by the presence of exogenous ligands such as ThX.

Together, these results suggest several considerations should be made when working with αSyn fibrils.

First, the inclusion of fluorescent dyes to monitor aggregation can influence the aggregation process. The morphology of fibrils formed in the presence and absence of exogeneous ligands or molecules cannot be assumed to be the same.

Second, using a single fluorescent ligand to monitor aggregation may fail to detect certain fibril morphologies or types of aggregates. A combination of fluorescent dyes or other analytical methods may provide a more comprehensive view of the aggregation pathway and products. However, using a single ligand may be useful for monitoring a subset of aggregated species in an otherwise more complex system.

Third, different analytical techniques cannot always distinguish polymorphs. A combination of methods should therefore be used to accurately characterise fibril morphology. Ideally, an analytical method should be used that is relevant to the intended application of the fibrils.

Fourth, the choice of αSyn aggregation time and storage conditions will affect the fibril polymorph used. For a given application, consistent aggregation and storage methods should be combined with thorough characterisation. Storage conditions for αSyn fibrils should be carefully chosen, and the effect on fibril morphology determined.

## METHODS

### Chemistry

All solvents and chemicals were obtained from commercial sources and used without further purification unless otherwise stated. Reactions were monitored by TLC or LCMS. TLC analyses were performed on Merck TLC Silica gel 60 F254 glass plates (0.2 mm). LCMS analyses of samples were performed using a Waters Acquity H-class UPLC coupled with a single quadrupole Waters SQD2. An Acquity UPLC CSH C18 Column, 130Å, 1.7 μm, 2.1 mm x 50 mm was used as the UPLC column.

Purification of compounds by silica column chromatography were performed using an automated system (Combiflash® Rf+) with prepackaged silica cartridges (25 μm PuriFlash® columns). ^1^H and ^13^C NMR spectra were recorded using a Bruker 600 MHz Avance 600 BBI spectrometer or a 400 MHz Avance III HD Smart Probe spectrometer at 298.0 ± 0.1 K. Residual solvent peaks were used as an internal standard for calibration. All chemical shifts are quoted in ppm on the δ scale and the coupling constants are expressed in Hz. Signal splitting patterns are described as a singlet (s), broad singlet (br s), doublet (d), triplet (t), quartet (q), or multiplet (m).

### Small Molecule Synthesis

Compounds **BTA**, **OXI**, and **S5H** were prepared as previously reported.^44^

*2-((1-(4-iodobenzyl)-1H-indol-3-yl)methylene)malononitrile (**XIA**).* To a suspension of **S1** (108 mg, 0.30 mmol, 1.0 equiv.) in methanol was added malonitrile (90 µL, 1.62 mmol, 5.4 equiv.) followed by a saturated solution of K2CO3 in methanol (50 µL), yielding a yellow solution. The reaction mixture was stirred for 15 h then cooled on ice and filtered. The filtrate was washed with ice-cold methanol to afford **XIA** as a light yellow solid (104 mg, 0.25 mmol, 83%). **Yield:** 104 mg (0.25 mmol, 83%). ^1^H NMR (400 MHz, DMSO-*d6*), δ (ppm): 8.71 (d, *J* = 9.5 Hz, 2H), 8.11 – 8.04 (m, 1H), 7.71 (dt, *J* = 8.3, 1.8 Hz, 2H), 7.59 (ddd, *J* = 5.0, 3.6, 2.7 Hz, 1H), 7.35 – 7.28 (m, 2H), 7.10 (dt, *J* = 8.4, 1.9 Hz, 2H), 5.65 (s, 2H). ^13^C NMR (101 MHz, DMSO-*d6*), δ (ppm): 151.9, 137.6, 136.1, 136.0, 135.4, 129.6, 127.5, 124.2, 123.0, 119.4, 115.8, 115.6, 111.9, 110.5, 94.1, 70.2, 49.7. HRMS (ESI+): 409.0053 m/z: Calculated for C19H12IN3^+^ = 409.0070 [M]^+^. 410.0162 m/z: Calculated for C19H13IN3^+^ = 410.0149 [M+H]^+^. 431.9983 m/z: Calculated for C19H12IN3Na^+^ = 431.9968 [M+Na]^+^. IR (ATR, cm^-1^): 2225, 1592, 1578, 1515, 1470, 1401, 1366, 1190, 1007, 835. *(2E,4E)-2-cyano-5-(4-(dimethylamino)phenyl)penta-2,4-dienoic acid (**AAR**)*. A mixture of the methyl ester **S3** (1.00 g, 3.9 mmol) and LiOH (500 mg, 21 mmol) in THF (32 mL) and water (8 mL) was stirred for 16 h. The reaction mixture was acidified with 1 M HCl, then extracted with CH2Cl2 (3 x 50 mL). The combined organic extracts were evaporated under reduced pressure to afford the hydrochloride salt of **AAR** as a crimson solid (1.15 g, 3.5 mmol, 70%). Yield: 1.15 g (3.50 mmol, 70%); ^1^H NMR (400 MHz, DMSO-*d6*), δ (ppm): 8.00 (d, *J* = 11.8 Hz, 1H), 7.58 – 7.48 (m, 3H), 6.92 (dd, *J* = 14.9, 11.9 Hz, 1H), 6.78 (d, *J* = 8.6 Hz, 2H), 3.03 (s, 6H). ^13^C NMR (101 MHz, DMSO-*d6*), δ (ppm): 164.0, 156.3, 152.2, 150.8, 130.8, 122.3, 117.2, 115.9, 112.2, 99.0. HRMS (ESI+): 243.1118 m/z: Calculated for C H N O ^+^ = 243.1128 [M+H]^+^. IR (ATR, cm^-1^): 2922, 2851, 2552, 2216, 1666, 1585, 1541, 1483, 1441, 1412, 1358, 1317, 1285, 1270, 1237, 1159, 1089, 1065, 1050, 1032, 996, 976, 945, 915, 870, 810.

*6-methoxy-3-methyl-2-(4-(pyrrolidin-1-yl)phenyl)benzo[d]thiazol-3-ium (**ThX**).* A mixture of **S4** (1.02 g, 3.3 mmol) and methyl iodide (2.5 mL) in nitrophenol (8 mL) were heated to 110 °C under microwave irradiation for 4 h. The reaction was cooled to room temperature and triturated with Et2O. The precipitate was collected by filtration and washed thoroughly with Et2O to afford **ThX** as a yellow-orange solid (1.49 g, 3.3 mmol, 100%). Yield: 1.49 g (3.3 mmol, 100%). ^1^H NMR (400 MHz, DMSO-*d6*), δ (ppm): 8.14 (d, J = 9.3 Hz, 1H), 7.97 (d, J = 2.6 Hz, 1H), 7.79 (dt, J = 9.0, 2.9 Hz, 2H), 7.46 (dd, J = 9.2, 2.6 Hz, 1H), 6.81 (dt, J = 9.1, 3.0 Hz, 2H), 4.21 (s, 3H), 3.91 (s, 3H), 3.44 – 3.38 (m, 4H), 2.05 – 1.98 (m, 4H). ^13^C NMR (101 MHz, DMSO-*d6*), δ (ppm): ^13^C NMR (101 MHz, DMSO) δ 171.3, 158.8, 151.0, 137.0, 132.2, 129.7, 118.1, 117.8, 112.3, 110.6, 106.5, 56.2, 47.6, 40.1, 39.9, 39.7, 39.5, 39.3, 39.1, 38.9, 38.3, 24.9. HRMS (ESI+): 325.1347 m/z: Calculated for C19H21N2OS^+^ = 325.1369 [M]^+^. IR (ATR, cm^-1^): 2967, 1600, 1504, 1483, 1441, 1405, 1350, 1299, 1271, 1234, 1206, 1035, 824.

*(E)-2-((E)-3-(4-hydroxy-3-methoxyphenyl)allylidene)-2,3-dihydro-1H-inden-1-one (**IND**).* To a solution of 4-hydroxy-3-methoxycinnamaldehyde (175 mg, 0.98 mmol, 1.00 equiv.) and 1-indanone (130 mg, 0.98 mmol, 1.00 equiv.) in acetic acid (5.0 mL) was added concentrated HCl (11.7 M, 0.25 mL) dropwise. The resultant dark orange solution was stirred at 110 °C for 16 h, yielding a dark green solution. The reaction mixture was cooled to room temperature then poured into ice water and filtered. The precipitate was washed with ice water then purified using silica column chromatography (PE to 1:1 PE:EtOAc) to afford **IND** as a red solid (105 mg, 0.36 mmol, 37%). Yield: 105 mg (0.36 mmol, 37%). ^1^H NMR (400 MHz, DMSO-*d6*), δ (ppm): 9.50 (s, 1H), 7.74 (d, *J* = 7.6 Hz, 1H), 7.64 (d, *J* = 7.5 Hz, 1H), 7.49 – 7.44 (m, 1H), 7.32 – 7.25 (m, 2H), 7.16 – 7.04 (m, 3H), 6.81 (d, *J* = 8.2 Hz, 1H), 3.95 – 3.90 (m, 2H), 3.85 (s, 3H). ^13^C NMR (101 MHz, DMSO-*d6*), δ (ppm): 192.5, 149.2, 148.4, 148.0, 142.9, 138.8, 134.6, 134.4, 133.7, 128.0, 127.5, 126.6, 123.3, 122.1, 122.0, 115.7, 110.5, 55.7, 30.2. HRMS (ESI+): 293.3527 m/z: Calculated for C H O ^+^ = 293.3415 [M+H]^+^. *(E)-2-(benzo[d]thiazol-2-yl)-3-(4-(dimethylamino)phenyl)acrylonitrile (**TPHN-1**)*. A suspension of 4-(dimethylamino)-benzaldehyde (298 mg, 2.0 mmol, 1.0 equiv.) in methanol (6.0 mL) was heated to 65 °C. To the resultant brown solution was added 2-(benzo[*d*]thiazol-2-yl)acetonitrile (383 mg, 2.2 mmol, 1.1 equiv.) then a saturated solution of K2CO3 in methanol (500 µL). The resultant red suspension was stirred for 1 h. The reaction mixture was added to CH2Cl2 (50 mL) and H2O (50 mL). The aqueous layer was washed with CH2Cl2 (2 x 50 mL) and the combined organic extracts were washed with brine, dried over Na2SO4, then filtered and dried *in vacuo*. The crude residue was then recrystallized from hot methanol to afford **TPHN-1** as a red-orange solid (506 mg, 1.7 mmol, 83%). Yield: 506 mg (1.7 mmol, 83%). ^1^H NMR (400 MHz, chloroform-*d*), δ (ppm): 8.09 (s, 1H), 8.02 (dt, *J* = 8.3, 0.9 Hz, 1H), 8.00 – 7.96 (m, 2H), 7.86 (dt, *J* = 8.0, 0.9 Hz, 1H), 7.48 (ddd, *J* = 8.3, 7.2, 1.2 Hz, 1H), 7.37 (ddd, *J* = 8.3, 7.2, 1.2 Hz, 1H), 6.76 – 6.70 (m, 2H), 3.10 (s, 6H). ^13^C NMR (101 MHz, chloroform-*d*), δ (ppm): 154.0, 153.0, 147.1, 134.8, 133.2, 126.7, 125.2, 123.0, 121.6, 120.3, 118.4, 111.8, 97.8, 40.2. HRMS (ESI+): 306.1086 m/z: Calculated for C18H16N3S = 306.1059 [M+H]^+^. *2-(3,4-dimethylphenoxy)-N-(3-(4-fluorophenyl)isoxazol-5-yl)acetamide (**FER**)*. To a solution of 3,4-dimethylphenol (267 mg, 2.19 mmol, 3.0 equiv.) and Cs2CO3 (951 mg, 2.92 mmol, 4.0 equiv.) in DMF (4 mL) was added **S6** (186 mg, 0.73 mmol, 1.0 equiv.) in DMF (2 mL). The resultant dark brown solution was heated at 90 °C for 15.5 h then diluted in 1 M HCl (50 mL) and washed with EtOAc (3 x 50 mL). The combined organic extracts were then washed with 1 M HCl (2 x 50 mL), brine (50 mL), dried over Na2SO4 and concentrated *in vacuo*. The crude residue was purified by silica column chromatography (PE to 1:2 PE:EtOAc) to afford **FER** as an off-white solid(104 mg, 0.31 mmol, 14%). Yield: 104 mg, 0.31 mmol, 14%). ^1^H NMR (400 MHz, chloroform-*d*), δ (ppm): 7.62 – 7.55 (m, 2H), 7.17 (s, 1H), 7.13 – 7.06 (m, 3H), 6.79 (d, *J* = 2.7 Hz, 1H), 6.71 (dd, *J* = 8.3, 2.8 Hz, 1H), 4.66 (s, 2H), 2.25 (s, 3H), 2.21 (s, 3H). ^13^C NMR (101 MHz, chloroform-*d*), δ (ppm): 164.0, 161.5, 155.0, 138.6, 131.0, 130.9, 125.9, 125.9, 123.9, 121.1, 116.4, 116.3, 116.1, 111.8, 67.7, 20.0, 19.00. ^19^F NMR (376 MHz, chloroform-*d*), δ (ppm): -112.3. HRMS (ESI+): 341.1421 m/z: Calculated for C19H18FN2O3^+^ = 341.1296 [M+H]^+^. 363.1295 m/z: Calculated for C19H17FN2O3^+^Na^+^ = 363.1115 [M+Na]^+^.

### Fingerprinting and Diversity Analysis

Reported αSyn ligands with a dissociation constant (*Kd*) below 1.0 µM were selected from a previously reported database.^41^ Morgan fingerprints were calculated with a radius of 2 using the Python cheminformatics toolkit RDKit.^68^ Dimensionality reduction was performed using principal component analysis (PCA) followed by t-distributed stochastic neighbor embedding (t-SNE) using the Python toolkit scikit-learn to visualize the chemical space.^69^

Tanimoto similarity coefficients were calculated between each of the panel ligands using Morgan fingerprints (radius 2), also using the Python cheminformatics toolkit RDKit.^68^

### Preparation of αSyn Fibrils

#### Preparation of Monomeric αSyn

A solution of recombinant monomeric αSyn in 1xPBS (180 µM), expressed and purified as previously described,^10^ was added to an Amicon Ultra-15 Centrifugal filter (15 kDa MWCO) and centrifuged (15 min, 4000 x g). The retained monomeric αSyn was washed with Tris·HCl (50 mM, pH 7.5) and NaCl (100 mM) by adding 5 mL of buffer and centrifuging (15 min, 4000 x g). This wash step was repeated four times in total, and the retained filtrate was stored at -79 °C until use.

#### Aggregation of αSyn Aggregates

Monomeric αSyn was thawed on ice over 1 h, then transferred to a 1.5 mL Protein LoBind Eppendorf microtubule with a PTFE magnetic stirrer bar (8 mm x 1.5 mm). The solution was diluted to 50 µM αSyn to yield a final buffer concentration of 50 mM Tris with 1.5 mM (0.01%) NaN3 (pH 7.4), and optionally 20 µM ThX. The tube was sealed with parafilm and the solution was stirred at 50 rpm (37 °C) until a total of four weeks had elapsed, with aliquots periodically removed for analysis.

#### Effects of αSyn Storage

Aliquots of αSyn aggregates that had been aggregated for 96 h were taken from the main aggregation reaction solution into a 0.5 mL Protein LoBind Eppendorf microtubule and either stored directly in a fridge at 4 °C, or flash frozen in liquid nitrogen and stored at -79 °C for a further 25 days.

#### Incubation of αSyn with Ligands

Aliquots of αSyn aggregates that had been aggregated for 96 h were taken from the main aggregation reaction solution into a 0.5 mL Protein LoBind Eppendorf microtubule with a PTFE magnetic stirrer bar (8 mm x 1.5 mm). Solutions of ThX, CR, TPHN-1, AAR, or IND in DMSO (0.5 mM) were added to afford a final concentration of 20 µM ligand and 48 µM αSyn. The tube was sealed with parafilm and the solutions were stirred at 50 rpm (37 °C) for a further 25 days.

#### Aggregation of αSyn Aggregates for Long-Term Storage

Monomeric αSyn was thawed in a room temperature water bath, then transferred to a 1.5 mL Protein LoBind Eppendorf microtubule with a PTFE magnetic stirrer bar (8 mm x 1.5 mm). The solution was stirred at 50 rpm (37 °C) for 96 h then aliquots were flash frozen in liquid nitrogen and stored at -79 °C. Aliquots were stored for either 1 year and 3 months, or 1 year and 8 months. Aliquots were thawed in room temperature water when used.

#### Freeze-Thaw Experiments

Aliquots of αSyn fibrils stored at -79 °C were thawed in a room temperature water bath. These aliquots were then flash-frozen in liquid nitrogen. This process was repeated until the desired number of freeze thaw cycles were obtained.

### Fitting of Kinetic Data

Fits to the kinetic data were obtained using AmyloFit.^28^ A secondary nucleation dominated, unseeded kinetic model was chosen, and 50 basin hops were used when fitting. All parameters were fitted except monomer concentration, which was fixed at 50 μM.

### General Procedure for Fluorescence Measurements

Aliquots of αSyn aggregation mixture were diluted to a final concentration of 1.0 µM αSyn with

2.0 µM reporting ligand and optionally 5.0 µM competing ligand in 1xPBS (pH 7.4) in Corning 384-well black non-binding surface plates with clear flat bottoms. The fluorescence of the resultant solution was measured using a BMG Labtech CLARIOstar Plus plate reader.

### Fluorescence Intensity Normalisation

When competing ligands were added to a solution of reporting ligand and αSyn fibril, data were normalised so that a value of 1 corresponded to the fluorescence of reporting ligand in the presence of αSyn fibrils, and a value of 0 corresponded to the fluorescence of reporting ligand in the absence of αSyn fibrils (i.e. complete displacement). This normalisation was performed according to Eq. 1,

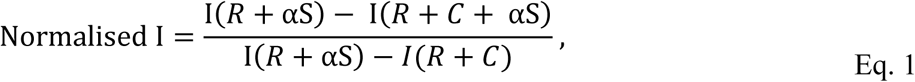

where I(*R* + αS) is the measured fluorescence intensity of the reporting ligand, *R*, in the presence of αSyn fibrils (i.e. the expected fluorescence intensity if *C* does not displace any *R*), I(*R* + *C* + αS) is the measured fluorescence intensity of *R* and the competing ligand, *C*, in the presence of αSyn fibrils, and I(*R* + *C*) is the measured fluorescence intensity of *R* in the presence of *C* (i.e. the expected fluorescence intensity if *C* completely displaces *R*).

The metric ΔNorm is defined as the change in reporting ligand displaced by a given competing ligand between two samples containing different αSyn fibrils, calculated according to Eq. 2,

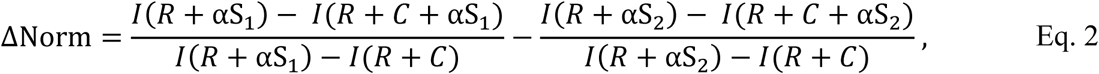

where I(*R* + αSλ) is the measured fluorescence intensity of *R* in the presence of αSyn fibrils prepared via a specific method (αSλ), I(*R* + *C* + αSλ) is the measured fluorescence intensity of *R* in the presence of *C* and αSλ, I(*R* + αSλ) is the measured fluorescence intensity of *R* in the presence of αSλ.

## ASSOCIATED CONTENT

### Supporting Information

Synthesis of intermediates and characterization data of all novel compound, additional analytical procedures, biophysical characterisation of αSyn aggregates, binding assay procedures and data (PDF)

Database of αSyn ligands used in fingerprinting; list of selected panel ligands (CSV)

Code to visualize the chemical space of the αSyn ligand database; code for similarity analysis (IPYNB)

### Corresponding Author

*Timothy S. Chisholm. *

### Funding Sources

T.S.C. thanks the Cambridge Philosophical Society Henslow Fellowship for funding.

## Supporting information

Supporting experimental information

Supporting code and .csv files

## ACKNOWLEDGMENT

T.S.C. thanks the Cambridge Philosophical Society Henslow Fellowship for funding, and Professor Christopher Hunter for useful discussions and providing feedback on manuscript drafts.

T.S.C thanks the EPSRC Underpinning MultiUser Equipment Call (EP/P030467/1).

## ABBREVIATIONS

αSyn, α-synuclein; CD, circular dichroism; CR, Congo red; PD, Parkinson’s disease; LCMS, liquid chromatography–mass spectrometry; MWCO, molecular weight cut-off; NMR, nuclear magnetic resonance; PTFE, polytetrafluoroethylene; TEM, transmission electron microscopy; ThR, thiazine red; ThT, Thioflavin T; ThX, Thioflavin X; TLC, thin layer chromatography; t-SNE, t-distributed stochastic neighbor embedding; UPLC, ultra-performance liquid chromatography.

