## Supporting experimental information for "The Morphology of α-Synuclein Fibrils Changes during Formation, Storage, and upon Exposure to Ligands"

Timothy S. Chisholm [a,*]

[a] Yusuf Hamied Department of Chemistry, University of Cambridge, Lensfield Road, Cambridge CB2 1EW, UK..

### Supporting Information

#### Materials and Instrumentation

All solvents and chemicals were obtained from commercial sources and used without further purification unless otherwise stated. Reactions were monitored by TLC or LCMS. TLC analyses were performed on Merck TLC Silica gel 60 $F_{254}$ glass plates (0.2 mm). LCMS analyses of samples were performed using a Waters Acquity H-class UPLC coupled with a single quadrupole Waters SQD2. An Acquity UPLC CSH C18 Column, 130Å, 1.7 µm, 2.1 mm x 50 mm was used as the UPLC column.

Purification of compounds by silica column chromatography were performed using an automated system (Combiflash® Rf+) with prepackaged silica cartridges (25 µm PuriFlash® columns). $^{1}H$ and $^{13}C$ NMR spectra were recorded using a Bruker 600 MHz Avance 600 BBI spectrometer or a 400 MHz Avance III HD Smart Probe spectrometer at 298.0 ± 0.1 K. Residual solvent peaks were used as an internal standard for calibration. All chemical shifts are quoted in ppm on the δ scale and the coupling constants are expressed in Hz. Signal splitting patterns are described as a singlet (s), broad singlet (br s), doublet (d), triplet (t), quartet (q), or multiplet (m).

UV-vis spectra were collected on an Agilent Cary 60 UV-vis spectrophotometer controlled by Cary WinUV software. Fluorescence spectroscopic data were recorded using an Agilent Cary Eclipse Fluorescence Spectrophotometer controlled by Cary WinUV software, and equipped with a Cary Eclipse Automated Polarizer for anisotropy measurements. Plate reader measurements were performed using a BMG Labtech CLARIOstar Plus plate reader. FT-IR spectra were collected with an ALPHA FT-IR Spectrometer from Bruker. Melting points were recorded with a Mettler Toledo MP90 melting point apparatus.

Protein LoBind (Eppendorf) microtubes were used for preparing and storing all solutions containing protein. All plate reader measurements were performed using Corning 384-well black non-binding surface plates with clear flat bottoms (REF: 3766, LOT: 00422037). Low retention pipette tips were used for all aqueous fluid handling. All buffers were prepared with Milli-Q water and filtered through 0.22 µm filters.

#### Small Molecule Synthesis

**2-(2-(2-((2-(4-(dimethylamino)phenyl)benzo[d]thiazol-6-yl)oxy)ethoxy)ethoxy)ethan-1-ol** (**BTA**)

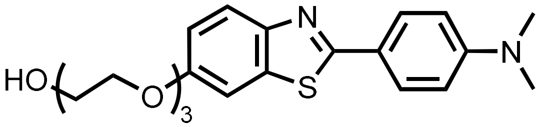

**BTA** was prepared as previously reported.[1]

**(E)-3-((E)-3-(4-nitrophenyl)allylidene)indolin-2-one** (**OXI**)

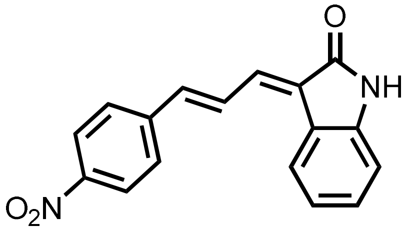

**OXI** was prepared as previously reported.[1,2]

**(E)-5-(4-(benzyloxy)styryl)-3-(thiazol-2-yl)isoxazole / (E)-3-(4-(benzyloxy)styryl)-5-(thiazol-2-yl)isoxazole** (**S5H**)

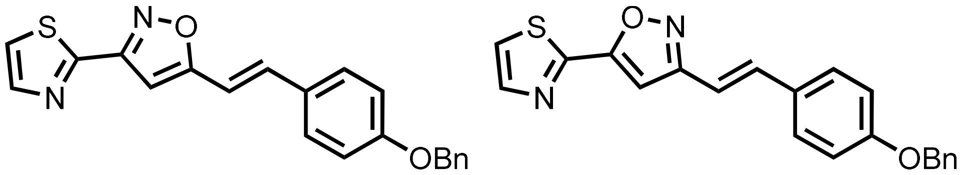

**S5H** was prepared as a mixture of isomers as previously reported.[1,3]

**2-amino-5-methoxybenzenethiol** (**S1**)

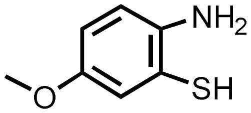

**S1** was prepared as previously reported.[1,2]

**2-(4-bromophenyl)-6-methoxybenzo[d]thiazole** (**S2**)

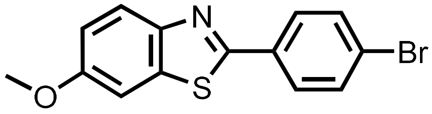

**S2** was prepared was prepared as previously reported.[4]

**6-(2-(2-(2-(2-hydroxyethoxy)ethoxy)ethoxy)ethoxy)-N-(4-(4-(pyridin-2-yl)piperazin-1-yl)phenyl)nicotinamide** (**BF**)

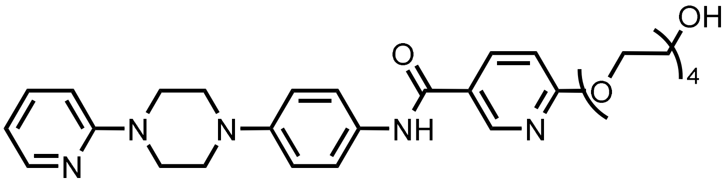

**BF** was prepared was prepared as previously reported.[5]

**Methyl (2E,4E)-2-cyano-5-(4-(dimethylamino)phenyl)penta-2,4-dienoate** (**S3**)

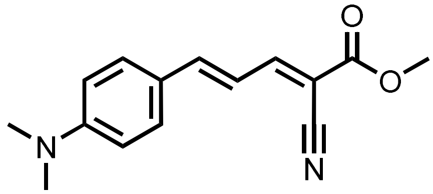

**S3** was prepared was prepared based on a previously reported procedure.[6] To a solution of 4-dimethylaminocinnamaldehyde (876 mg, 5.00 mmol, 1.0 equiv.) in methanol (13.0 mL) was added methyl 2-cyanoacetate (507 μL, 5.75 mmol, 1.15 equiv.) and catalytic piperidine (50 μL). The resultant suspension was stirred for at room temperature for 16 h. The solvent was removed *in vacuo* and the residue was purified using silica column chromatography ($CH_2Cl_2$) to afford **S3** as a crimson solid (1.09 g, 5.26 mmol, 85%).

**Yield:** 1.09 g (5.26 mmol, 85%)

**Aspect:** crimson solid

**$^1$H NMR (400 MHz, chloroform-*d*), δ (ppm):** 7.97 (d, J = 11.7 Hz, 1H), 7.48 (dt, J = 9.1, 2.2 Hz, 2H), 7.20 (d, J = 15.0 Hz, 1H), 7.06 (dd, J = 15.0, 11.7 Hz, 1H), 6.67 (dt, J = 9.0, 2.2 Hz, 2H), 3.86 (s, 3H), 3.07 (s, 6H).

**$^{13}$C NMR (101 MHz, chloroform-*d*), δ (ppm):** 164.0, 157.0, 152.6, 150.7, 131.1, 122.7, 118.3, 115.9, 112.0, 98.8, 52.8, 40.2.

**HRMS (ESI+):** 257.1466 m/z: Calculated for $C_{15}H_{17}N_2O_2^+$ = 257.1284 $[M+H]^+$.

**IR (ATR, cm$^{-1}$):** 2951, 2906, 2214, 1716, 1616, 1592, 1554, 1485, 1435, 1373, 1307, 1289, 1236, 1192, 1160, 1081, 987, 945, 810.

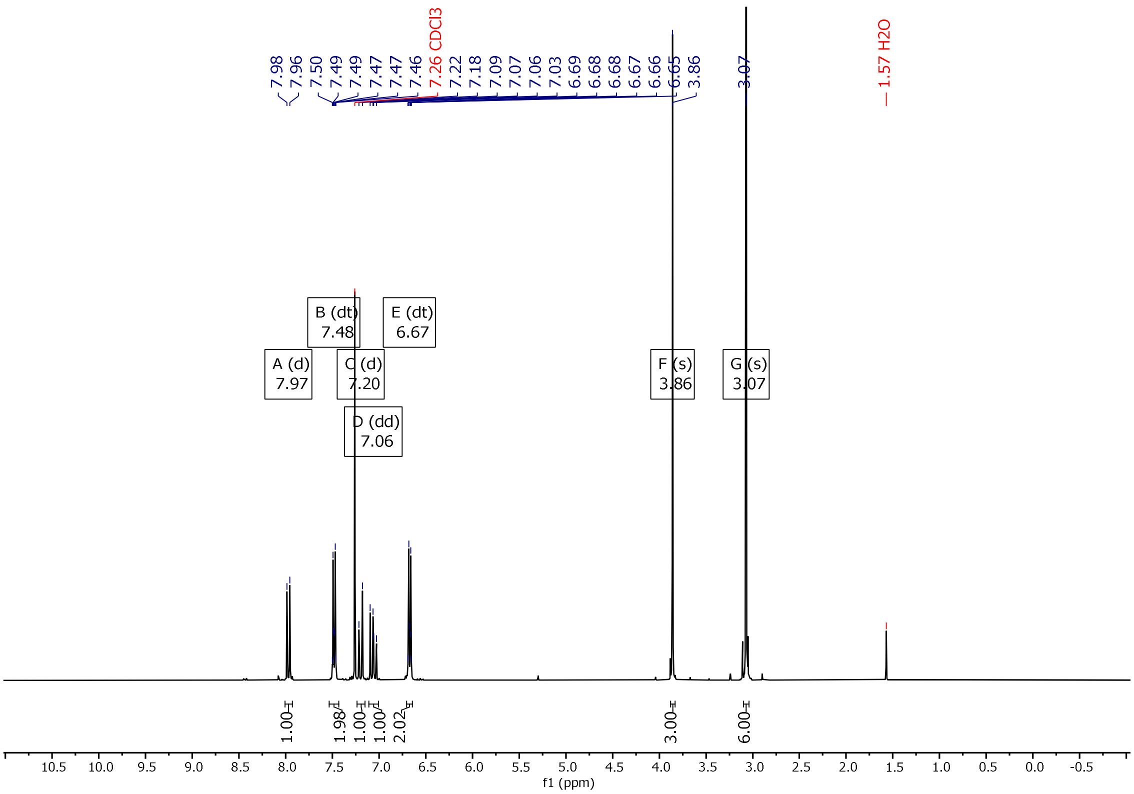

**Figure S1**. $^{1}H$ NMR (400 MHz, $CDCl_3$) spectra of **S3**.

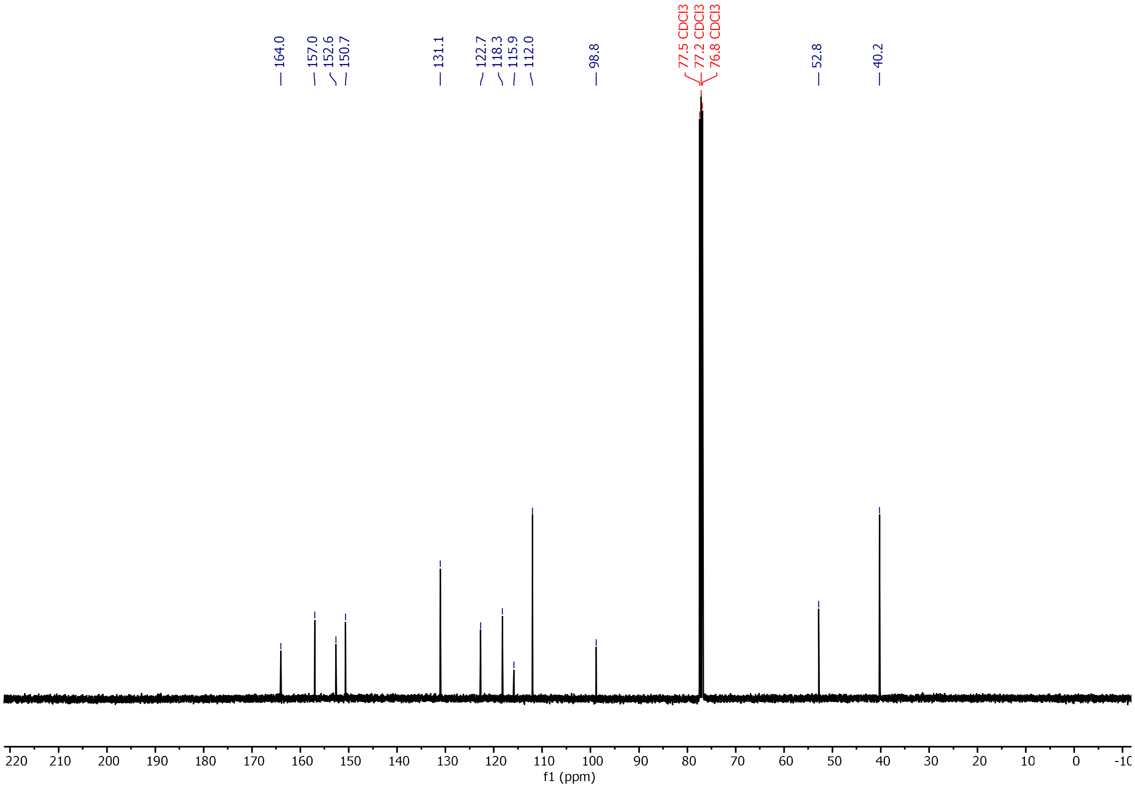

**Figure S2**. $^{13}C$ NMR (101 MHz, $CDCl_3$) spectra of **S3**.

**(2E,4E)-2-cyano-5-(4-(dimethylamino)phenyl)penta-2,4-dienoic acid (AAR)**

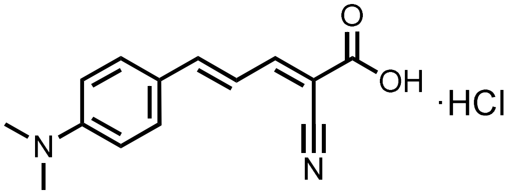

**AAR** was prepared was prepared based on a previously reported procedure.[6]

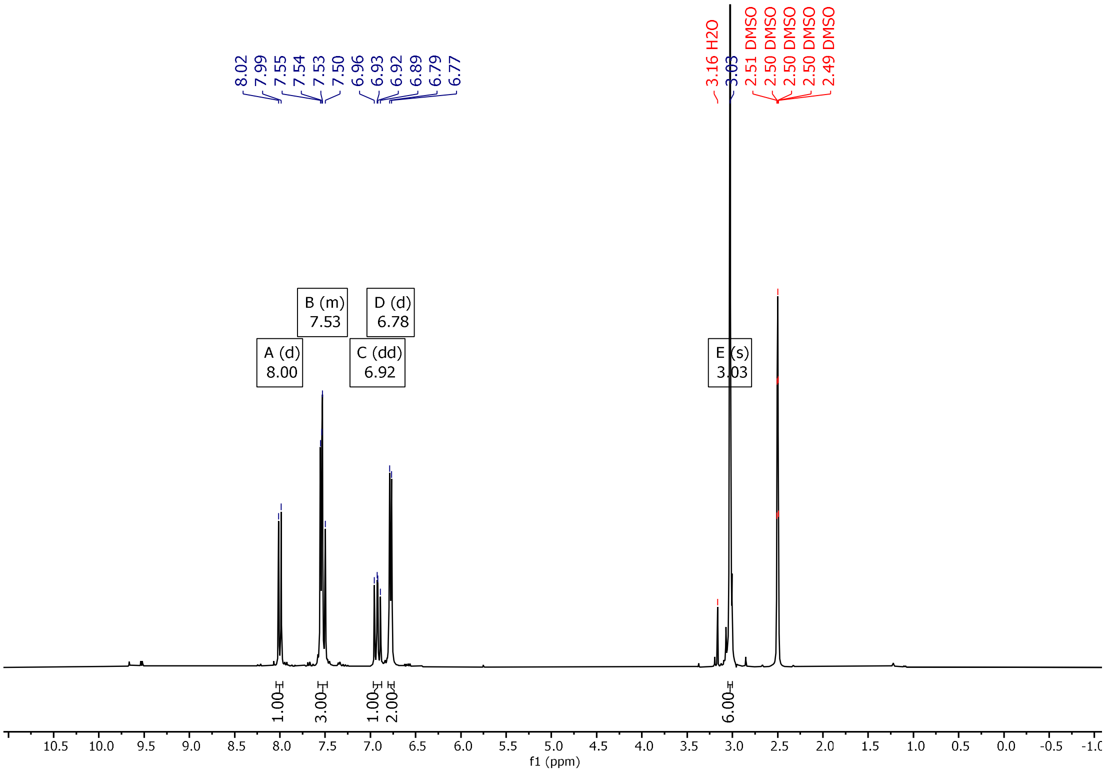

**Figure S3**. $^{1}H$ NMR (400 MHz, DMSO-$d_6$) spectra of **AAR**.

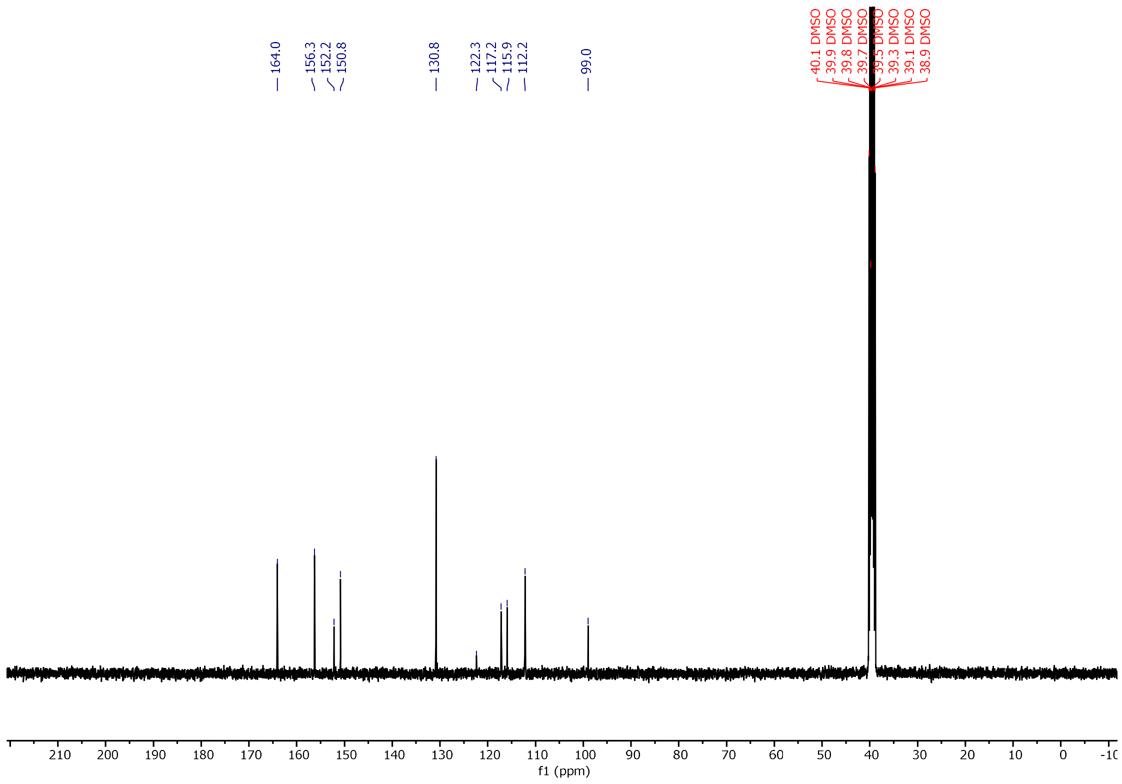

**Figure S4**. $^{13}C$ NMR (101 MHz, DMSO-$d_6$) spectra of **AAR**.

**6-methoxy-2-(4-(pyrrolidin-1-yl)phenyl)benzo[d]thiazole** (**S4**)

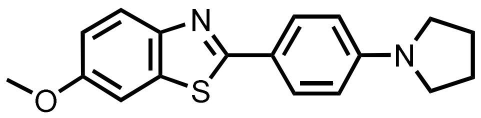

**S4** was prepared was prepared based on a previously reported procedure.[4] To a mixture of **S2** (1.30 g, 4.06 mmol,1.0 equiv.), $Pd_2(dba)_3$ (37.5 mg, 0.041 mmol, 0.01 equiv.), (*R*)-BINAP (76.6 mg, 0.123 mmol, 0.03 equiv.) and NaO*t*-Bu (552 mg, 5.74 mmol, 1.4 equiv.) in toluene (16 mL) was added pyrrolidine (0.47 mL, 4.9 mmol,1.2 equiv.). The resultant brown suspension was heated to 80 °C for 1 h under microwave irradiation. The reaction was then cooled to room temperature, diluted with EtOAc (30 mL), then filtered through celite. The filtrate was concentrated then suspended in MeOH and filtered with celite. The residue was washed with methanol until the filtrate ran clear, then eluted with $CH_2Cl_2$ to afford **S4** as a yellow solid (1.04 g, 3.34 mmol, 81%).

**Yield**: 1.04 g, 3.34 mmol, 81%.

**Aspect**: yellow solid

**$^1$H NMR (400 MHz, chloroform-*d*), δ (ppm):** 7.94 – 7.87 (m, 2H), 7.85 (d, *J* = 8.9 Hz, 1H), 7.33 – 7.29 (m, 1H), 7.26 (s, 1H), 7.03 (dt, *J* = 8.9, 1.8 Hz, 1H), 6.62 – 6.56 (m, 2H), 3.88 (d, *J* = 1.2 Hz, 3H), 3.37 (q, *J* = 4.3 Hz, 4H), 2.04 (q, *J* = 4.9 Hz, 4H).

**$^{13}$C NMR (101 MHz, chloroform-*d*), δ (ppm):** 166.9, 157.1, 149.6, 149.2, 135.9, 128.8, 122.8, 121.1, 114.9, 111.6, 104.5, 55.9, 47.7, 25.6.

**HRMS (ESI+):** 311.1364 m/z: Calculated for $C_{18}H_{19}N_2OS^+$ = 311.1213 $[M+H]^+$.

**IR (ATR, cm$^{-1}$):** 2966, 2855, 1606, 1557, 1509, 1484, 1460, 1421, 1390, 1351, 1321, 1268, 1230, 1205, 1186, 1062, 1025, 818.

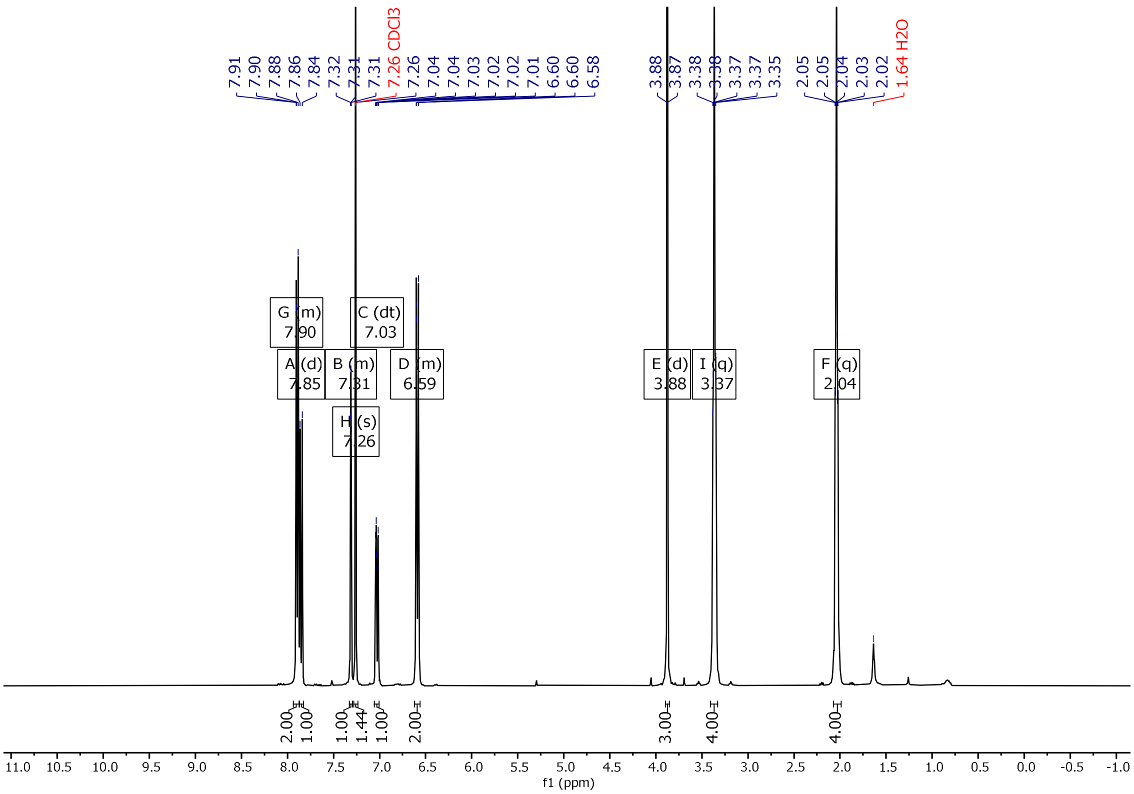

**Figure S5**. $^{1}H$ NMR (400 MHz, $CDCl_3$) spectra of **S4**.

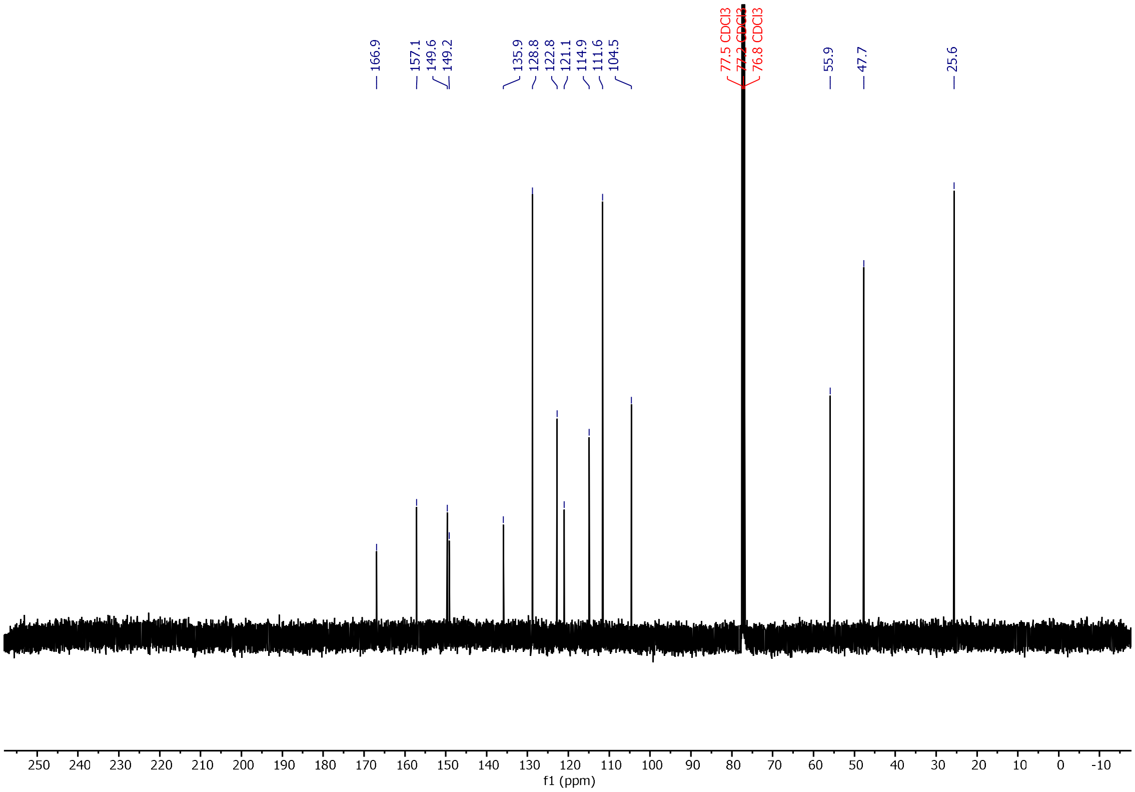

**Figure S6**. $^{13}C$ NMR (101 MHz, $CDCl_3$) spectra of **S4**.

**6-methoxy-3-methyl-2-(4-(pyrrolidin-1-yl)phenyl)benzo[d]thiazol-3-ium** (**ThX**).

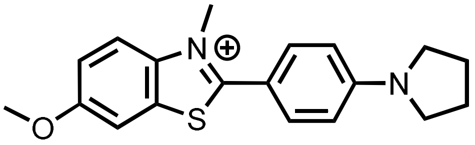

**ThX** was prepared based on a previously reported procedure.[4]

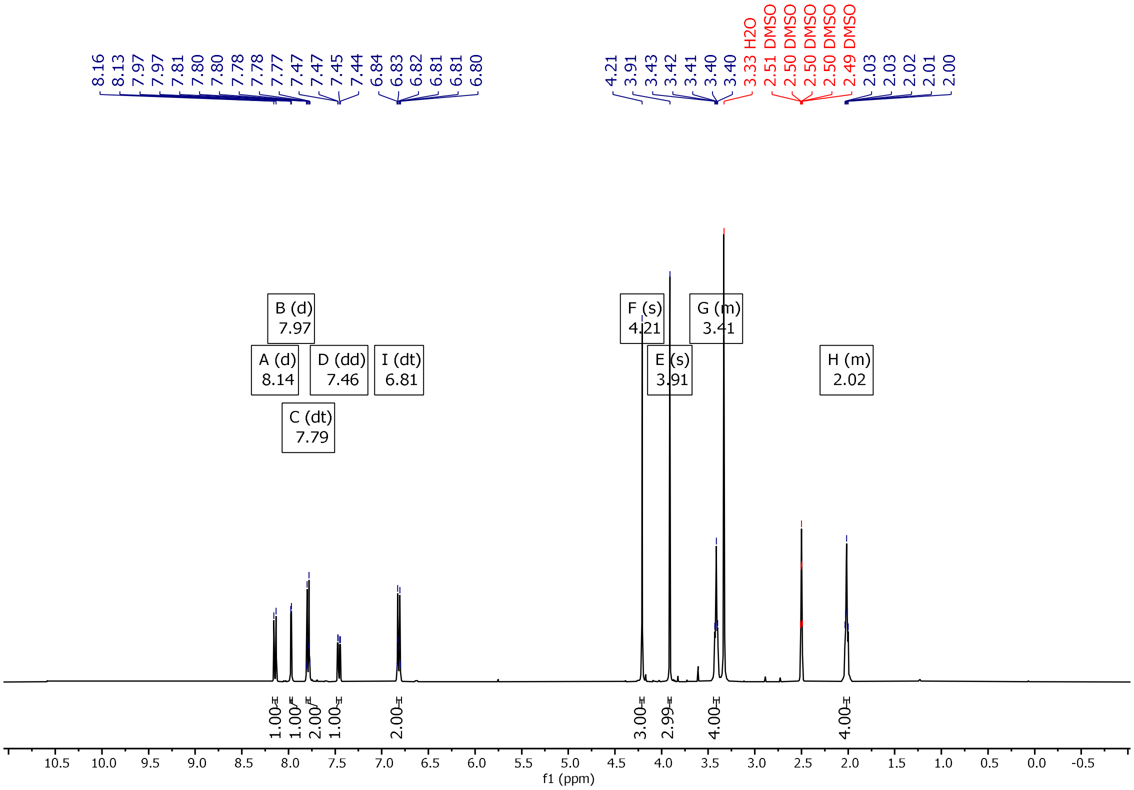

**Figure S7**. $^1$H NMR (400 MHz, DMSO-$d_6$) spectra of **ThX**.

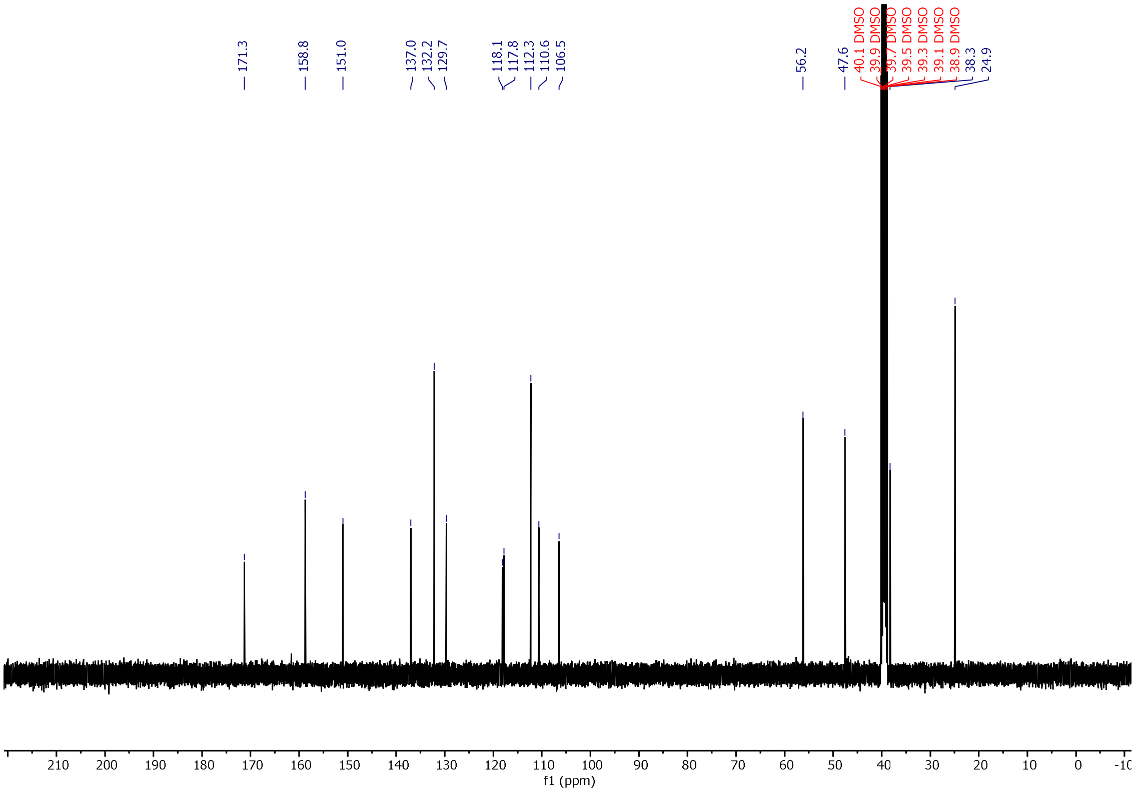

**Figure S8**. $^{13}C$ NMR (101 MHz, DMSO-$d_6$) spectra of **ThX**.

**1-(4-iodobenzyl)-1H-indole-3-carbaldehyde** (**S5**)

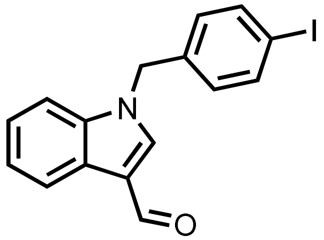

**S5** was prepared was prepared based on a previously reported procedure.[7] To a solution of 1*H*-indole-3-carbaldehyde (146 mg, 1.0 mmol, 1.0 equiv.), $K_2CO_3$ (366 mg, 2.7 mmol, 2.7 equiv.), and KI (80 mg, 0.7 mmol, 0.7 equiv.) in MeCN (5.0 mL) was added 1-(bromomethyl)-4-iodobenzene (388 mg, 1.3 mmol, 1.3 equiv.). The resultant suspension was stirred for 2 h. The precipitate formed was isolated by filtration and added to $CH_2Cl_2$ (50 mL) and $H_2O$ (50 mL). The aqueous layer was washed with $CH_2Cl_2$ (2 x 50 mL) and the combined organic extracts were washed with brine (50 mL), dried over $Na_2SO_4$, then filtered and reduced *in vacuo*. The crude residue was purified using silica column chromatography (PE to 1:1 PE:EtOAc) to afford **S5** as an off-white solid (357 mg, 0.99 mmol, 99%).

**Yield:** 357 mg (0.99 mmol, 99%)

**Aspect:** off-white solid

**$^1$H NMR (400 MHz, DMSO-*d*$_6$), δ (ppm):** 9.94 (s, 1H), 8.46 (s, 1H), 8.14 – 8.09 (m, 1H), 7.71 (dt, *J* = 8.2, 1.9 Hz, 2H), 7.58 – 7.53 (m, 1H), 7.29 – 7.22 (m, 2H), 7.11 (dt, *J* = 8.2, 1.5 Hz, 2H), 5.51 (s, 2H).

**$^{13}$C NMR (101 MHz, DMSO-*d*$_6$), δ (ppm):** 184.7, 141.0, 137.5, 136.9, 136.6, 129.6, 124.8, 123.7, 122.6, 121.1, 117.5, 111.3, 93.9, 49.3.

**HRMS (ESI+)**: 362.0047 m/z: Calculated for $C_{16}H_{13}INO^+$ = 362.0036 $[M+H]^+$.

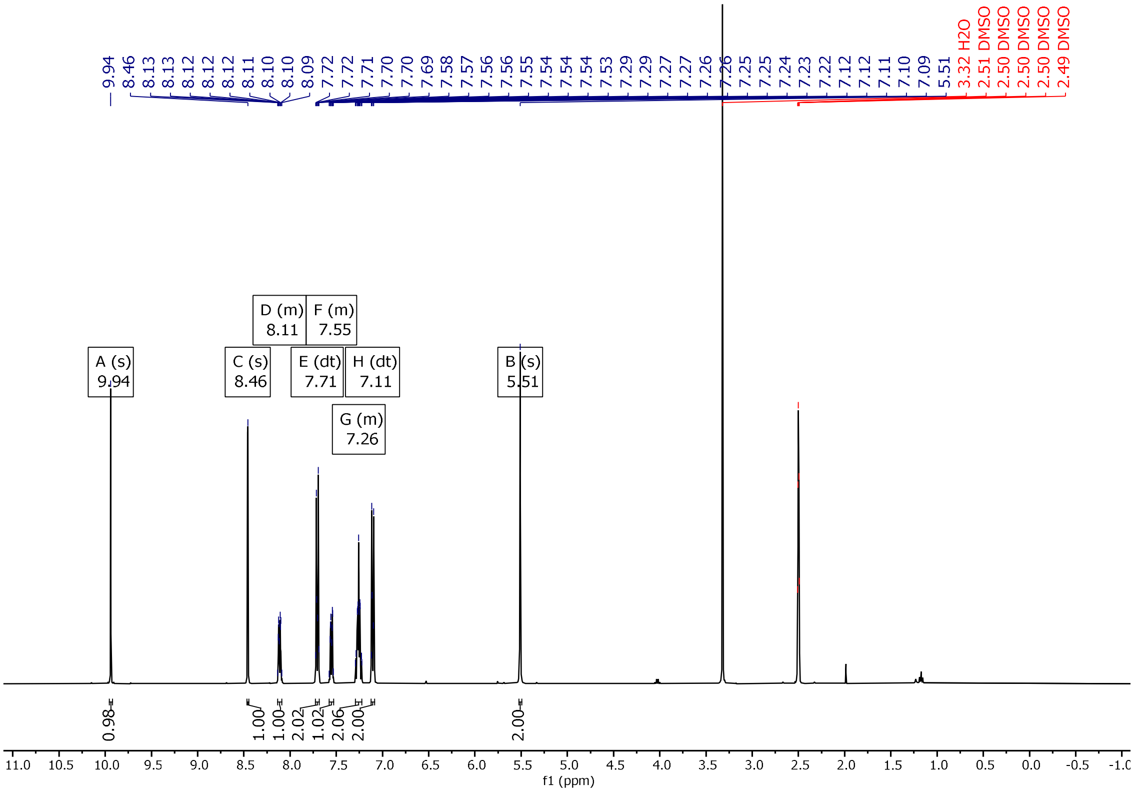

**Figure S9**. $^{1}H$ NMR (400 MHz, DMSO-$d_6$) spectra of **S5**.

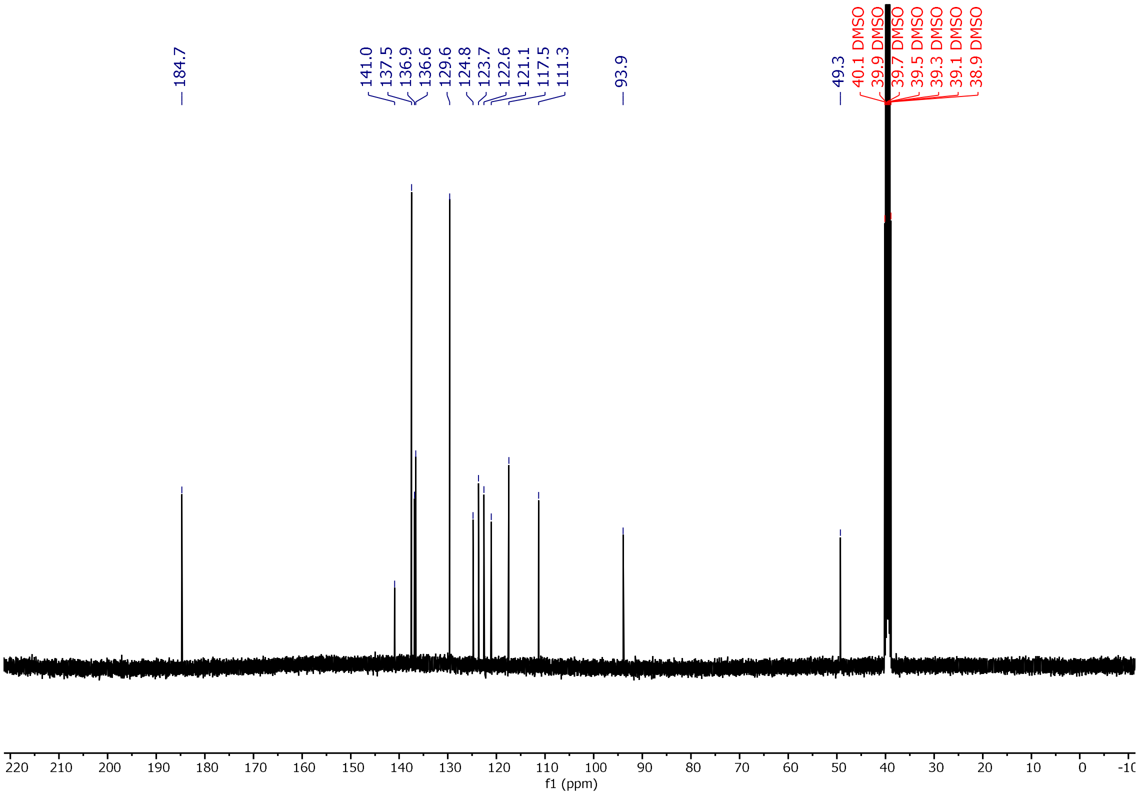

**Figure S10**. $^{13}C$ NMR (101 MHz, DMSO-$d_6$) spectra of **S5**.

**2-((1-(4-iodobenzyl)-1H-indol-3-yl)methylene)malononitrile (XIA)**

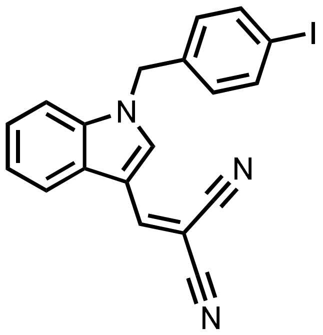

**XIA** was prepared was prepared based on a previously reported procedure.[7]

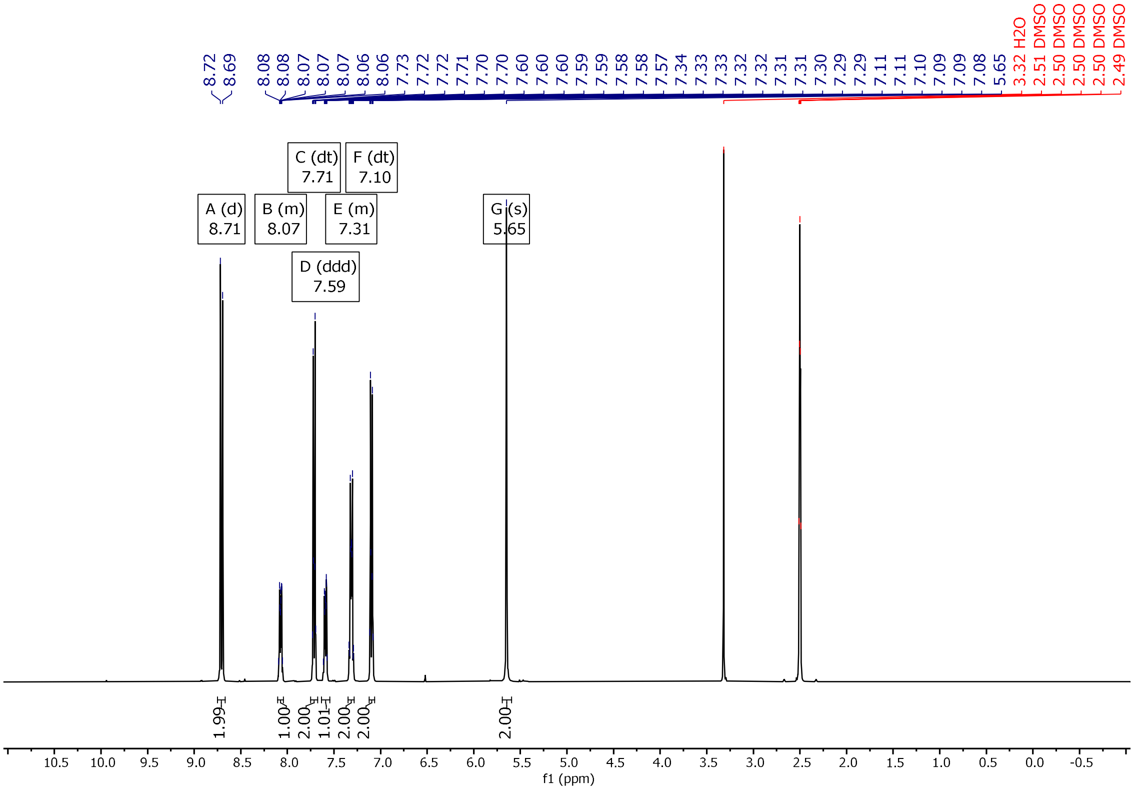

**Figure S11**. $^1$H NMR (400 MHz, DMSO-d$_6$) spectra of **XIA**.

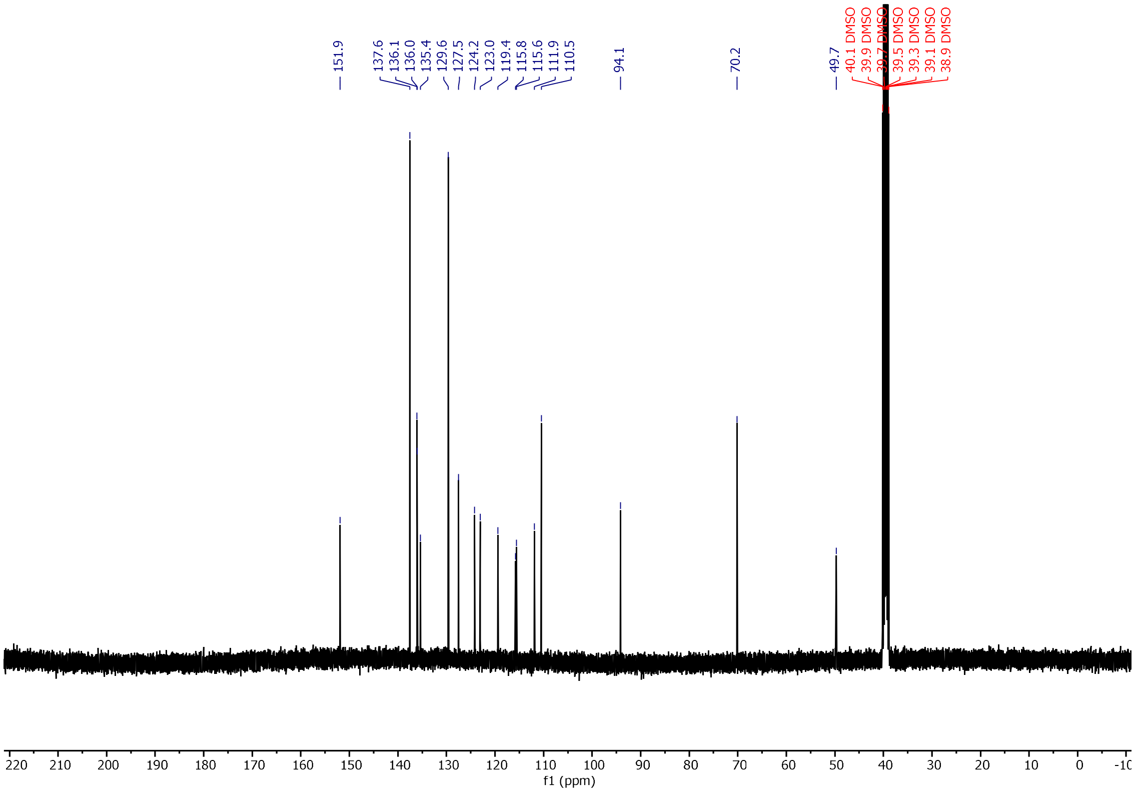

**Figure S12**. [13]C NMR (101 MHz, DMSO-d6) spectra of **XIA**.

**(E)-2-((E)-3-(4-hydroxy-3-methoxyphenyl)allylidene)-2,3-dihydro-1H-inden-1-one (IND)**

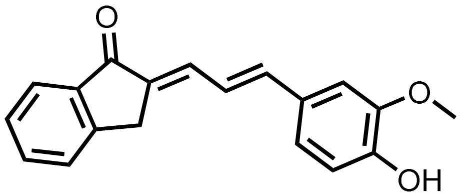

**IND** was reported based on a previously reported procedure.[8]

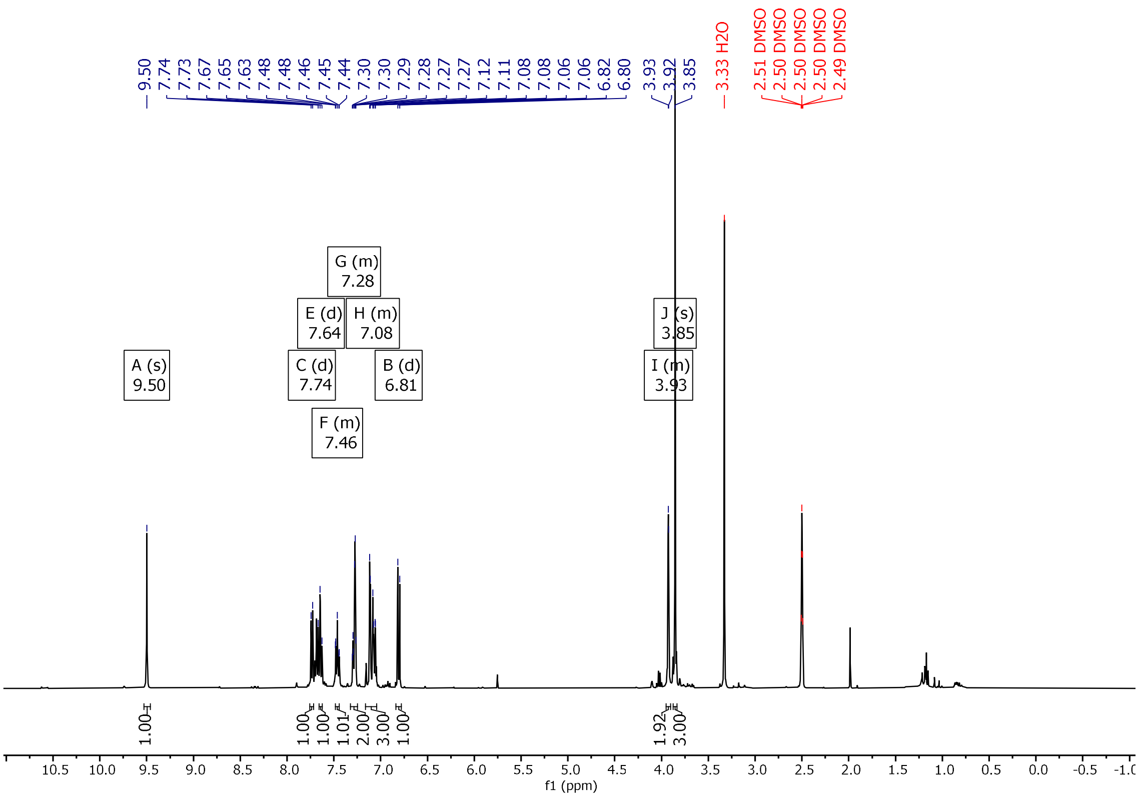

**Figure S13.** $^1$H NMR (400 MHz, DMSO-$d_6$) spectra of **IND**.

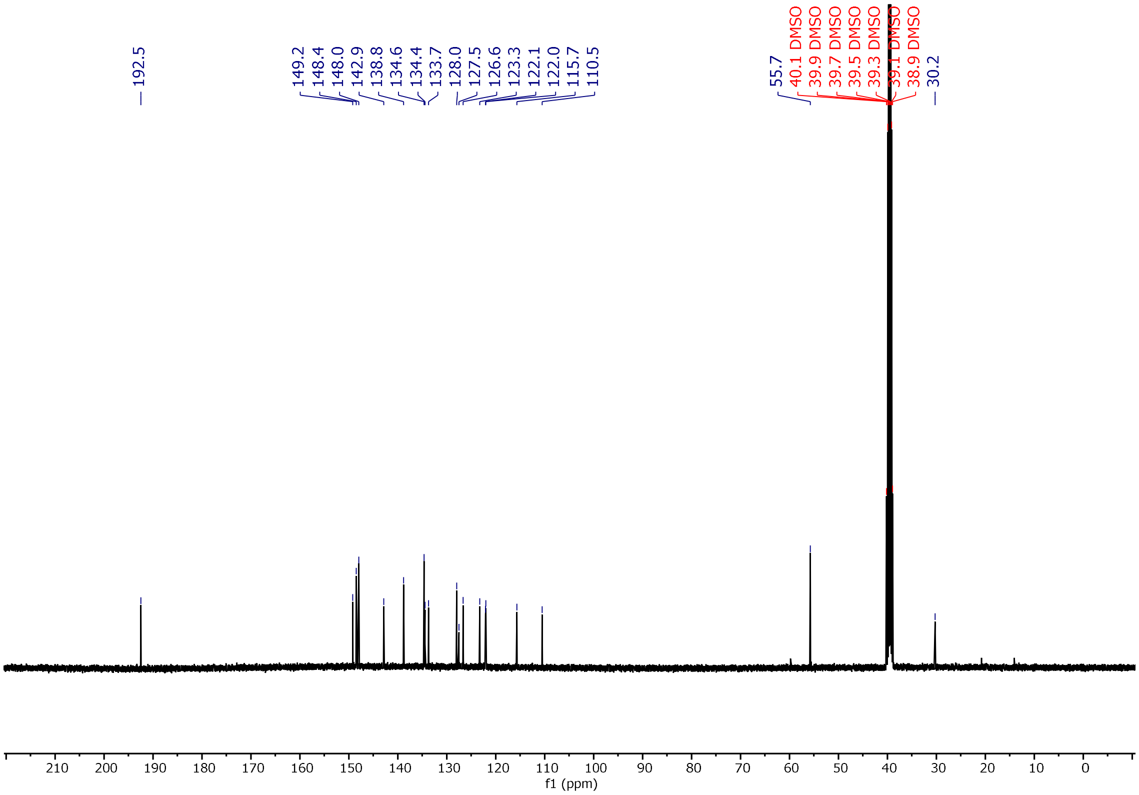

**Figure S14**. $^{13}C$ NMR (101 MHz, DMSO-$d_6$) spectra of **IND**.

**(E)-2-(benzo[d]thiazol-2-yl)-3-(4-(dimethylamino)phenyl)acrylonitrile** (**TPHN-1**)

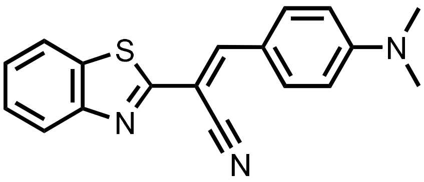

**TPHN-1** was prepared based on a previously reported procedure.[9]

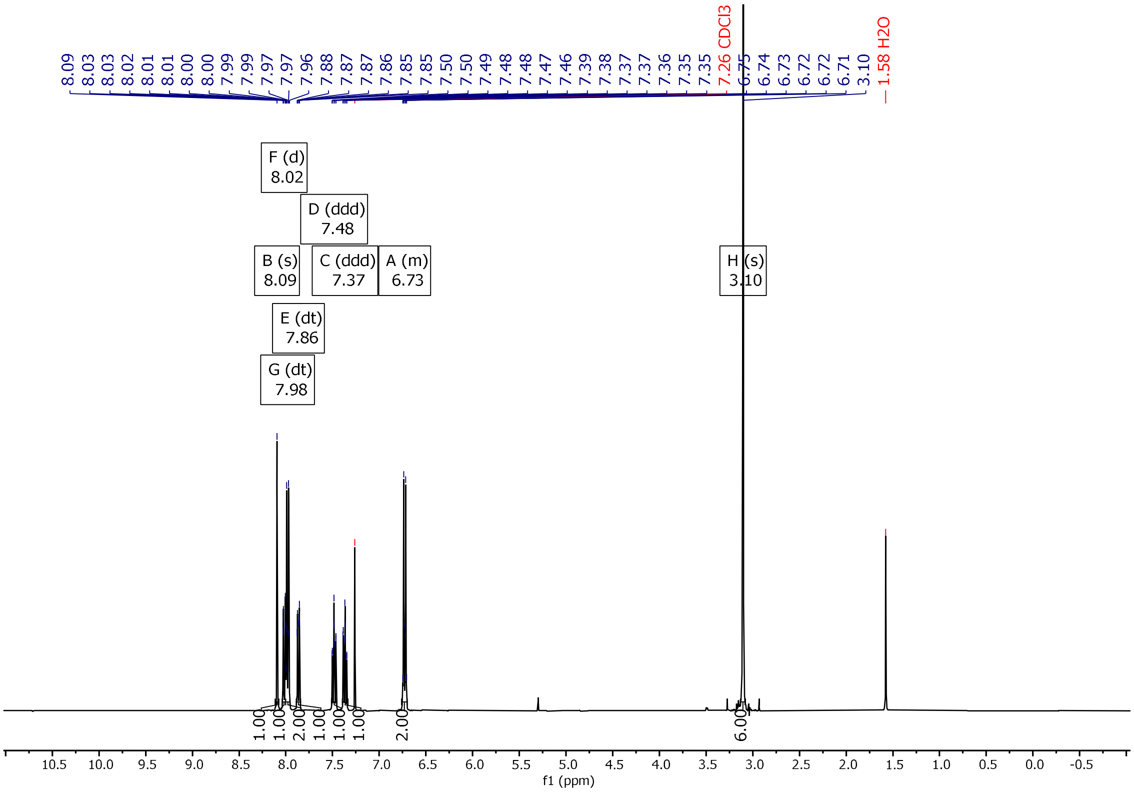

**Figure S15**. $^1H$ NMR (400 MHz, $CDCl_3$) spectra of **TPHN-1**.

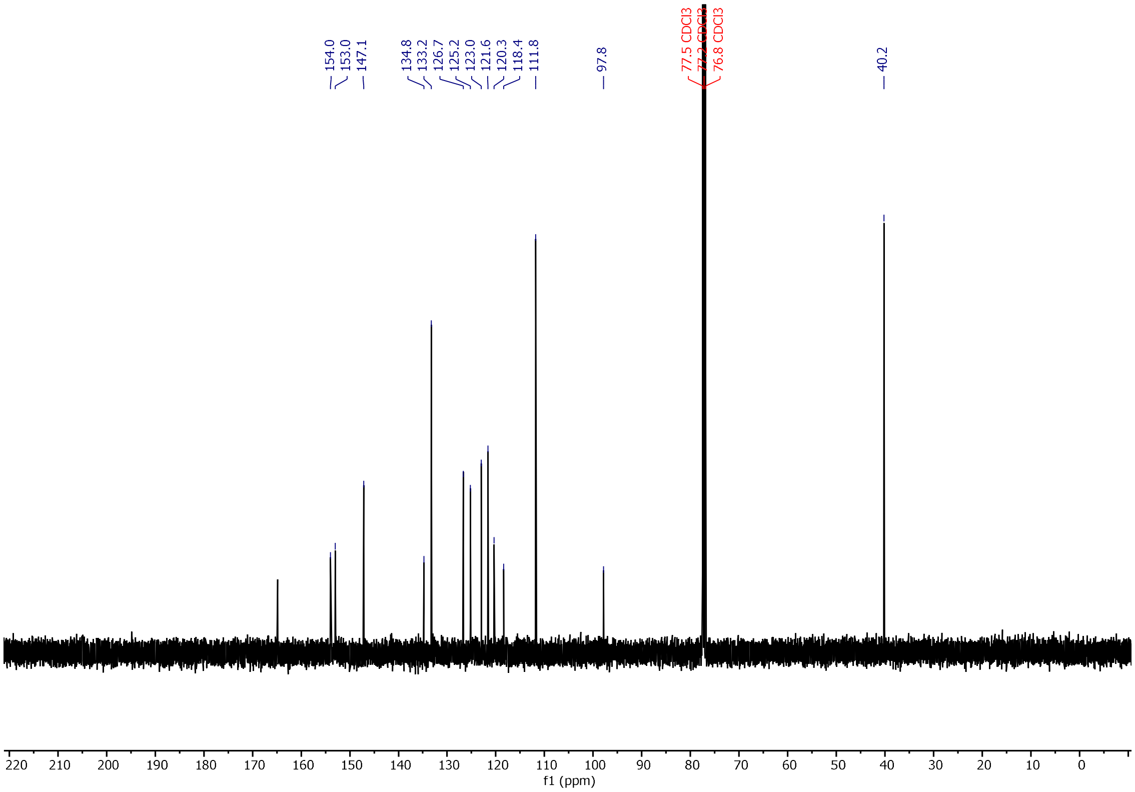

**Figure S16**. [13]C NMR (101 MHz, $CDCl_3$) spectra of **TPHN-1**.

**2-chloro-N-(3-(4-fluorophenyl)isoxazol-5-yl)acetamide** (**S6**)

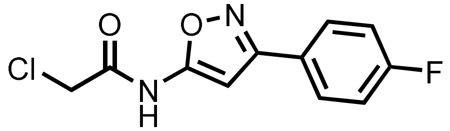

**S6** was prepared based on a previously reported procedure.[10] To a mixture of 3-(4-iodophenyl)isoxazole-5-amine (178 mg, 1.0 mmol, 1.0 equiv.), catalytic DMAP (102 mg), and triethylamine (560 µL, 4.0 mmol, 4.0 equiv.) in DMF (9 mL) on ice was added 2-chloroacetyl chloride (326 µL, 4.1 mmol, 4.1 equiv.) dropwise. The resultant dark brown suspension was stirred on ice for 30 min, then warmed to room temperature and stirred for a further 17 h. The reaction mixture was then added to EtOAc (50 mL) and 1 M HCl (50 mL). The organic layer was isolated, and the aqueous layer washed with EtOAc (2 x 50 mL). The combined organic extracts were then washed with 1 M HCl (2 x 50 mL), brine (50 mL), dried over $Na_2SO_4$, then concentrated *in vacuo*. The crude residue was then purified by silica column chromatography (PE to 1:1 PE:EtOAc) to afford **S6** as an off-white solid (206 mg, 0.81 mmol, 81%).

**Yield:** 206 mg (0.81 mmol, 81%)

**Aspect:** off-white solid

**$^1$H NMR (400 MHz, DMSO-*$d_6$*), δ (ppm):** 11.46 (s, 1H), 8.00 – 7.94 (m, 2H), 7.41 – 7.34 (m, 3H), 4.34 (s, 2H).

**$^{13}$C NMR (101 MHz, DMSO-*$d_6$*), δ (ppm):** 168.1, 165.1, 164.5, 162.0, 158.4, 128.2, 128.1, 123.4, 123.4, 116.5, 116.2, 94.4, 42.9.

**$^{19}$F NMR (376 MHz, DMSO-*$d_6$*) δ (ppm):** -109.5.

**HRMS (ESI+):** 255.0336 m/z: Calculated for $C_{11}H_9ClFN_2O_2^+$ = 255.033 [M+H]$^+$. 277.0153 m/z: Calculated for $C_{11}H_8ClFN_2O_2Na^+$ = 277.0151 [M+Na]$^+$.

**IR (ATR, cm$^{-1}$):** 3233, 3067, 1717, 1629, 1601, 1586, 1558, 1519, 1216, 1160.

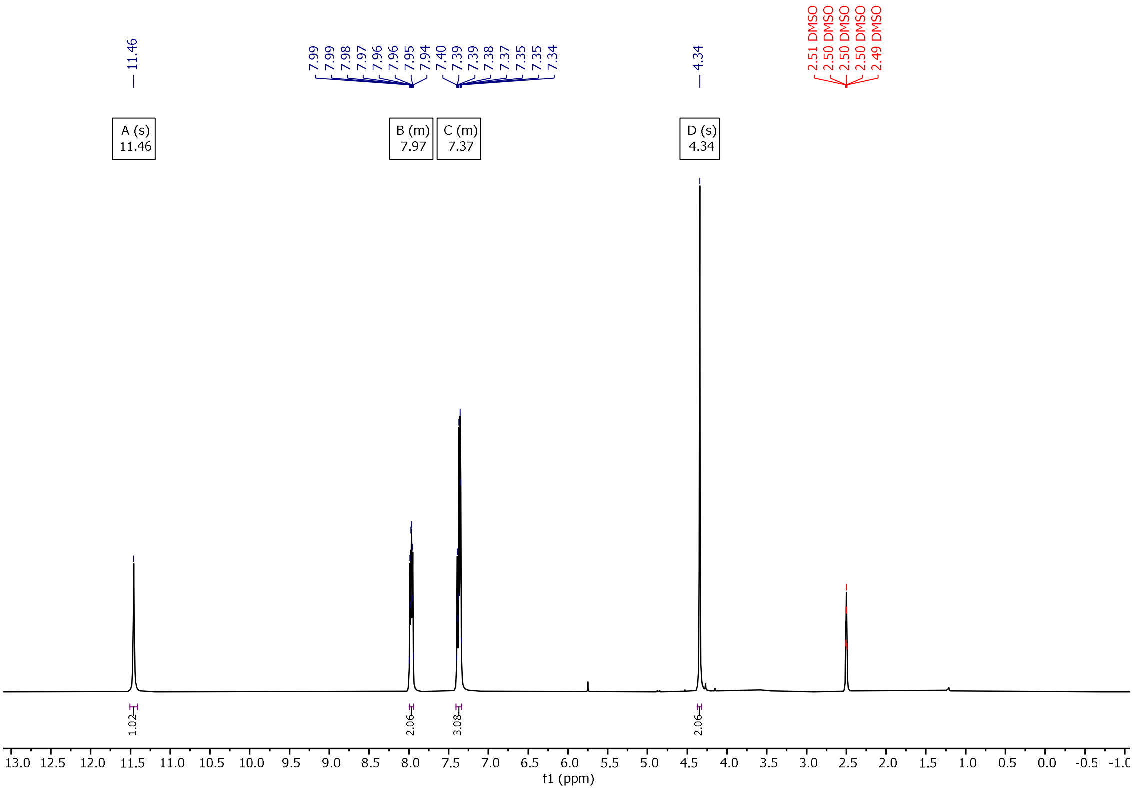

**Figure S17**. $^{1}H$ NMR (400 MHz, DMSO-$d_6$) spectra of **S6**.

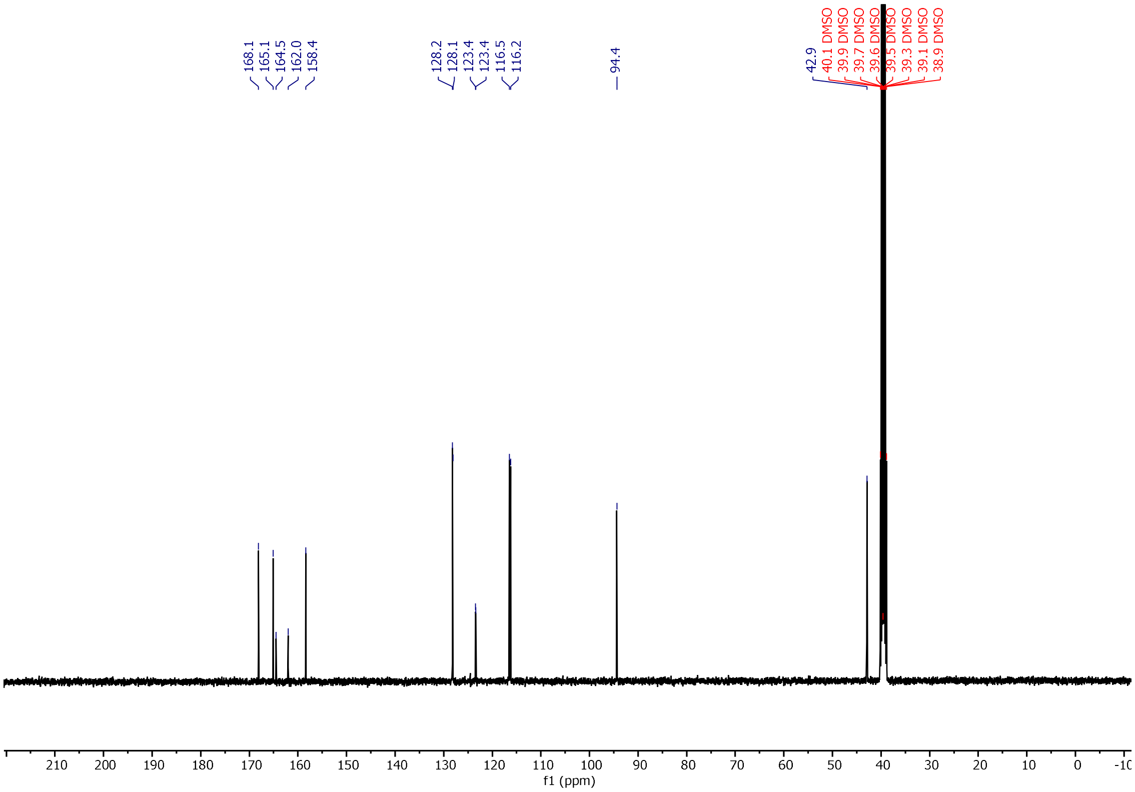

**Figure S18**. $^{13}C$ NMR (101 MHz, DMSO-$d_6$) spectra of **S6**.

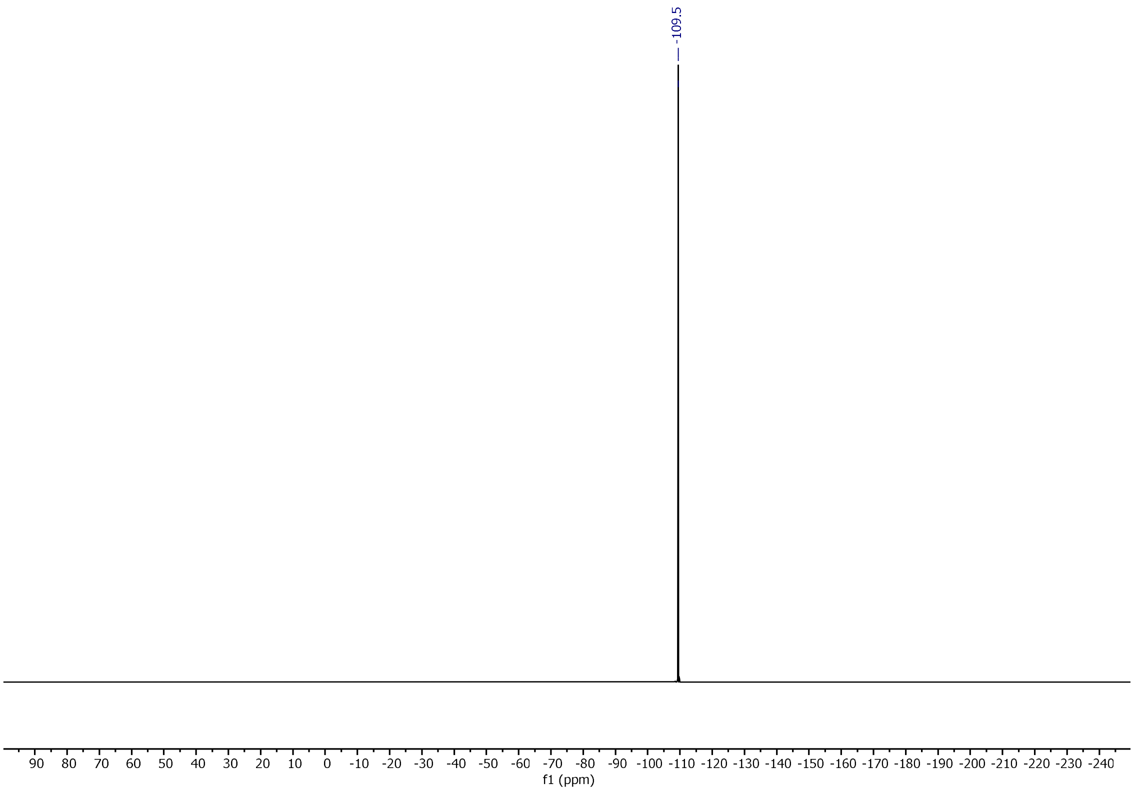

**Figure S19**. $^{19}F$ NMR (376 MHz, DMSO-$d_6$) spectra of **S6**.

**2-(3,4-dimethylphenoxy)-N-(3-(4-fluorophenyl)isoxazol-5-yl)acetamide** (**FER**)

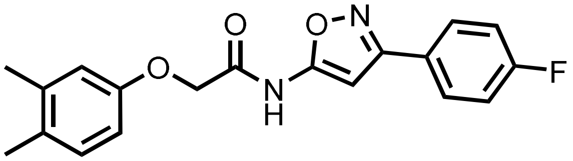

**FER** was prepared based on a previously reported procedure.[10] To a solution of 3,4-dimethylphenol (267 mg, 2.19 mmol, 3.0 equiv.) and $Cs_2CO_3$ (951 mg, 2.92 mmol, 4.0 equiv.) in DMF (4 mL) was added **S6** (186 mg, 0.73 mmol, 1.0 equiv.) in DMF (2 mL). The resultant dark brown solution was heated at 90 °C for 15.5 h then diluted in 1 M HCl (50 mL) and washed with EtOAc (3 x 50 mL). The combined organic extracts were then washed with 1 M HCl (2 x 50 mL), brine (50 mL), dried over $Na_2SO_4$ and concentrated *in vacuo*. The crude residue was purified by silica column chromatography (PE to 1:2 PE:EtOAc) to afford **FER** as an off-white solid(104 mg, 0.31 mmol, 14%).

**Yield:** 104 mg, 0.31 mmol, 14%)

**Aspect:** off-white solid

**$^1$H NMR (400 MHz, chloroform-*d*), δ (ppm):** 7.62 – 7.55 (m, 2H), 7.17 (s, 1H), 7.13 – 7.06 (m, 3H), 6.79 (d, *J* = 2.7 Hz, 1H), 6.71 (dd, *J* = 8.3, 2.8 Hz, 1H), 4.66 (s, 2H), 2.25 (s, 3H), 2.21 (s, 3H).

**$^{13}$C NMR (101 MHz, chloroform-*d*), δ (ppm):** 164.0, 161.5, 155.0, 138.6, 131.0, 130.9, 125.9, 125.9, 123.9, 121.1, 116.4, 116.3, 116.1, 111.8, 67.7, 20.0, 19.00.

**$^{19}$F NMR (376 MHz, chloroform-*d*), δ (ppm):** -112.3.

**HRMS (ESI+):** 341.1421 m/z: Calculated for $C_{19}H_{18}FN_2O_3^+$ = 341.1296 [M+H]$^+$. 363.1295 m/z: Calculated for $C_{19}H_{17}FN_2O_3^+Na^+$ = 363.1115 [M+Na]$^+$.

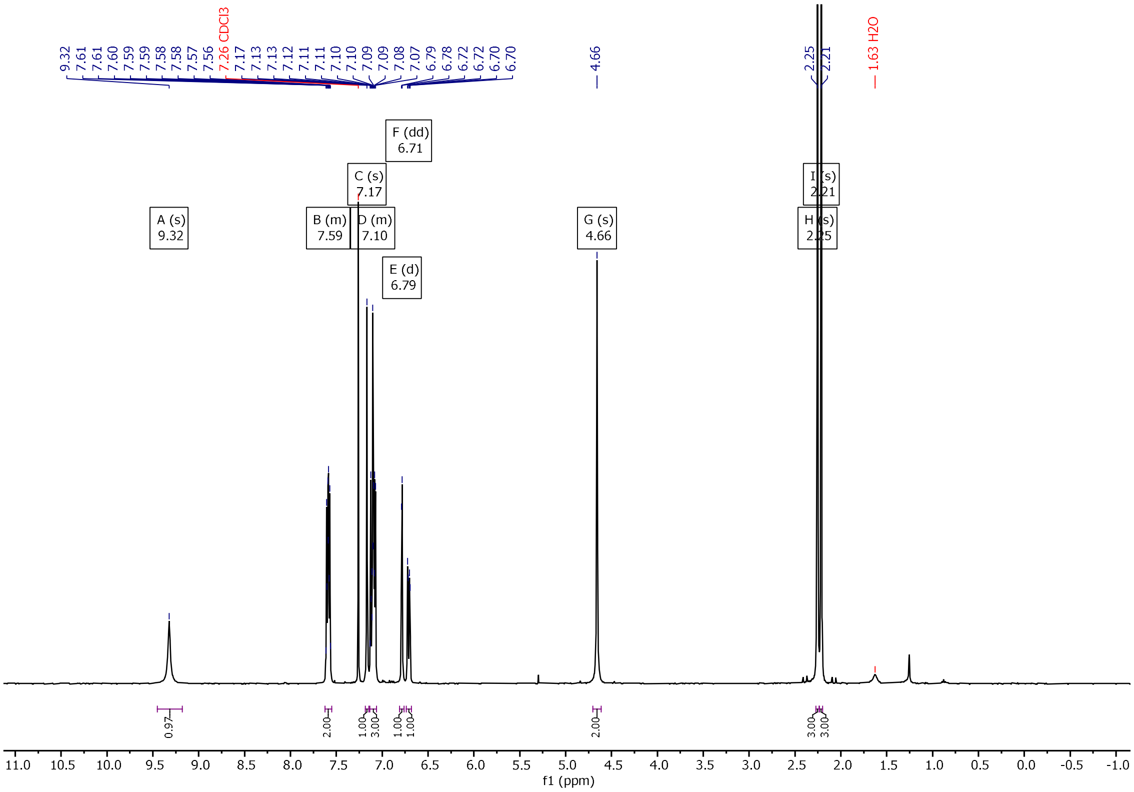

**Figure S20**. $^{1}H$ NMR (400 MHz, $CDCl_3$) spectra of **FER**.

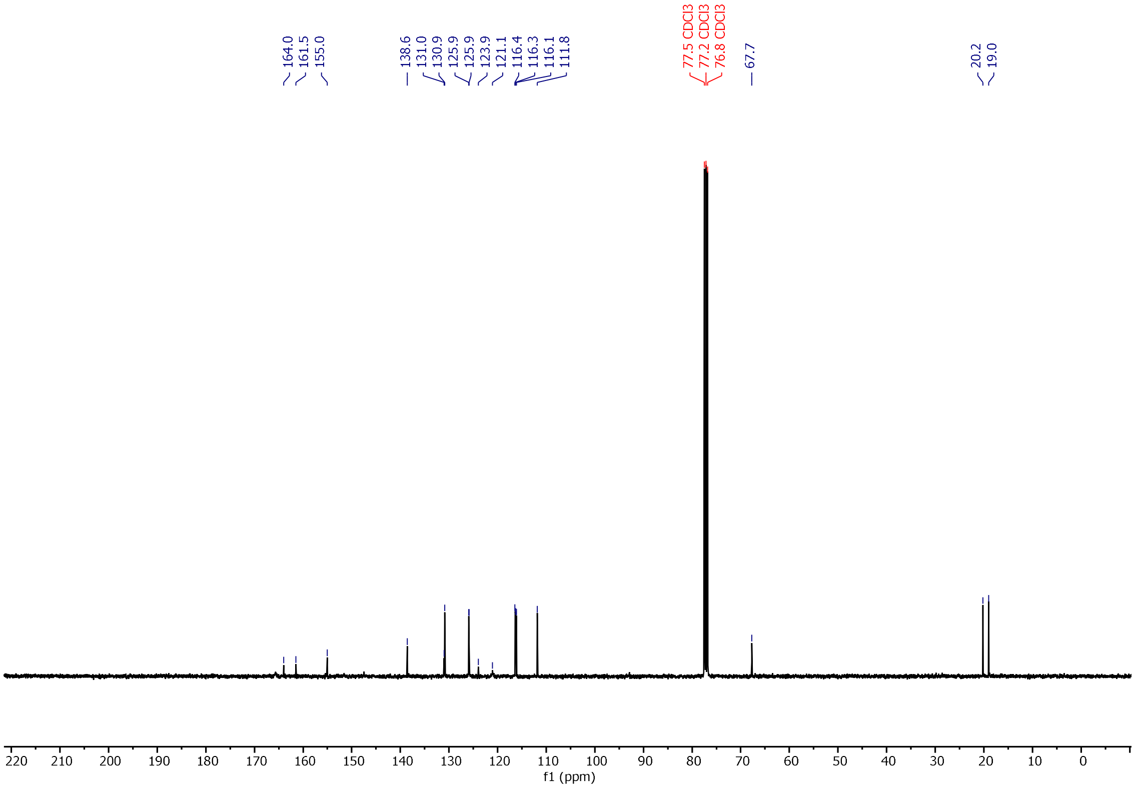

**Figure S21**. $^{13}C$ NMR (101 MHz, $CDCl_3$) spectra of **FER**.

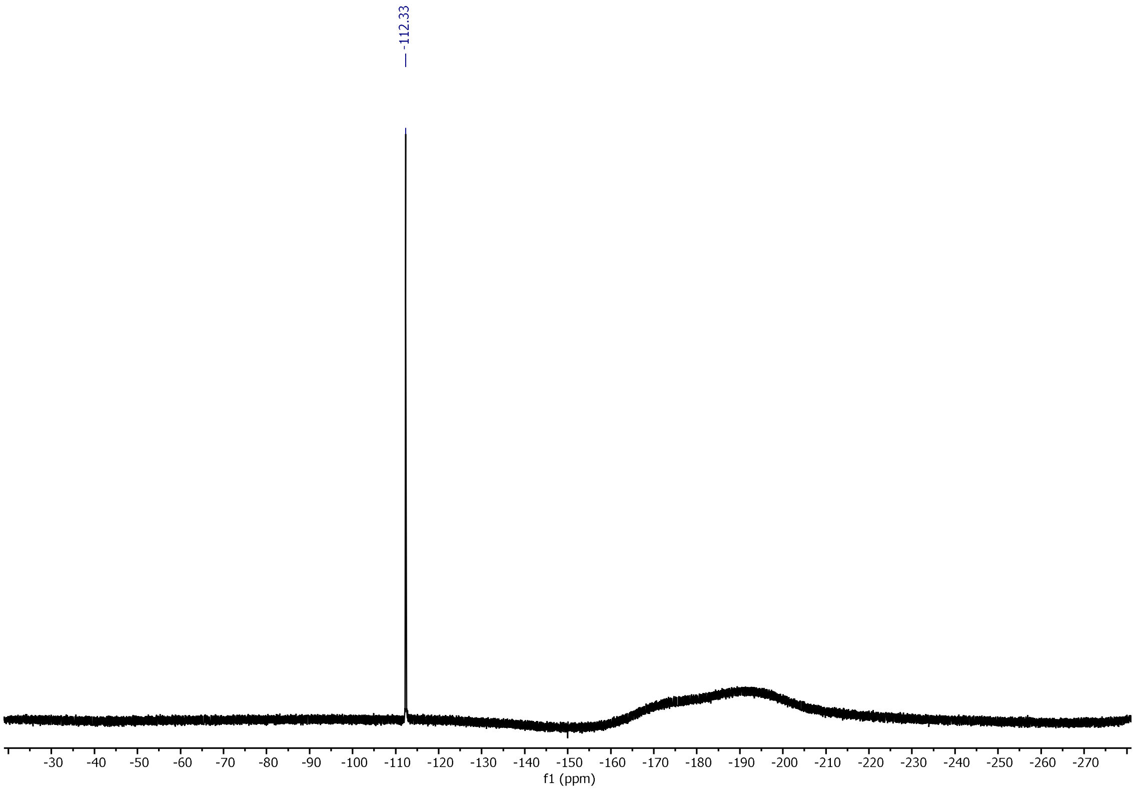

**Figure S22**. $^{19}F$ NMR (376 MHz, DMSO-$d_6$) spectra of **FER**.

#### Fingerprinting and Diversity Analysis

The associated .csv files ("Ligand Database.csv", "Panel Ligands.csv") and iPython notebooks ("Chemical space.ipynb", "Tanimoto Heatmap.ipynb") provide the data and code to replicate these calculations.

### UV-Visible Characterisation

Stock solutions of ligand in DMSO (10 mM) were diluted into ethanol to obtain a 50 μM solution and placed into a quartz fluorescence cuvette (Hellma Analytics) with a 1 cm pathlength. UV-visible spectra were obtained with an Agilent Cary 60 UV-vis spectrophotometer controlled by Cary WinUV software using a scan rate of 600 nm/min, a data interval of 1.0 nm and an averaging time of 0.10 at 25°C.

The UV-visible characterisation of **OXI**, **ThR**, **BTA**, and **S5H** has been previously reported.[1]

Stock solutions of ligand in DMSO (10 mM) were diluted into ethanol to obtain the desired concentration and placed into a quartz fluorescence cuvette (Hellma Analytics) with a 1 cm pathlength. UV-visible spectra were obtained with an Agilent Cary 60 UV-vis spectrophotometer controlled by Cary WinUV software using a scan rate of 600 nm/min, a data interval of 1.0 nm and an averaging time of 0.10 at 25°C.

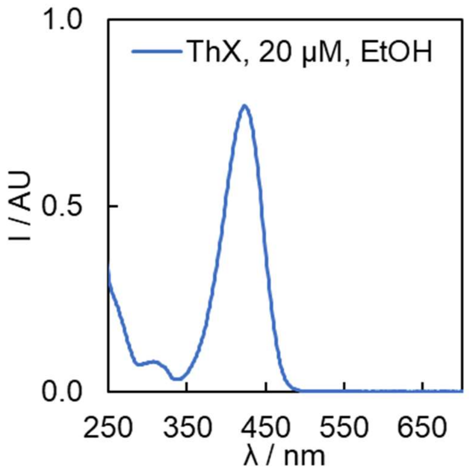

**Figure S23**. UV-vis spectrum of **ThX** (20 μM) in EtOH at 298 K, with $\lambda_{max}$ = 424 nm.

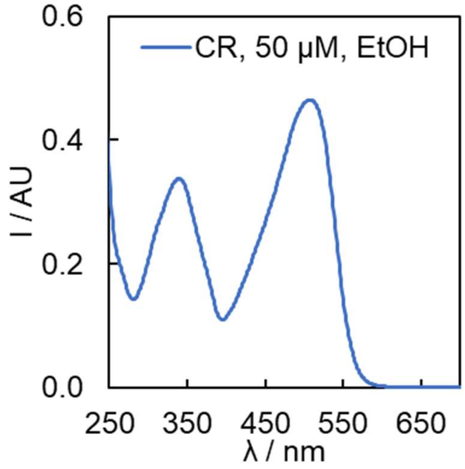

**Figure S24**. UV-vis spectrum of **CR** (50 μM) in EtOH at 298 K, with $\lambda_{max}$ = 508 nm.

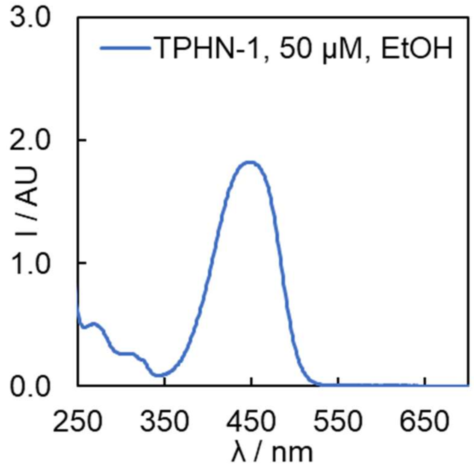

**Figure S25**. UV-vis spectrum of **TPHN-1** (50 μM) in EtOH at 298 K, with $\lambda_{max}$ = 448 nm.

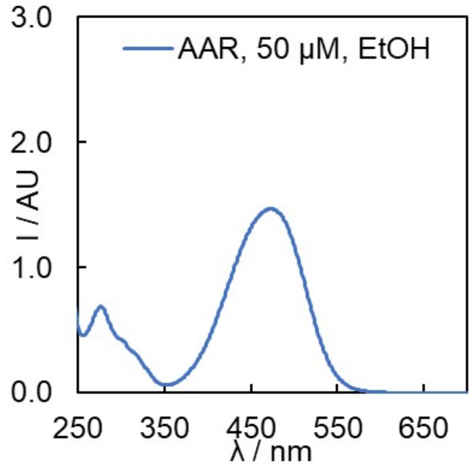

**Figure 26**. UV-vis spectrum of **AAR** (50 μM) in EtOH at 298 K, with $\lambda_{max}$ = 472 nm.

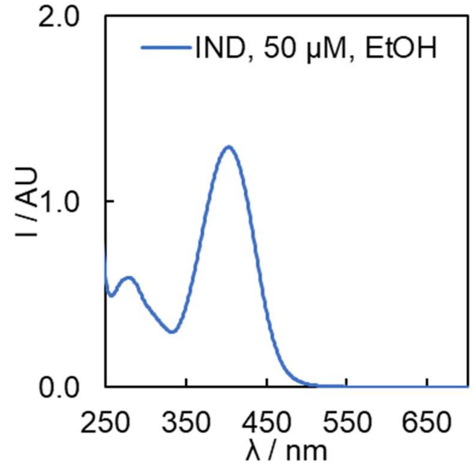

**Figure 27**. UV-vis spectrum of **IND** (50 μM) in EtOH at 298 K, with $\lambda_{max}$ = 403 nm.

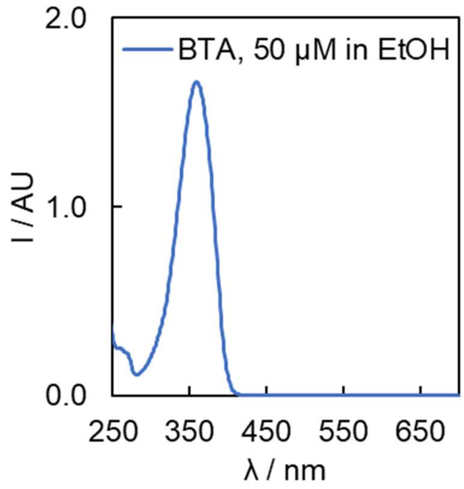

**Figure S28**. UV-vis spectrum of **BTA** (50 μM) in EtOH at 298 K, with $\lambda_{max}$ = 359 nm.

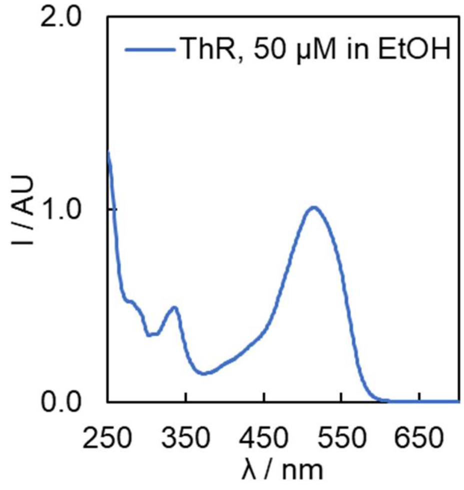

**Figure S29**. UV-vis spectrum of **ThR** (50 μM) in EtOH at 298 K, with $\lambda_{max}$ = 515 nm.

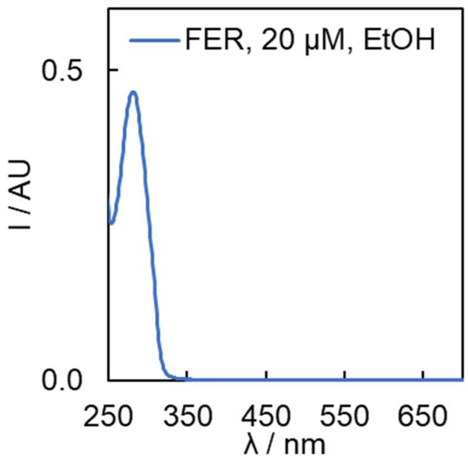

**Figure S30**. UV-vis spectrum of **FER** (20 μM) in EtOH at 298 K, with $\lambda_{max}$ = 299 nm.

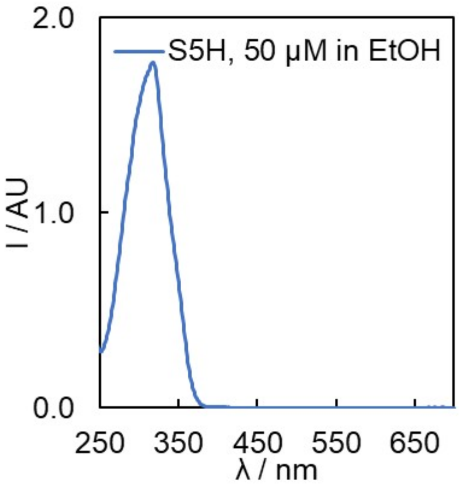

**Figure S31**. UV-vis spectrum of **S5H** (50 μM) in EtOH at 298 K, with $\lambda_{max}$ = 318 nm.

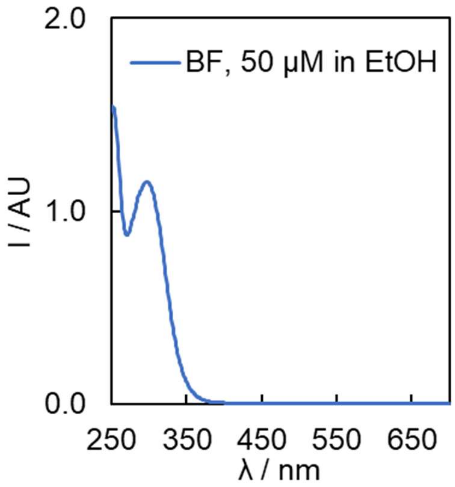

**Figure S32**. UV-vis spectrum of **BF** (50 μM) in EtOH at 298 K, with $\lambda_{max}$ = 251 nm.

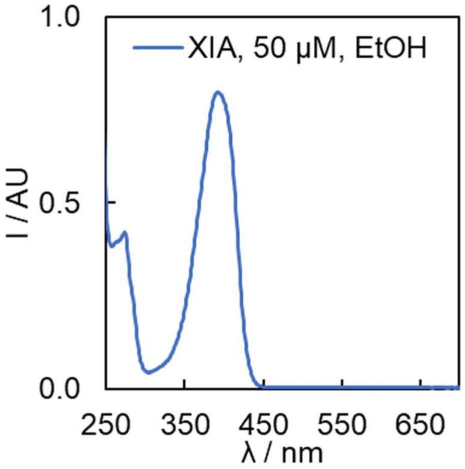

**Figure S33**. UV-vis spectrum of **XIA** (50 μM) in EtOH at 298 K, with $\lambda_{max}$ = 392 nm.

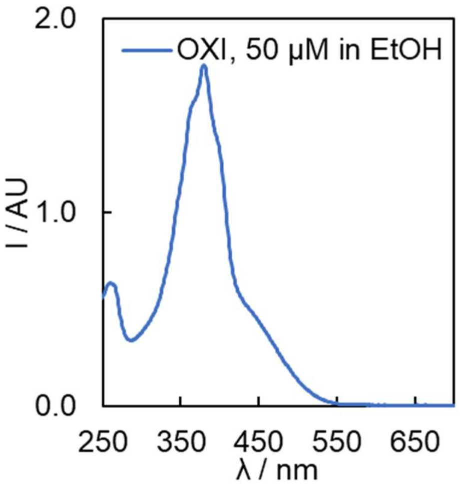

**Figure S34**. UV-vis spectrum of **OXI** (50 µM) in EtOH at 298 K, with $\lambda_{max}$ = 380 nm.

**Table S1.** UV-vis absorbance maxima of the ligands in EtOH.

| Ligand | $\lambda_{max}$ / nm |
|---|---|
| **ThX** | 424 |
| **CR** | 508 |
| **TPHN-1** | 448 |
| **AAR** | 472 |
| **IND** | 403 |
| **BTA** | 359 |
| **ThR** | 515 |
| **FER** | 299 |
| **S5H** | 318 |
| **BF** | 299 |
| **XIA** | 392 |
| **OXI** | 380 |

#### Fluorescence Characterisation

Fluorescence spectral readings to obtain excitation and emission spectra were performed on an Agilent Cary Eclipse Fluorescence Spectrophotometer using a scan rate of 600 nm/min, a data interval of 1.0 nm and an averaging time of 0.10 at 25°C. Fluorescence experiments used 20 nm excitation and emission slits and medium PMT voltage, except when specified otherwise. Samples were prepared in a Hellma quartz cuvette (2 mL) in spectroscopy-grade ethanol.

Fluorescence spectral readings to obtain emission spectra in the presence and absence of αSyn were performed using a BMG Labtech CLARIOstar Plus plate reader. Solutions of ligand (2.5 μM) and optionally αSyn (1.0 μM) in 1xPBS (pH 7.4) were prepared and aliquoted in a 384-well plate. Final solution volumes of 40 μL were used. Measurements were made through the top optic, and focal height and gain were adjusted based on the well containing αSyn. Four independent wells were prepared and measured, and the resultant spectra were averaged.

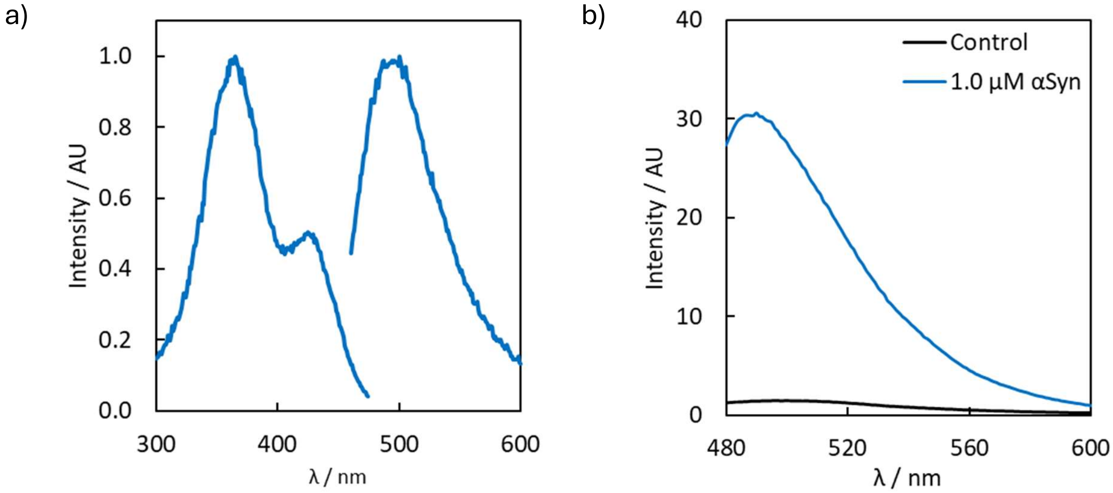

**Figure S35.** Fluorescence spectra of (a) **ThX** (50 μM) in EtOH ($\lambda_{ex}$ = 440, $\lambda_{em}$ = 399), recorded on a fluorimeter, and (b) emission spectra ($\lambda_{ex}$ = 440) of **ThX** (2.5 μM) in 1xPBS in the presence (blue line) and absence (black line) of αSyn (1.0 μM), recorded on a plate reader (focal height = 8.6 mm, gain = 1319).

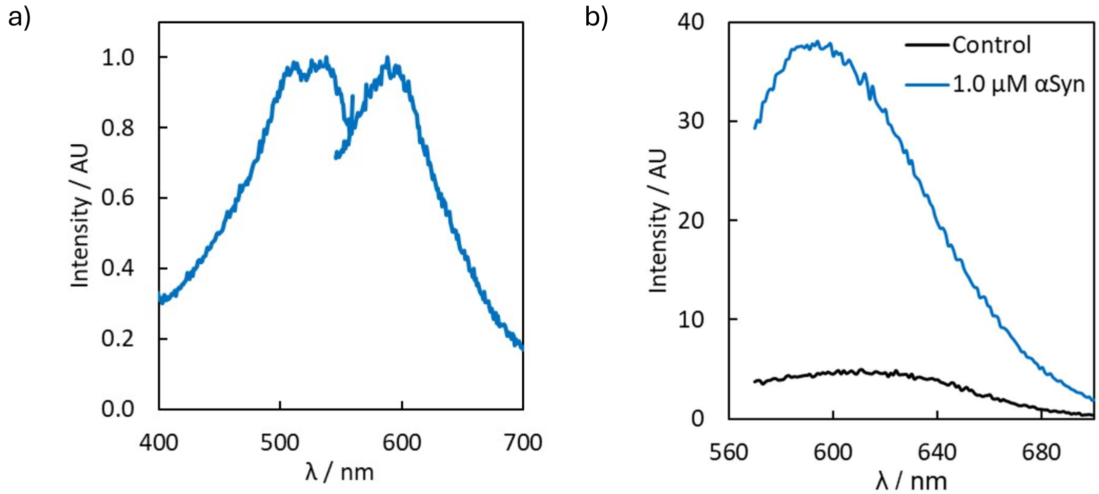

**Figure S36**. Fluorescence spectra of (a) **CR** (10 μM) in EtOH ($\lambda_{ex}$ = 530, $\lambda_{em}$ = 588), recorded on a fluorimeter, and (b) emission spectra ($\lambda_{ex}$ = 530) of **CR** (2.5 μM) in 1xPBS in the presence (blue line) and absence (black line) of αSyn (1.0 μM), recorded on a plate reader (focal height = 8.5 mm, gain = 2344).

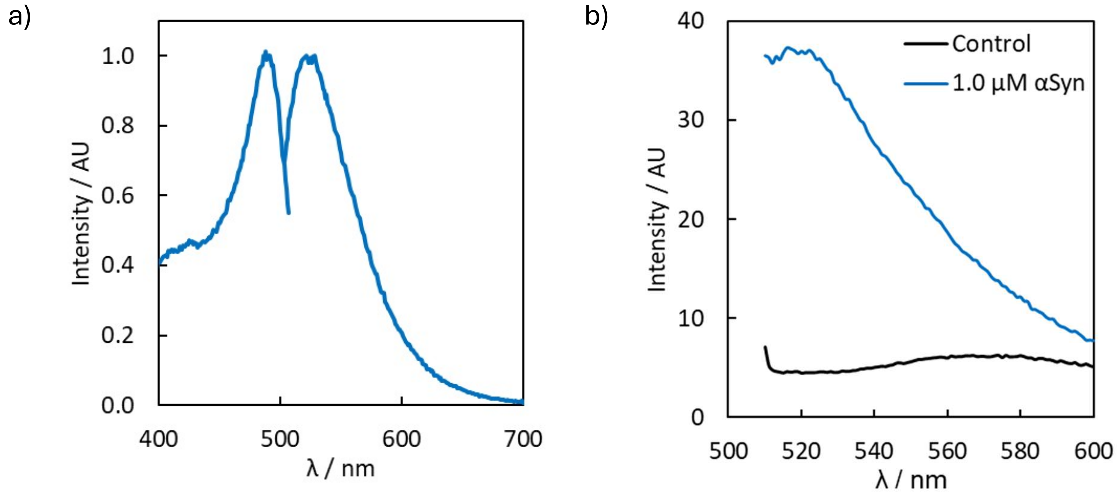

**Figure S37**. Fluorescence spectra of (a) **TPHN-1** (50 μM) in EtOH ($\lambda_{ex}$ = 488, $\lambda_{em}$ = 528), recorded on a fluorimeter, and (b) emission spectra ($\lambda_{ex}$ = 475) of **TPHN-1** (2.5 μM) in 1xPBS in the presence (blue line) and absence (black line) of αSyn (1.0 μM), recorded on a plate reader (focal height = 8.5 mm, gain = 1883).

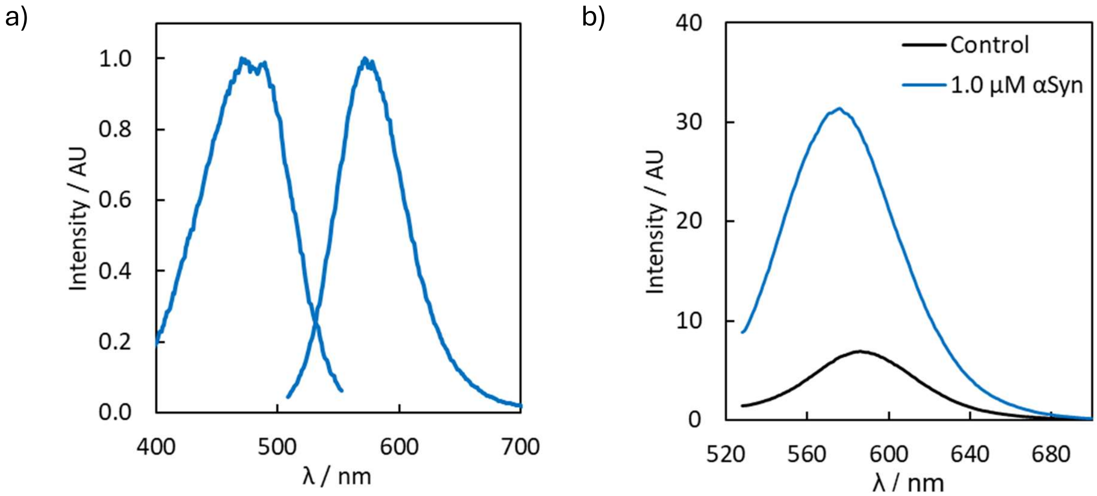

**Figure S38**. Fluorescence spectra of (a) **AAR** (50 μM) in EtOH ($\lambda_{ex}$ = 488, $\lambda_{em}$ = 572), recorded on a fluorimeter, and (b) emission spectra ($\lambda_{ex}$ = 488) of **AAR** (2.5 μM) in 1xPBS in the presence (blue line) and absence (black line) of αSyn (1.0 μM), recorded on a plate reader (focal height = 8.6 mm, gain = 1621).

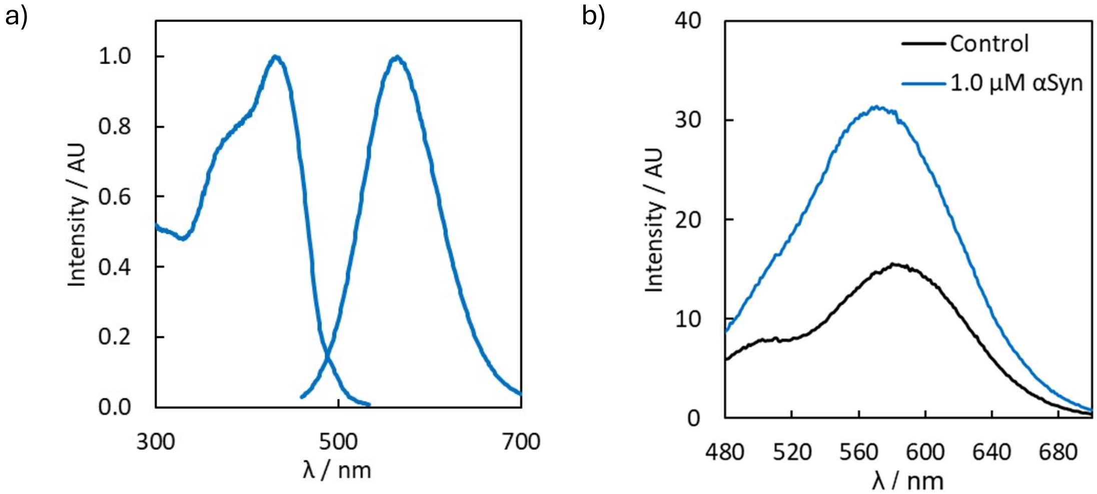

**Figure S39**. Fluorescence spectra of (a) **IND** (50 μM) in EtOH ($\lambda_{ex}$ = 430 nm, $\lambda_{em}$ = 564 nm), recorded on a fluorimeter, and (b) emission spectra ($\lambda_{ex}$ = 430) of **IND** (2.5 μM) in 1xPBS in the presence (blue line) and absence (black line) of αSyn (1.0 μM), recorded on a plate reader (focal height = 8.5 mm, gain = 1743).

#### Plate Reader Operation

All plate reader experiments were performed using a BMG Labtech CLARIOstar Plus plate reader with Corning 384-well black non-binding surface plates with clear flat bottoms (REF: 3766, LOT: 00422037). Fluorescence measurements were made through the top optic. Standardised focal heights and gains were used for each ligand, with settings summarised in Table S2.

**Table S2.** The plate reader settings used for data collection.

| | **ThX** | **CR** | **TPHN-1** | **AAR** | **IND** |
|---|---|---|---|---|---|
| Focal height / mm | 8.6 | 8.5 | 8.5 | 8.6 | 8.5 |
| Gain | 1319 | 2344 | 1883 | 1621 | 1743 |
| $\lambda_{ex}$ / nm | 440 | 530 | 475 | 488 | 430 |
| $\lambda_{em}$ range / nm | 480-530 | 570-630 | 510-560 | 550-610 | 530-610 |
| Bandwidth / nm | 20 | 25 | 25 | 25 | 25 |

### Ligand Control Measurements

Solutions (40 μL) of a fluorescent reporting ligand (2.0 μM) and competing ligand (5.0 μM) in 1xPBS (pH 7.4) were prepared and aliquoted into 384-well plates. The fluorescence of the reporting ligand was then measured, as above. Four independent wells were prepared for each combination of reporting and competing ligand.

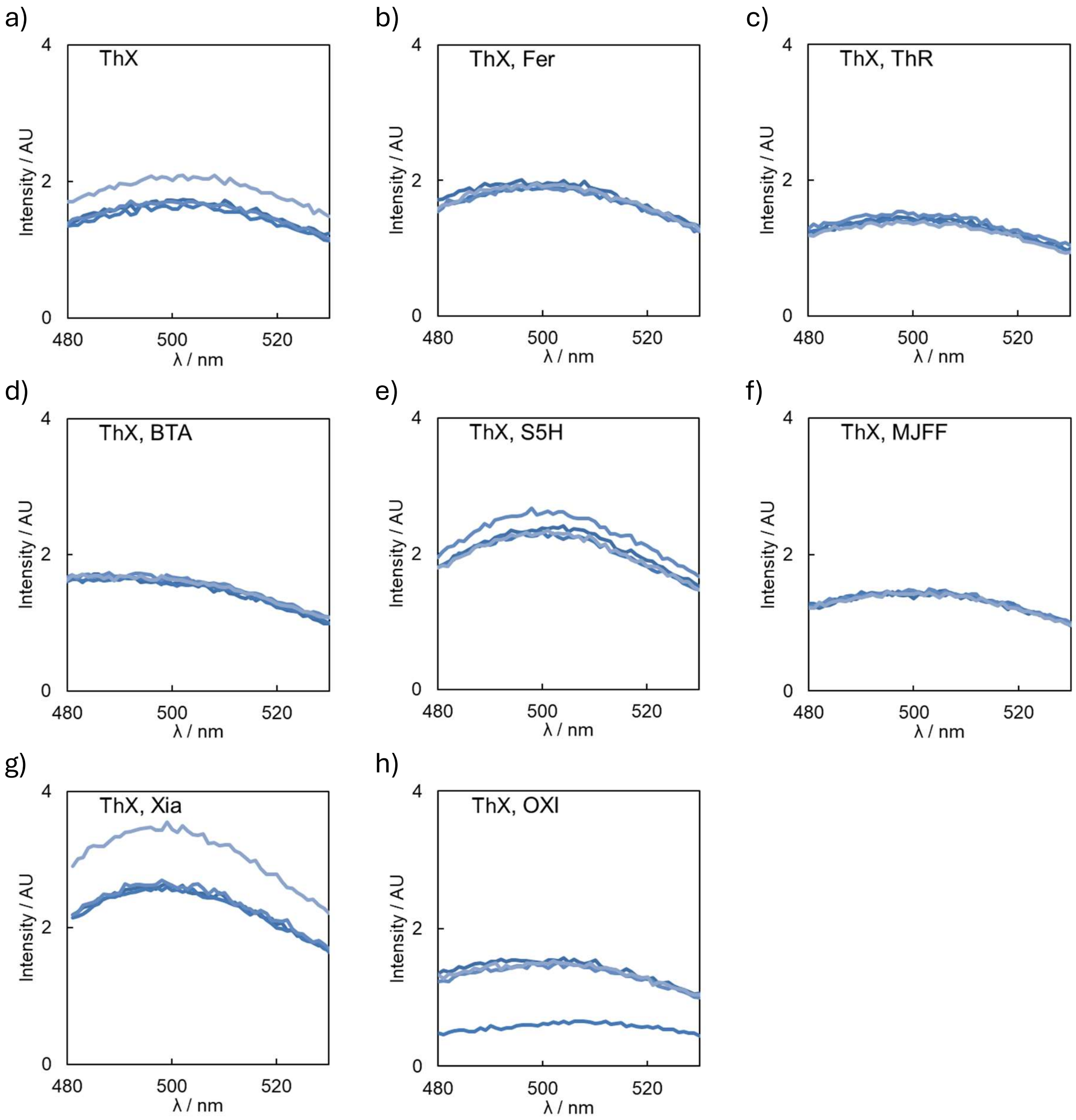

**Figure S40**. Fluorescence emission spectra for **ThX** (2.0 μM, $\lambda_{ex}$ = 440 nm) and competing ligands (5.0 μM) in 1xPBS (pH 7.4). Each graph shows four overlaid traces.

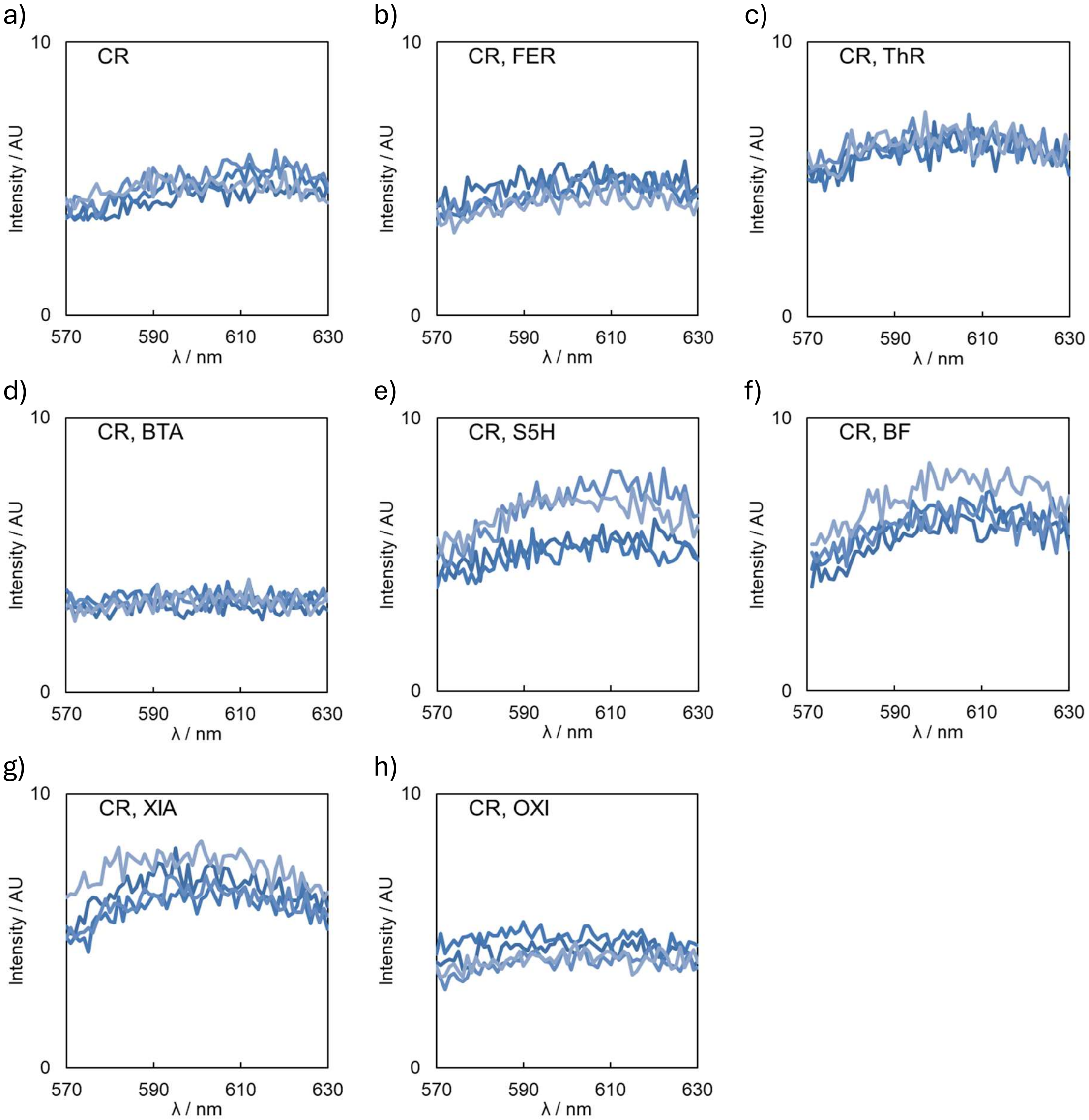

**Figure S41**. Fluorescence emission spectra for **CR** (2.0 μM, $\lambda_{ex}$ = 530 nm) and competing ligands (5.0 μM) in 1xPBS (pH 7.4). Each graph shows four overlaid traces.

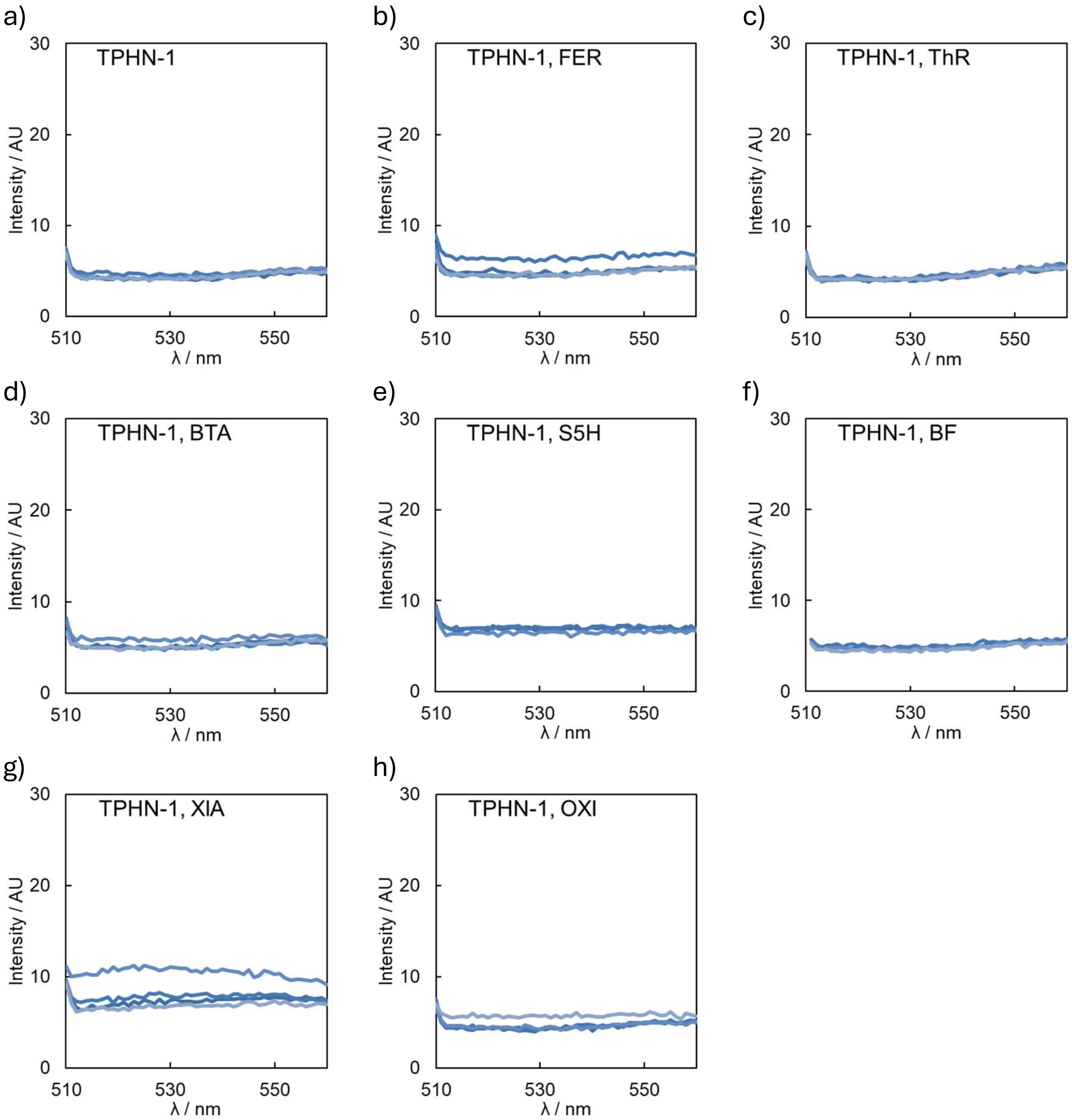

**Figure S42**. Fluorescence emission spectra for **TPHN-1** (2.0 μM, $\lambda_{ex}$ = 475 nm) and competing ligands (5.0 μM) in 1xPBS (pH 7.4). Each graph shows four overlaid traces.

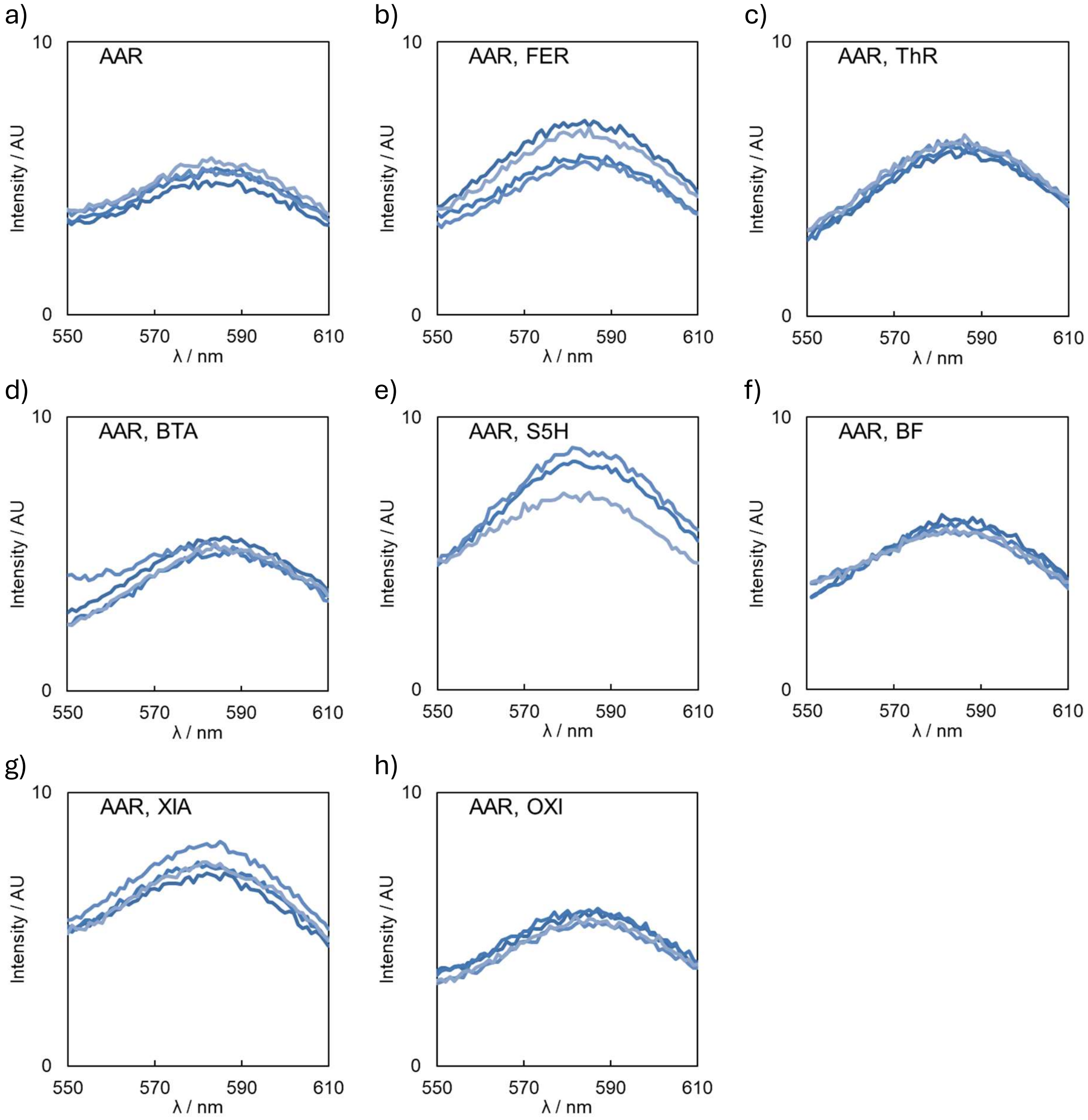

**Figure S43**. Fluorescence emission spectra for **AAR** (2.0 μM, $\lambda_{ex}$ = 488 nm) and competing ligands (5.0 μM) in 1xPBS (pH 7.4). Each graph shows four overlaid traces.

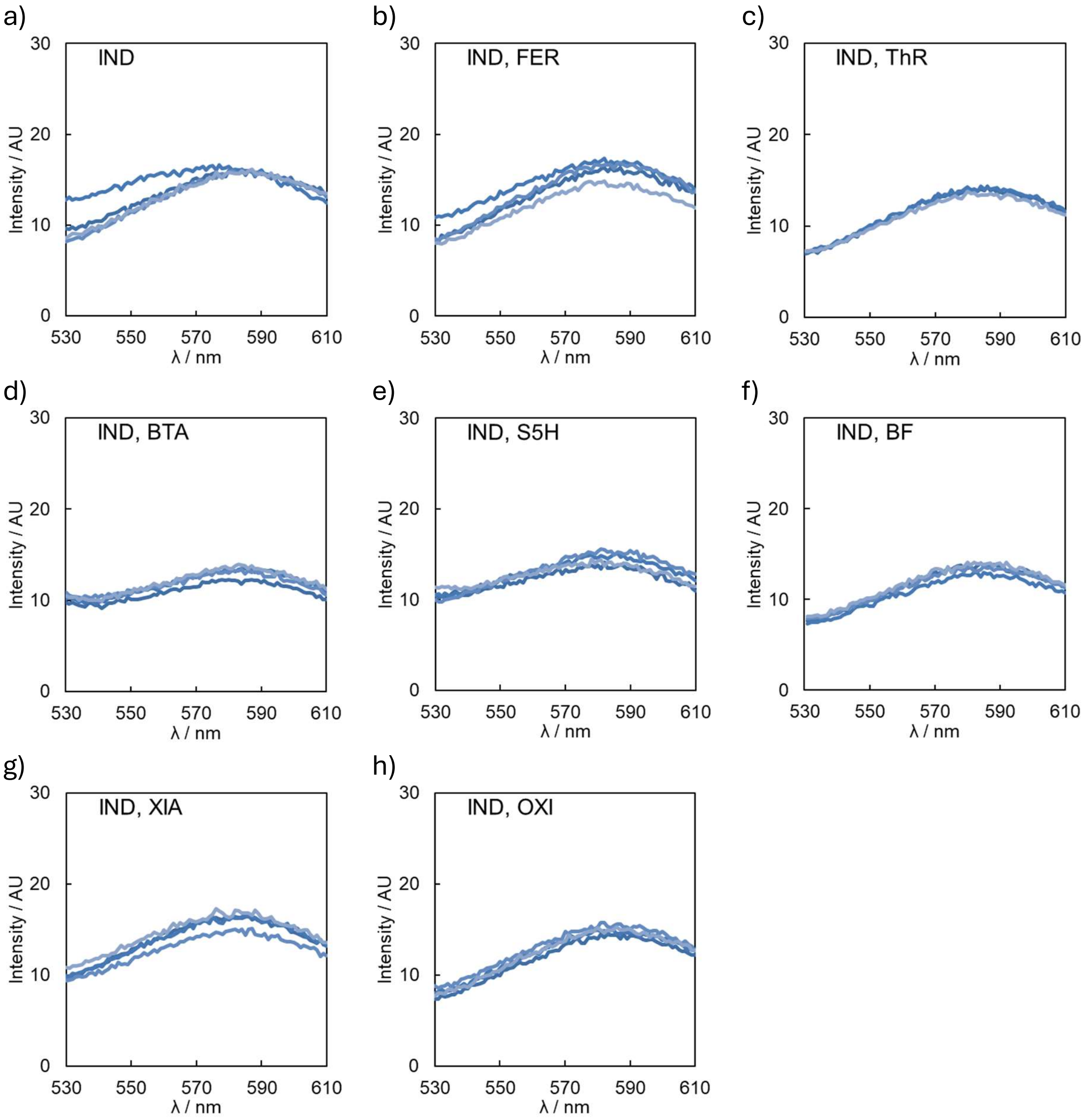

**Figure S44**. Fluorescence emission spectra for **IND** (2.0 μM, $\lambda_{ex}$ = 430 nm) and competing ligands (5.0 μM) in 1xPBS (pH 7.4). Each graph shows four overlaid traces.

#### Ligand Monomer Binding

Solutions (40 µL) of a fluorescent reporting ligand (2.0 µM), competing ligand (5.0 µM), and monomeric αSyn (1.0 µM) in 1xPBS (pH 7.4) were prepared and aliquoted into 384-well plates. The fluorescence of the reporting ligands was then measured (Figure S45). Four independent wells were prepared for each combination of reporting and competing ligand. Only **CR** demonstrated a change in fluorescence in the presence of monomeric αSyn.

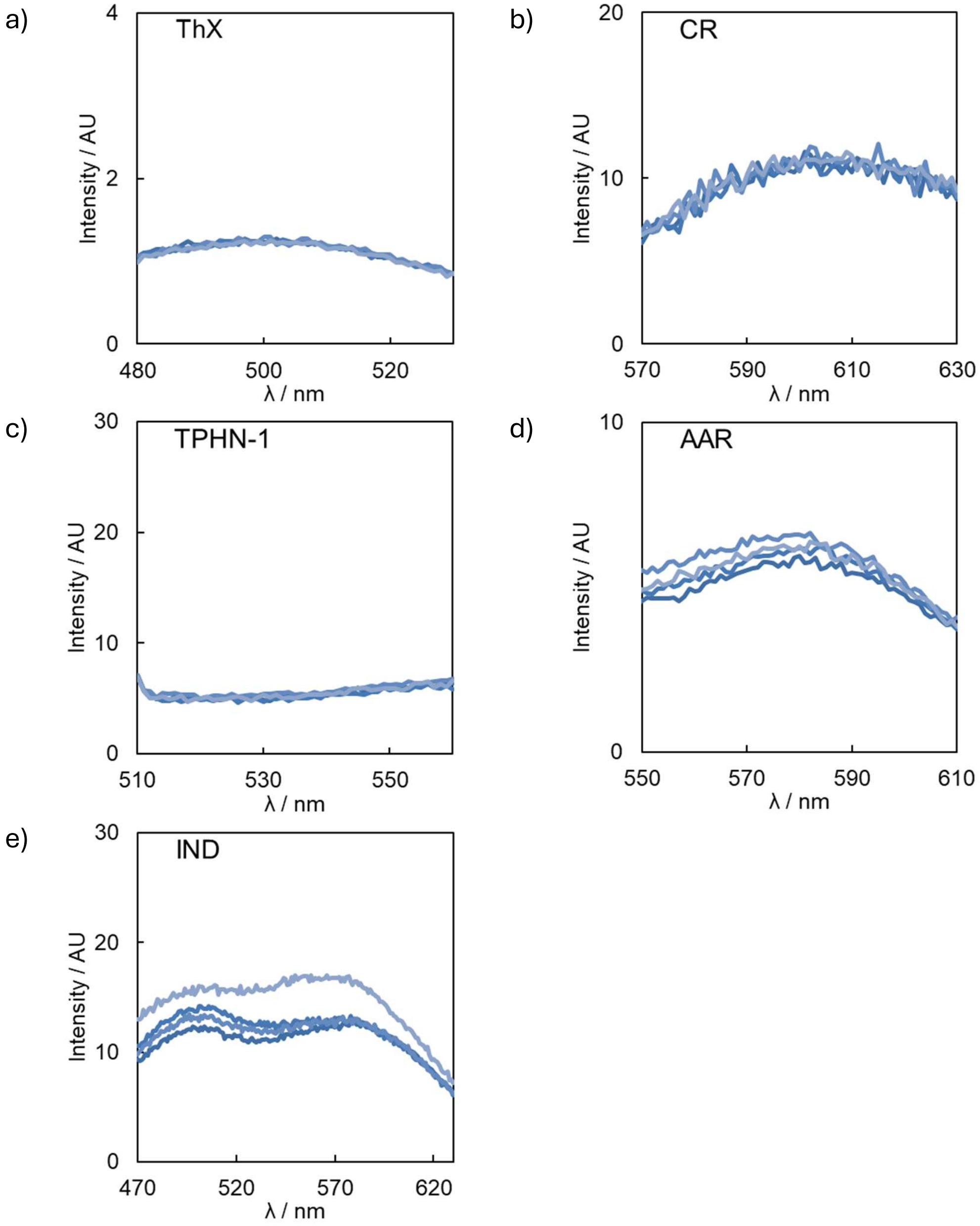

**Figure S45**. Fluorescence emission spectra for 2.0 µM of (a) ThX, (b) CR, (c) TPHN-1, (d) AAR, and (e) IND in the presence of monomeric αSyn (1.0 µM) in 1xPBS (pH 7.4). Each graph shows four overlaid traces.

#### Fitting of Kinetic Data

The fitted parameters and mean residual errors are summarised in Table S3.

**Table S3**. Fitted parameters and mean residual error from the fitting of kinetic data using AmyloFit.[11] MRE: mean residual error. ThX included: kinetic data when ThX was included within the aggregation reaction mixture. CD: circular dichroism.

| | **ThX** | **ThX** included | **CR** | **TPHN-1** | **AAR** | **IND** | CD |
|---|---|---|---|---|---|---|---|
| $k_+k_n$ | 6.81E-06 | 8.67E-07 | 2.19E-03 | 2.26E-04 | 1.61E-04 | 2.27E-03 | 1.52E-04 |
| $n_c$ | 0.00218 | 0.0207 | 0.381 | 0.116 | 0.113 | 0.323 | 2.53E-05 |
| $k_+k_2$ | 4.93E+05 | 1.04E+05 | 8.33E+10 | 1.34E+03 | 3.70E+06 | 6.02E+06 | 8.79E+05 |
| $n_2$ | 0.711 | 0.344 | 1.29 | 0.804 | 1.5 | 7.84 | 1.05 |
| MRE | 0.00199 | 0.0114 | 0.0143 | 0.0161 | 0.018 | 0.00873 | 0.0335 |

### Biophysical Characterisation of αSyn Aggregates

#### *Circular Dichroism*

A 20 μL aliquot of the aggregation reaction mixture was taken and diluted to 1 mL with 10 mM sodium phosphate buffer (pH 7.4). The solution was then passed through an Amicon Ultra 0.5 mL 100 kDa nominal molecular weight limit (NMWL) centrifugal filter (13,000 g x 5 min). The concentrate was then diluted with 10 mM sodium phosphate buffer (pH 7.4) to a final volume of 1 mL, and the process repeated twice more. The final concentrate was diluted with 10 mM sodium phosphate buffer (pH 7.4) to a final volume of 1 mL and circular dichroism (CD) spectra were recorded with a Chirascan CD1 Spectrometer (Applied Photonics Ltd.) equipped with a Series 800 Temperature Controller (Alpha Omega Instruments). Far-ultraviolet measurements (190-250 nm) were recorded at 25 °C with a 1.0 cm optical pathlength, a time-per-point of 1.0 s, a 1.0 nm bandwidth, and a wavelength step of 0.1 nm. CD spectra were averaged over six scans. Data were baseline corrected by subtracting the complete buffer spectrum of 10 mM sodium phosphate buffer (pH 7.4) averaged over six scans. Applied Photophysics Pro-Data Chirascan software was used to smooth the data using Savitsky-Golay smoothing and a window size of eight, and the data was converted to molar ellipticity.

CD spectra obtained at different timepoints are shown in Figure S46, and CD spectra of stored αSyn aggregates are shown in Figure S47.

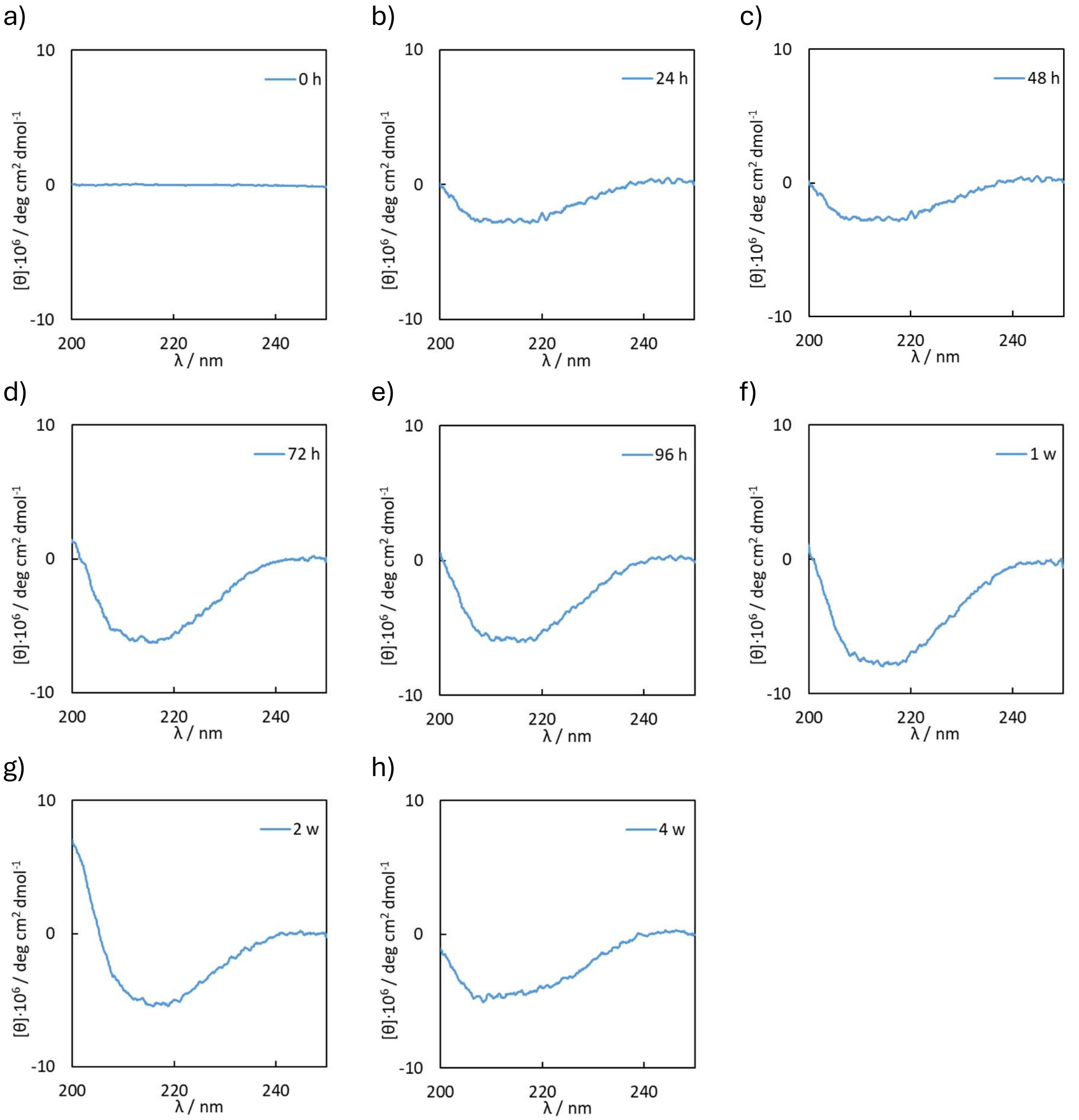

**Figure S46**. CD spectra obtained from aliquots taken at different timepoints: (a) 0 h, (b) 24 h, (c) 48 h, (d) 72 h, (e) 96 h, (f) 1 week, (g) 2 weeks, (h) 4 weeks.

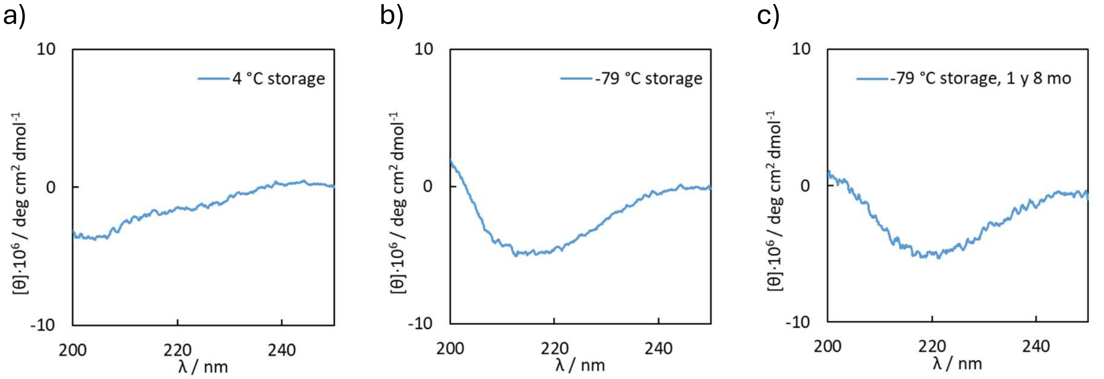

**Figure S47**. CD spectra obtained from aliquots of 96 h αSyn aggregates stored for 25 days at (a) 4 °C or (b) -79 °C, or (c) a separate αSyn aggregate preparation stored for 1 year and 8 months.

#### *Transmission Electron Microscopy*

Nanoscale morphologies of fibril samples were observed by transmission electron microscopy (TEM) using a Thermo Scientific (FEI Company) Talos F200X G2 microscope operating at 200 kV. Images were recorded with a Ceta 4k x 4k CMOS camera. For sample preparation, TEM grids (continuous carbon film on 300 mesh Cu) were glow discharged using a Quorom Technologies GloQube at 25 mA for 60 s. A 2 μL sample of the aggregation reaction mixture diluted into distilled water (1.0 μM concentration with respects to monomeric αSyn) was placed on a freshly glow-discharged grid, and after 30 s residual fluid was carefully removed by blotting with filter paper. The sample was negatively stained using 2.0 μL of 2% (w/v) uranyl acetate solution in ethanol for 30 s. The grid was blotted and dried in air for 10 min at room temperature before use.

Representative TEM micrographs for 72 h aggregation timepoints are shown in Figure S48. Representative TEM micrographs of αSyn aggregates after storage at -79 °C for 1 year and 8 months upon different numbers of freeze-thaw cycles are shown in Figure S49. The distribution of fibril widths in obtained TEM micrographs are shown for the aggregation of αSyn in the absence of **ThX** (Figure S50) and in the presence of **ThX** (Figure S51). The distribution of fibril widths in obtained TEM micrographs are shown for αSyn aggregates prepared in the presence of **ThX** then stored at either 4 °C or -79 °C (Figure S52).

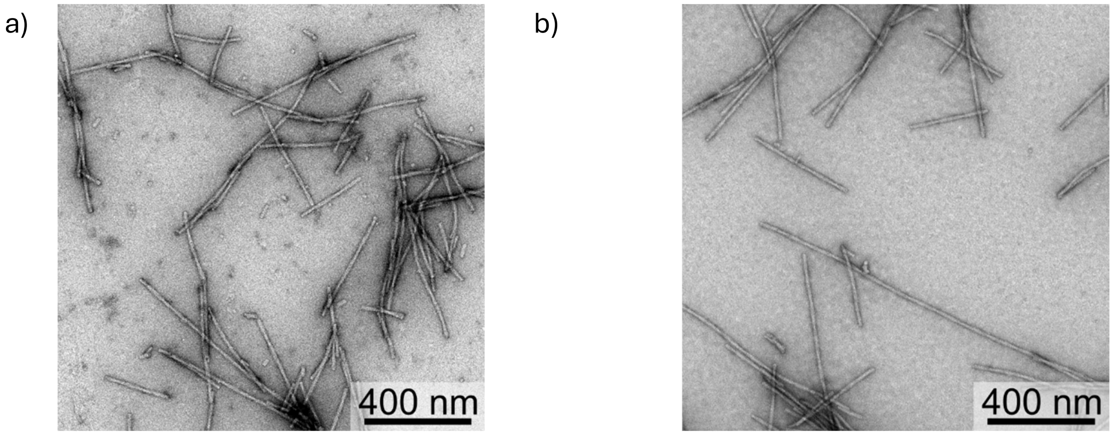

**Figure S48**. Representative TEM micrographs of αSyn aggregates obtained after 72 h after aggregation (a) without **ThX**, and (b) in the presence of 20 μM **ThX**.

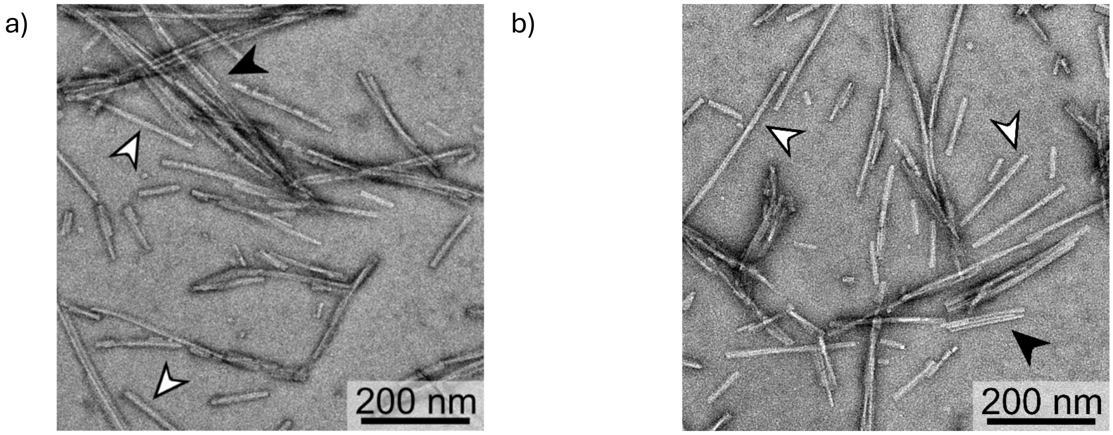

**Figure S49**. Representative TEM micrographs of αSyn aggregates after storage at -79 °C for 1 year and 8 months after a total of (a) 3 freeze-thaw cycles, and (b) 5 freeze-thaw cycles. Black-filled arrow heads identify thin fibrils, white-filled arrows identify thick fibrils.

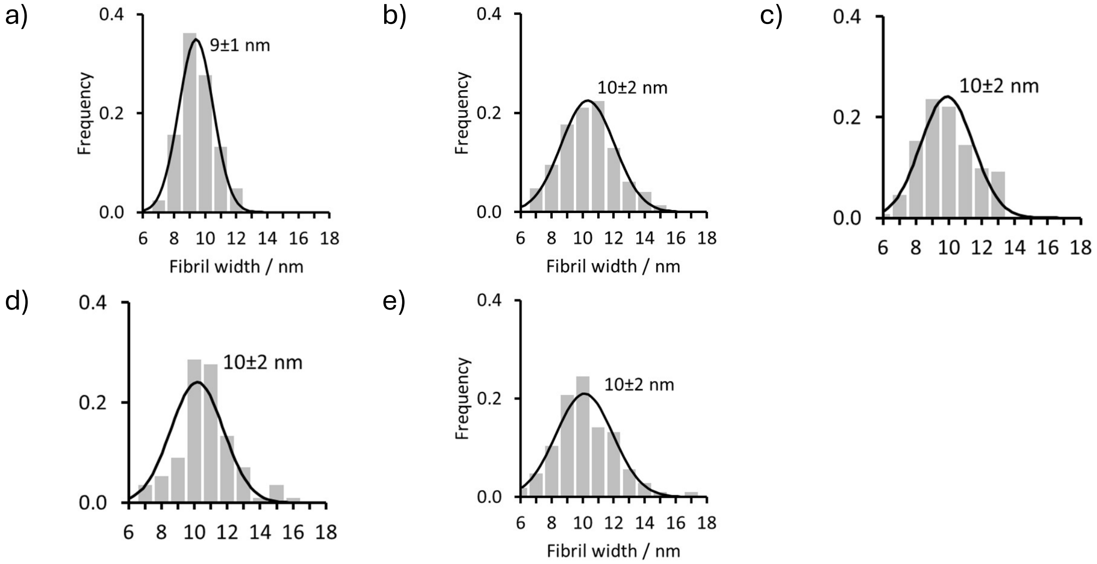

**Figure S50**. The distribution of αSyn fibril widths of (a) 24 h, (b) 48 h, (c) 72 h, (d) 96 h, and (e) 1 week aggregation timepoints. Fibril widths were measured as the full width half maximum (FWHM) in ImageJ,[12] and at least 100 measurements from multiple representative TEM micrographs were taken for each timepoint. Plotted black lines are the Gaussian distribution corresponding to the mean ± standard deviation shown on each graph. Aggregation was performed with 50 μM αSyn in 50 mM Tris and 1.5 mM $NaN_3$ (pH 7.4, 37 °C).

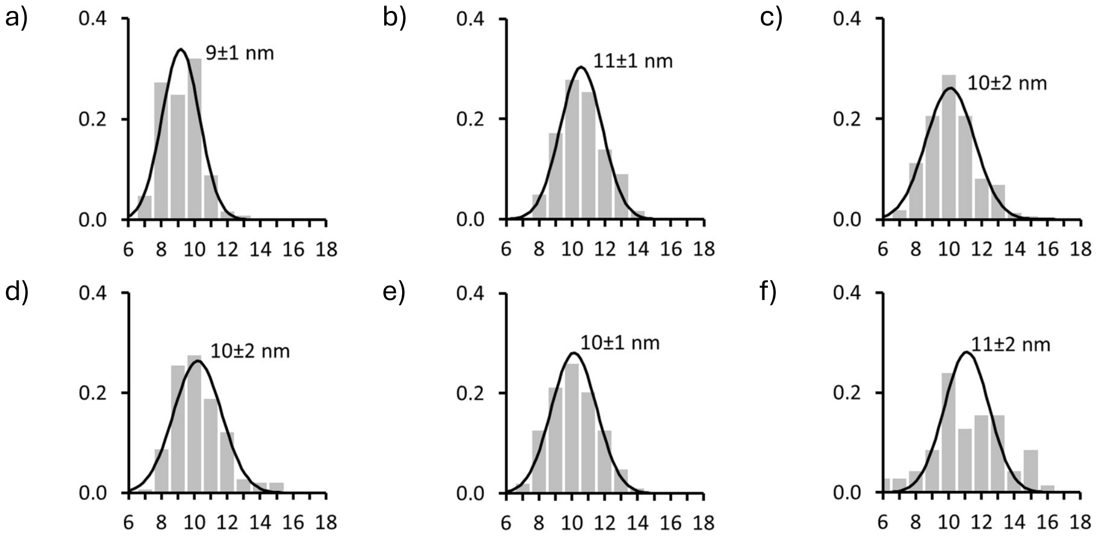

**Figure S51**. The distribution of αSyn fibril widths of (a) 24 h, (b) 48 h, (c) 72 h, (d) 96 h, (e) 1 week, and (f) 2 week aggregation timepoints. Fibril widths were measured as the full width half maximum (FWHM) in ImageJ,[12] and at least 100 measurements from multiple representative TEM micrographs were taken for each timepoint. Plotted black lines are the Gaussian distribution corresponding to the mean ± standard deviation shown on each graph. Aggregation was performed with 50 μM αSyn in 50 mM Tris and 1.5 mM $NaN_3$ with 20 μM **ThX** (pH 7.4, 37 °C).

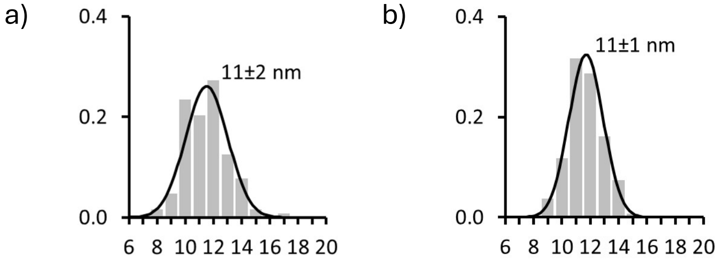

**Figure S52**. The distribution of αSyn fibril widths of 96 h αSyn aggregates after storage for 25 days at (c) -79 °C and (d) -4 °C. Fibril widths were measured as the full width half maximum (FWHM) in ImageJ,[12] and at least 100 measurements from multiple representative TEM micrographs were taken for each timepoint. Plotted black lines are the Gaussian distribution corresponding to the mean ± standard deviation shown on each graph. Aggregation was performed with 50 μM αSyn in 50 mM Tris and 1.5 mM $NaN_3$ with 20 μM **ThX** (pH 7.4, 37 °C).

### Fluorescence Data

#### *Ligand Profiling During Aggregation*

Aliquots of the αSyn aggregation mixture were diluted to a final concentration of 1.0 μM αSyn with 2.0 μM reporting ligand and optionally 5.0 μM competing ligand in 1xPBS (pH 7.4). The fluorescence of the resultant solution was measured using the plate reader. The change in fluorescence in the presence of the competing ligand is summarised in Figure S53 for **ThX**, Figure S54 for **ThX** when αSyn aggregates formed in the presence of **ThX** are used, Figure S55 for **CR**, Figure S56 for **TPHN-1**, Figure S57 for **AAR**, and Figure S58 for **IND**. Measurements are normalised to the fluorescence intensity measured in the absence of any competing ligand.

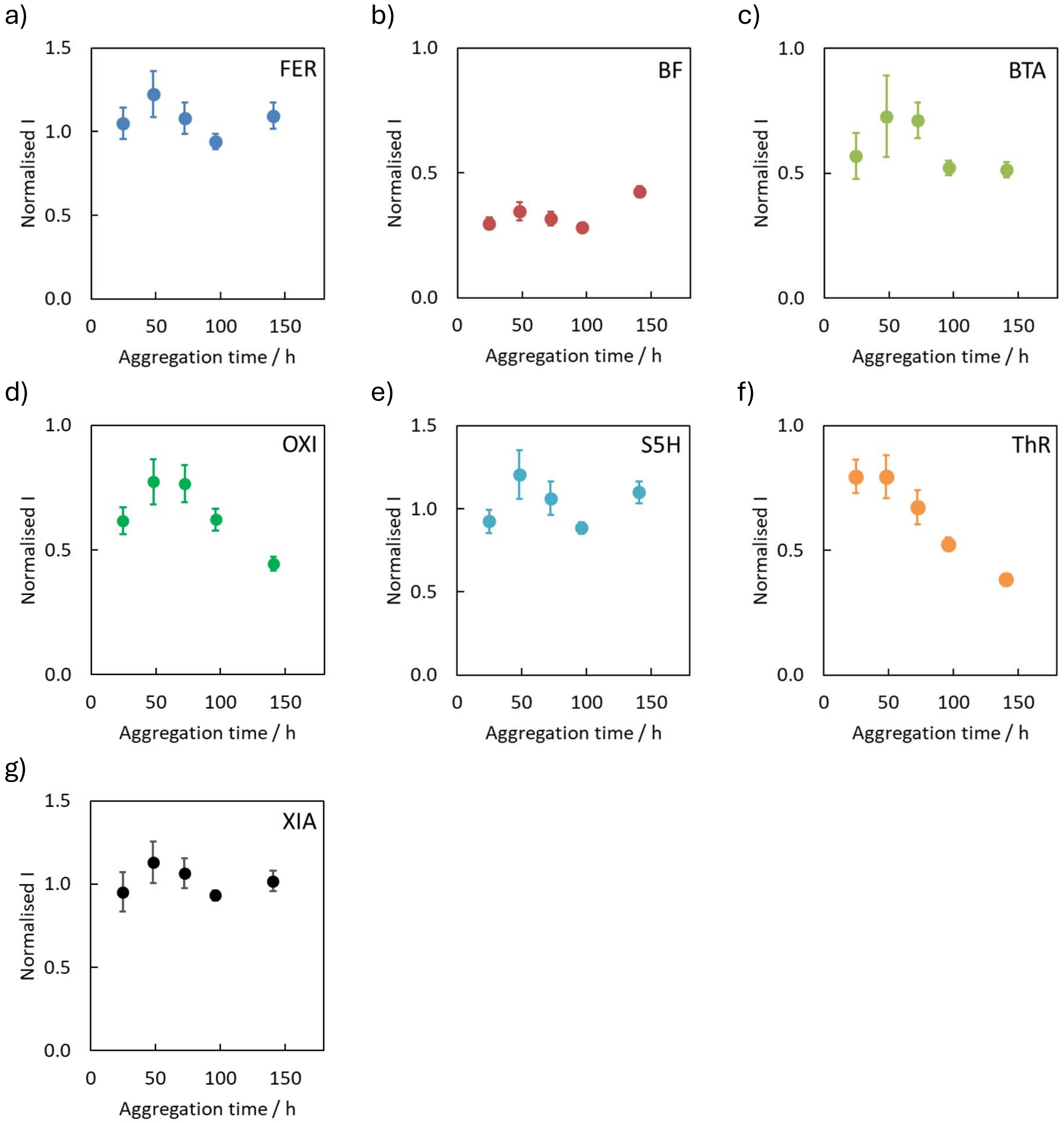

**Figure S53**. Fluorescence measurements for the displacement of **ThX** ($\lambda_{ex}$ = 440 nm, $\lambda_{em}$ = 488-492 nm) by competing ligands from αSyn fibrils at different aggregation timepoints. Aggregation was performed using 50 μM αSyn in 50 mM Tris with 1.5 mM $NaN_3$ (pH 7.4, 37 °C). The reaction was monitored by diluting aliquots to 1.0 μM αSyn in 1xPBS (pH 7.4, 25 °C) and adding 2.0 μM reporting ligand and 5.0 μM competing ligand. Data points are the average of at least three measurements with 95% confidence intervals shown.

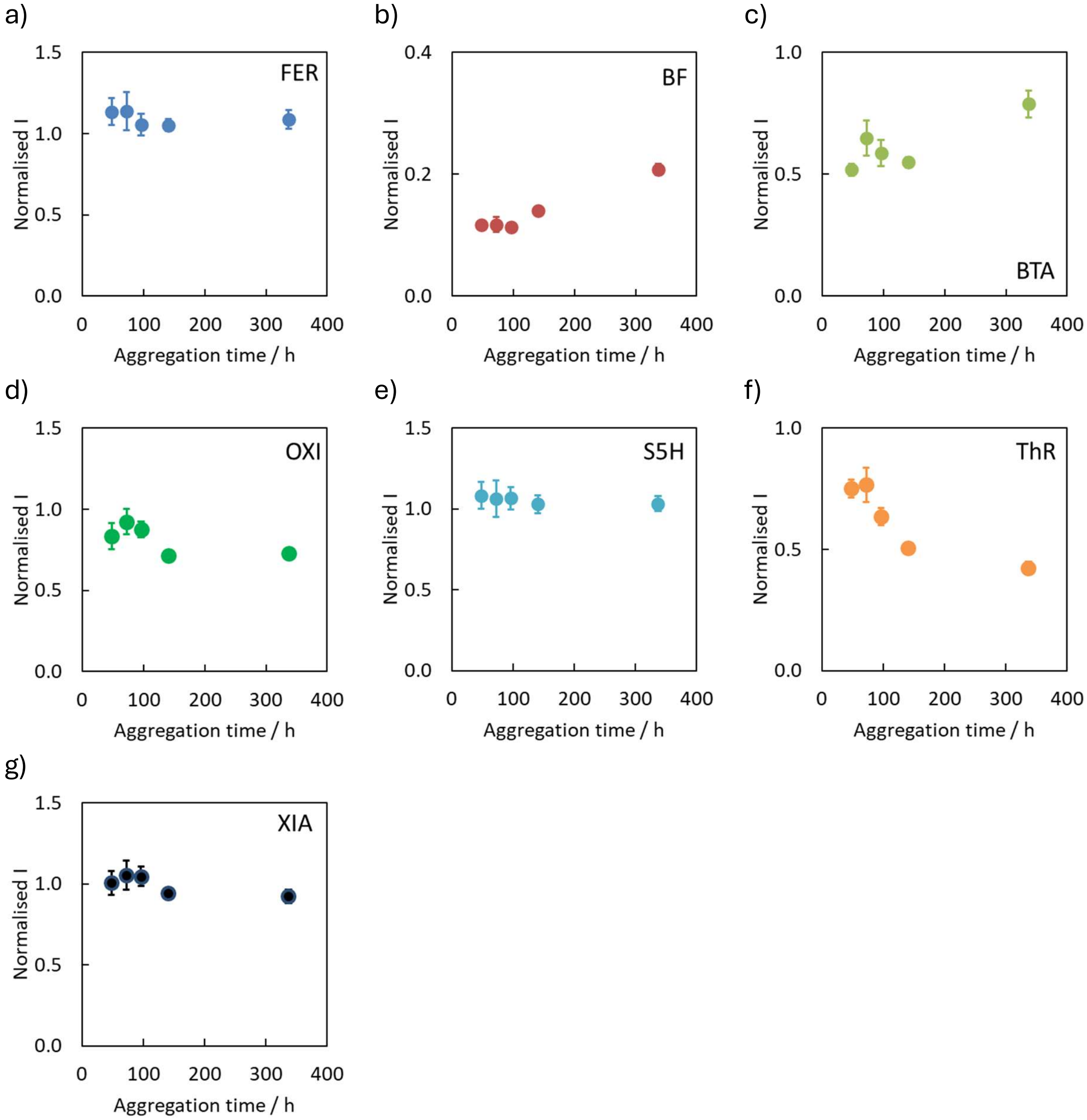

**Figure S54**. Fluorescence measurements for the displacement of **ThX** ($\lambda_{ex}$ = 440 nm, $\lambda_{em}$ = 488-492 nm) by competing ligands from αSyn fibrils at different aggregation timepoints. Aggregation was performed using 50 μM αSyn in 50 mM Tris with 1.5 mM $NaN_3$ in the presence of 20 μM **ThX** (pH 7.4, 37 °C). The reaction was monitored by diluting aliquots to 1.0 μM αSyn in 1xPBS (pH 7.4, 25 °C) and adding 2.0 μM reporting ligand and 5.0 μM competing ligand. Data points are the average of at least three measurements with 95% confidence intervals shown.

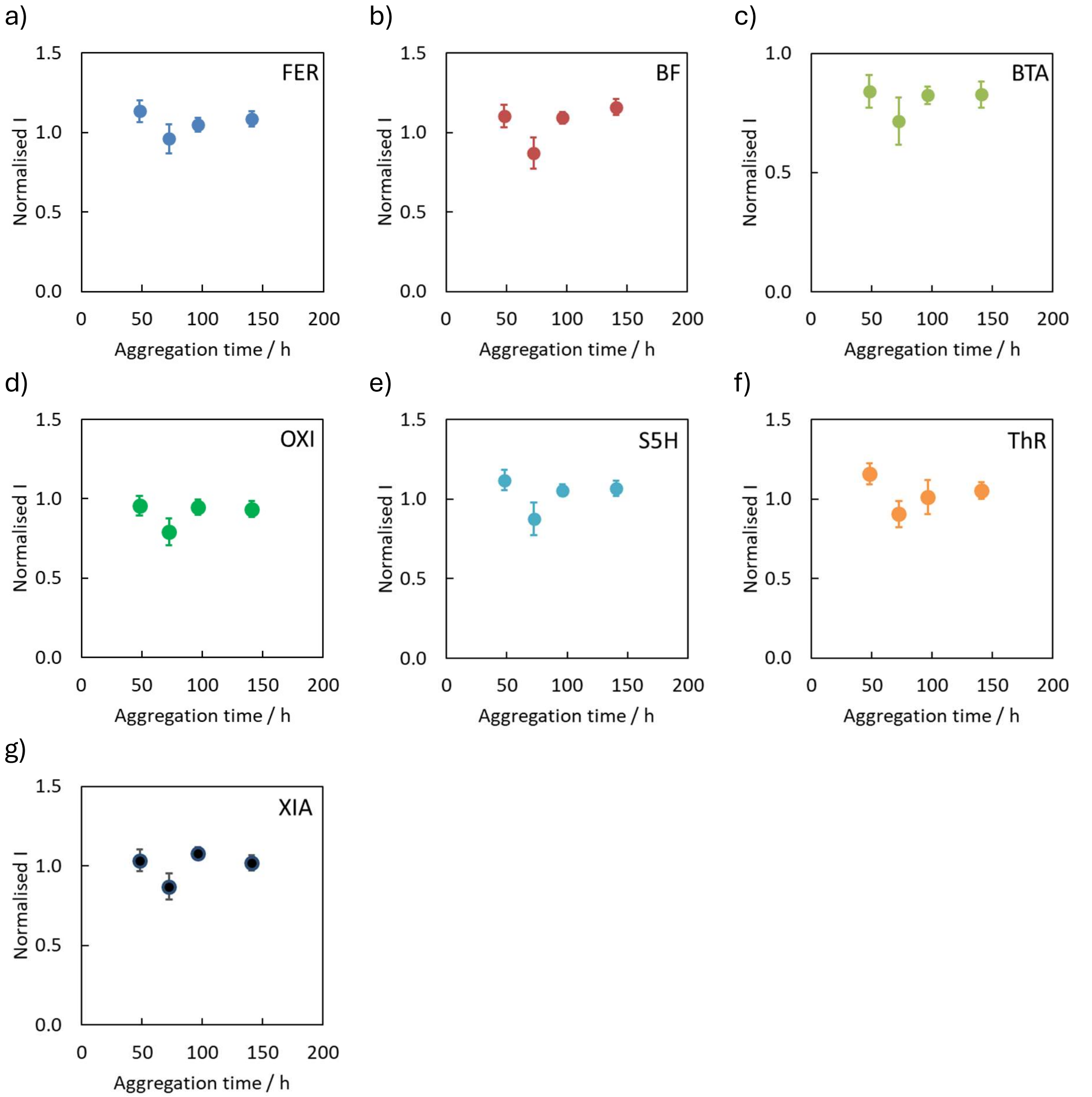

**Figure S55**. Fluorescence measurements for the displacement of **CR** ($\lambda_{ex}$ = 530 nm, $\lambda_{em}$ = 592-598 nm) by competing ligands from αSyn fibrils at different aggregation timepoints. Aggregation was performed using 50 μM αSyn in 50 mM Tris with 1.5 mM $NaN_3$ (pH 7.4, 37 °C). The reaction was monitored by diluting aliquots to 1.0 μM αSyn in 1xPBS (pH 7.4, 25 °C) and adding 2.0 μM reporting ligand and 5.0 μM competing ligand. Data points are the average of at least three measurements with 95% confidence intervals shown.

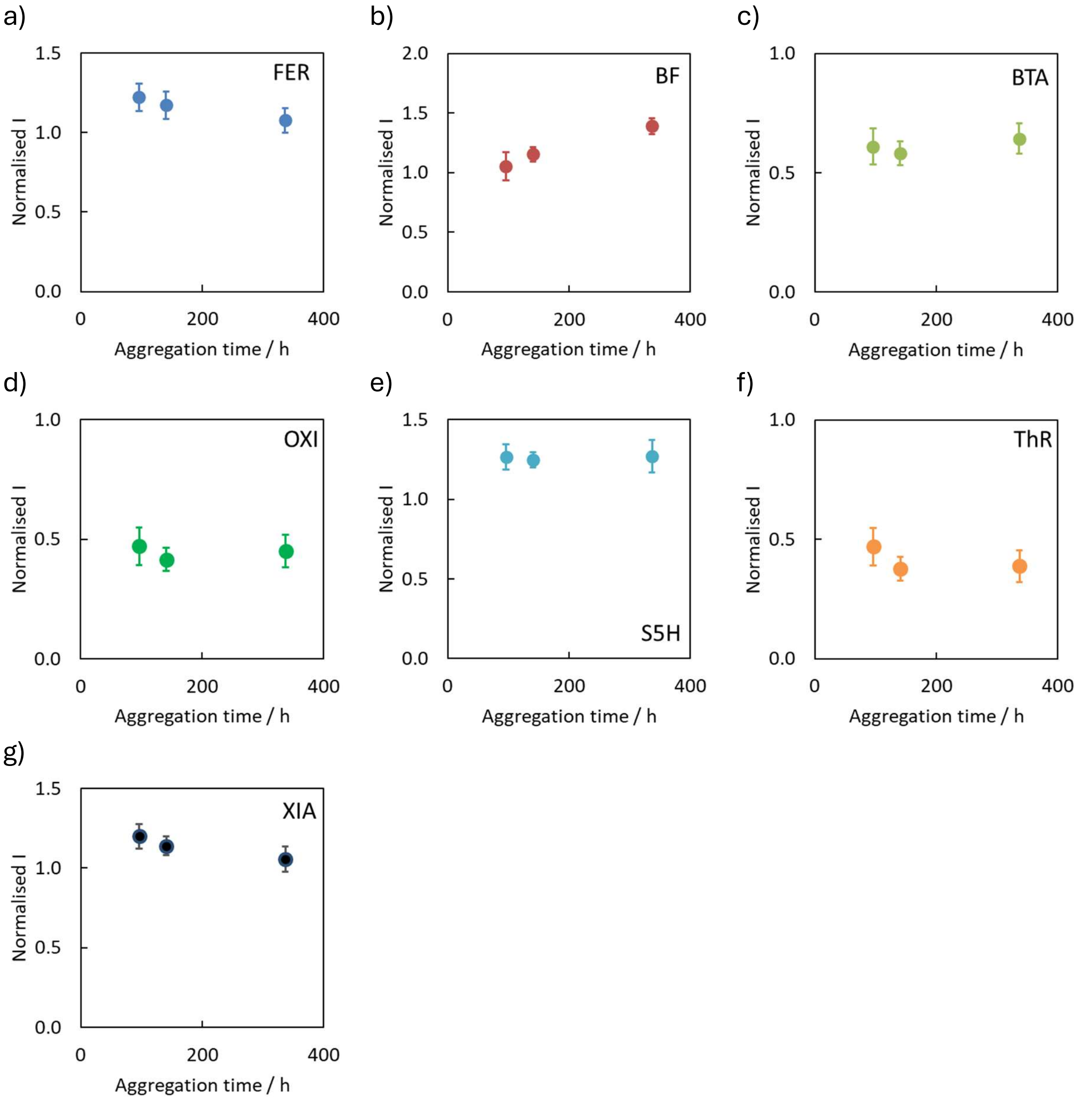

**Figure S56**. Fluorescence measurements for the displacement of **TPHN-1** ($\lambda_{ex}$ = 475 nm, $\lambda_{em}$ = 515-520 nm) by competing ligands from αSyn fibrils at different aggregation timepoints. Aggregation was performed using 50 µM αSyn in 50 mM Tris with 1.5 mM $NaN_3$ (pH 7.4, 37 °C). The reaction was monitored by diluting aliquots to 1.0 µM αSyn in 1xPBS (pH 7.4, 25 °C) and adding 2.0 µM reporting ligand and 5.0 µM competing ligand. Data points are the average of at least three measurements with 95% confidence intervals shown.

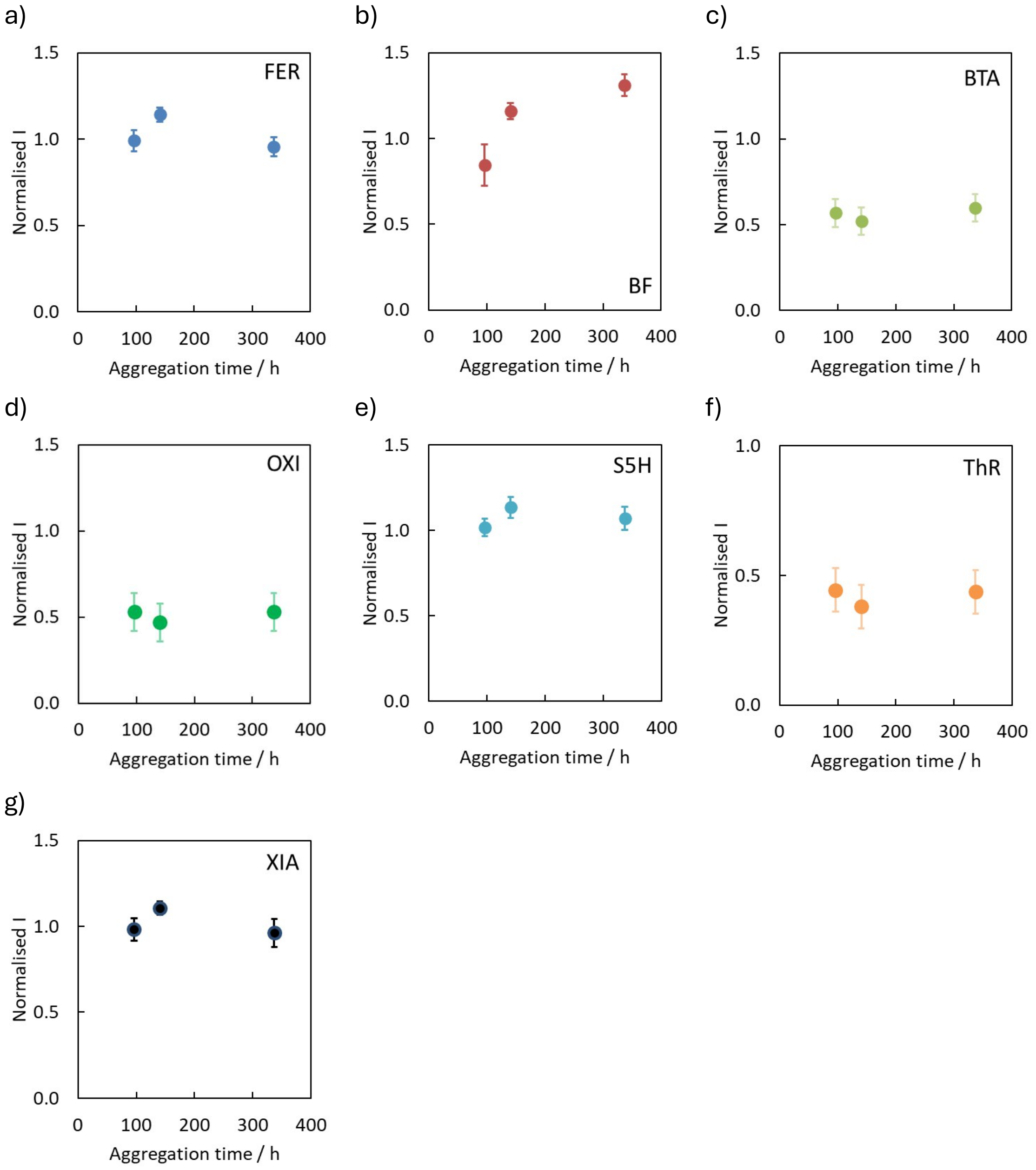

**Figure S57**. Fluorescence measurements for the displacement of **AAR** ($\lambda_{ex}$ = 488 nm, $\lambda_{em}$ = 592-598 nm) by competing ligands from αSyn fibrils at different aggregation timepoints. Aggregation was performed using 50 μM αSyn in 50 mM Tris with 1.5 mM $NaN_3$ (pH 7.4, 37 °C). The reaction was monitored by diluting aliquots to 1.0 μM αSyn in 1xPBS (pH 7.4, 25 °C) and adding 2.0 μM reporting ligand and 5.0 μM competing ligand. Data points are the average of at least three measurements with 95% confidence intervals shown.

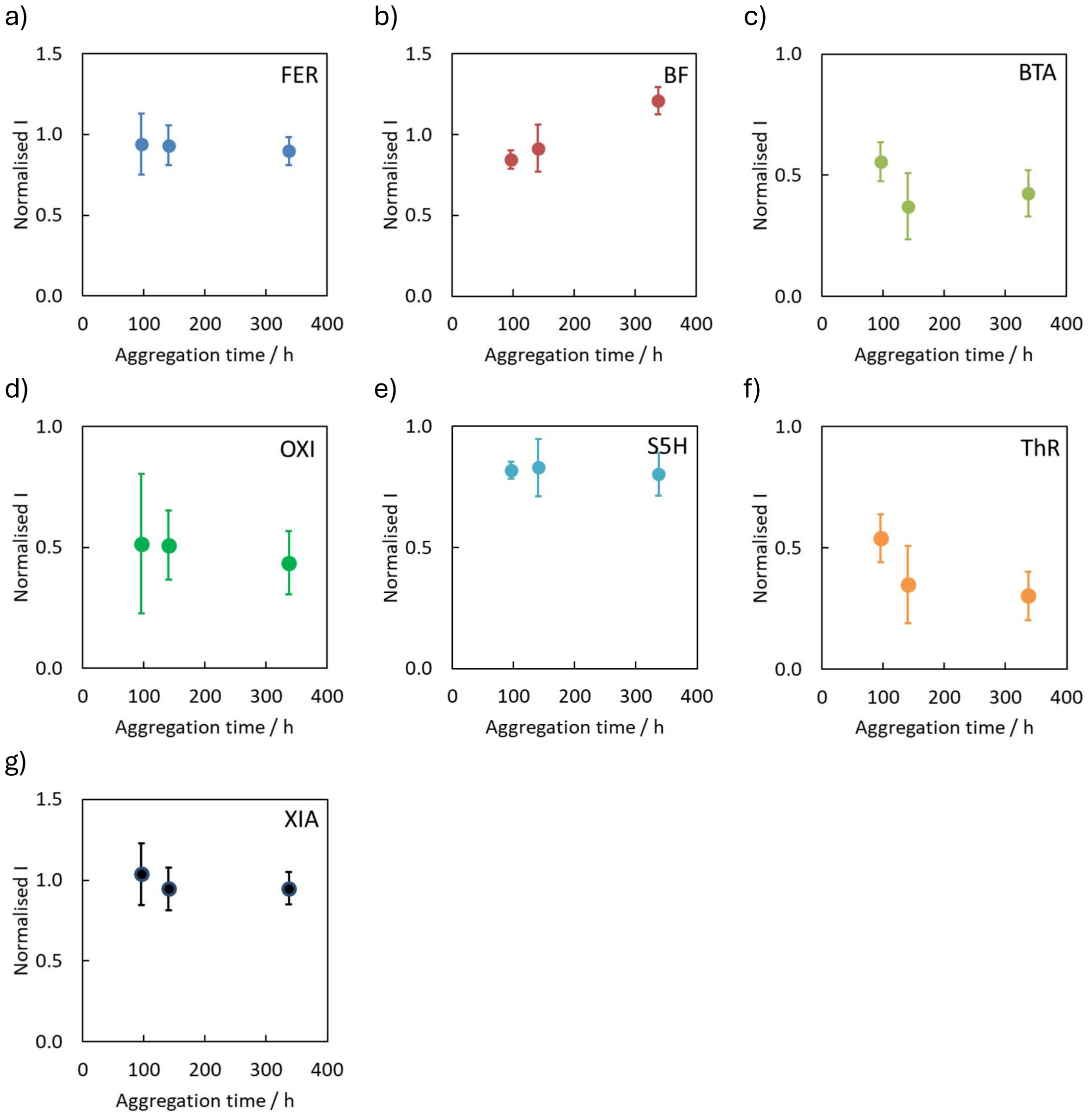

**Figure S58**. Fluorescence measurements for the displacement of **IND** ($\lambda_{ex}$ = 430 nm, $\lambda_{em}$ = 592-598 nm) by competing ligands from αSyn fibrils at different aggregation timepoints. Aggregation was performed using 50 μM αSyn in 50 mM Tris with 1.5 mM $NaN_3$ (pH 7.4, 37 °C). The reaction was monitored by diluting aliquots to 1.0 μM αSyn in 1xPBS (pH 7.4, 25 °C) and adding 2.0 μM reporting ligand and 5.0 μM competing ligand. Data points are the average of at least three measurements with 95% confidence intervals shown.

*Ligand Profiling After Storage*

The fluorescence measured for each reporting ligand (2.0 μM) bound to 96 h αSyn aggregates (1.0 μM) in 1xPBS (pH 7.4) after being stored at -79 °C, 4 °C, or 37 °C for 25 days, and in the absence of any competing ligand, is shown in Figure S59. Measurements were made using a plate reader. These data were normalised to the fluorescence intensity measured for the reporting ligand bound to 96 h αSyn aggregates pre-storage without any competing ligand present.

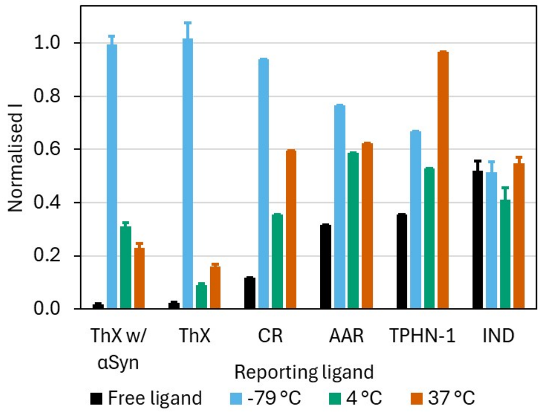

**Figure S59**. Fluorescence emission of different reporting ligands bound to 96 h αSyn aggregates that had been stored for 25 days at -79 °C, 4 °C, or 37 °C. Fluorescence data are normalised to the signal measured for the reporting ligands bound to 96 h αSyn aggregates pre-storage. Datapoints are the average of at least three measurements, with 95% confidence intervals shown.

The fluorescence measured for each reporting ligand (2.0 μM) bound to 96 h αSyn aggregates (1.0 μM) in 1xPBS (pH 7.4) after being stored at -79 °C, 4 °C, or 37 °C for 25 days, in the presence of each competing ligand (5.0 μM), is shown in Figure S60 for **ThX**, Figure S61 for **ThX** when αSyn aggregates formed in the presence of **ThX** are used, Figure S62 for **CR**, Figure S63 for **TPHN-1**, and Figure S64 for **AAR**. Measurements were made using a plate reader. These data were normalised to the fluorescence intensity measured for the reporting ligand bound to the respective fibril in the absence of any competing ligand. No measurements were made for **IND** in the presence of competing ligand given the absence of fluorescence measured when **IND** is combined with the stored fibrils.

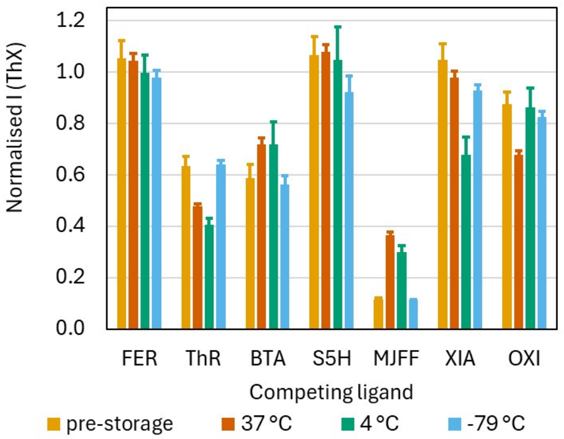

**Figure S60**. Fluorescence emission of **ThX** (2.0 μM) in the presence of competing ligand (5.0 μM) bound to 96 h αSyn aggregates before storage, and after storage for 25 days at 37 °C, 4 °C, or -79 °C. Data for each fibril target are normalised to the fluorescence emission measured in the absence of any competing ligand. Datapoints are the average of at least three measurements, with 95% confidence intervals shown.

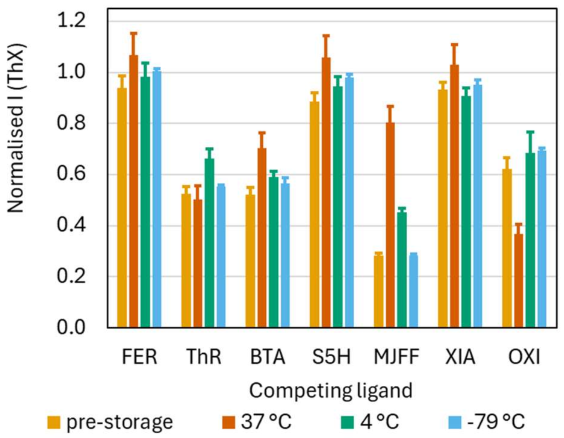

**Figure S61**. Fluorescence emission of **ThX** (2.0 μM) in the presence of competing ligand (5.0 μM) bound to 96 h αSyn aggregates (formed in the presence of 20 μM **ThX**) before storage and after storage for 25 days at 37 °C, 4 °C, or -79 °C. Data for each fibril target are normalised to the fluorescence emission measured in the absence of any competing ligand. Datapoints are the average of at least three measurements, with 95% confidence intervals shown.

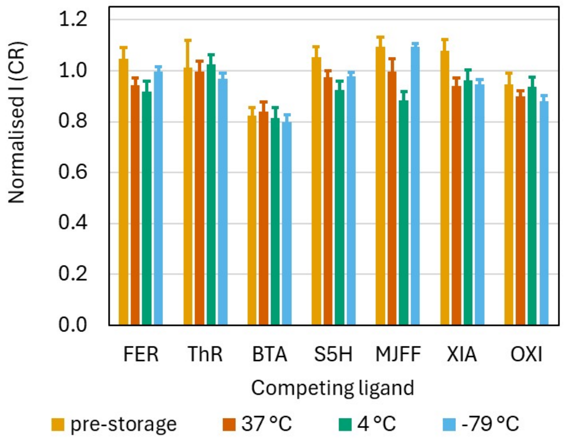

**Figure S62**. Fluorescence emission of **CR** (2.0 μM) in the presence of competing ligand (5.0 μM) bound to 96 h αSyn aggregates before storage, and after storage for 25 days at 37 °C, 4 °C, or -79 °C. Data for each fibril target are normalised to the fluorescence measured in the absence of any competing ligand. Datapoints are the average of at least three measurements, with 95% confidence intervals shown.

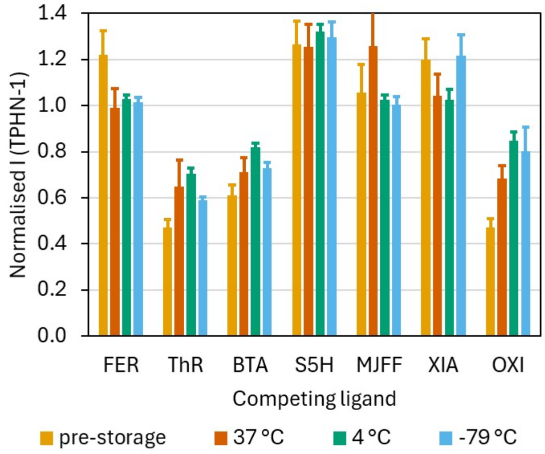

**Figure S63.** Fluorescence emission of **TPHN-1** (2.0 μM) in the presence of competing ligand (5.0 μM) bound to 96 h αSyn aggregates before storage, and after storage for 25 days at 37 °C, 4 °C, or -79 °C. Data for each fibril target are normalised to the fluorescence emission measured in the absence of any competing ligand. Datapoints are the average of at least three measurements, with 95% confidence intervals shown.

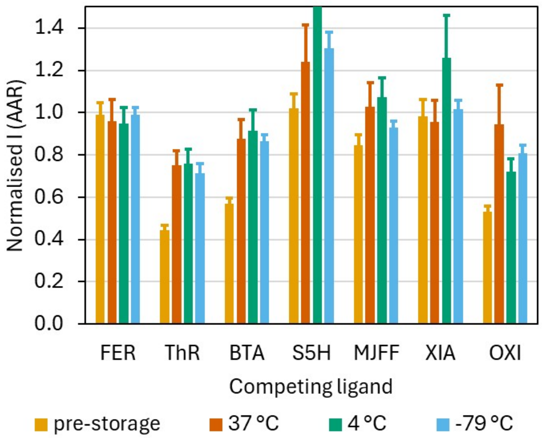

**Figure S64**. Fluorescence emission of **AAR** (2.0 μM) in the presence of competing ligand (5.0 μM) bound to 96 h αSyn aggregates before storage, and after storage for 25 days at 37 °C, 4 °C, or -79 °C. Data for each fibril target are normalised to the fluorescence emission measured in the absence of any competing ligand. Datapoints are the average of at least three measurements, with 95% confidence intervals shown.

#### *Ligand Profiling of Incubated Samples*

Aliquots of the 96 h αSyn aggregation mixture incubated with **ThX**, **CR**, **TPHN-1**, **AAR**, and **IND** at 37 °C for 25 days were diluted to a final concentration of 1.0 μM αSyn with 2.0 μM of the respective reporting ligand and optionally 5.0 μM competing ligand in 1xPBS (pH 7.4). The fluorescence of the resultant solution was measured on the plate reader. These data were normalised to the fluorescence intensity measured for the reporting ligand bound to the incubated αSyn aggregates in the absence of any competing ligands.

The fluorescence measurements for fibrils incubated with **ThX** are shown in Figure S65, with **CR** are shown in Figure S66, with **AAR** are shown in Figure S67, with **TPHN-1** are shown in Figure S68, and with **IND** are shown in Figure S69.

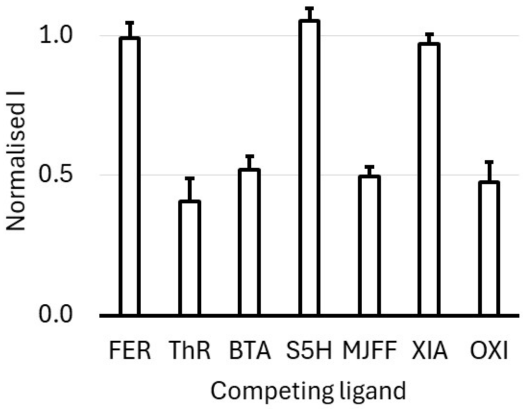

**Figure S65**. Fluorescence emission of **ThX** (2.0 μM) in the presence of competing ligand (5.0 μM), bound to 96 h αSyn aggregates that have been incubated with **ThX** (20 μM) for 25 days at 37 °C. Data are normalised to the fluorescence emission measured in the absence of any competing ligands. Datapoints are the average of at least three measurements, with 95% confidence intervals shown.

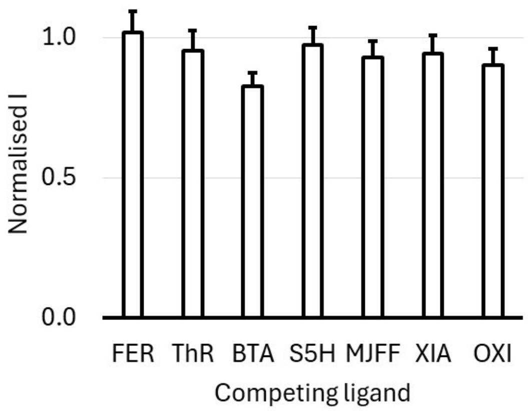

**Figure S66**. Fluorescence emission of **CR** (2.0 μM) in the presence of competing ligand (5.0 μM), bound to 96 h αSyn aggregates that have been incubated with **CR** (20 μM) for 25 days at 37 °C. Data are normalised to the fluorescence emission measured in the absence of any competing ligands. Datapoints are the average of at least three measurements, with 95% confidence intervals shown.

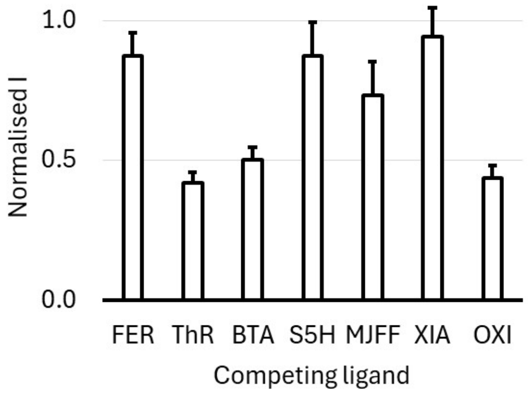

**Figure S67**. Fluorescence emission of **AAR** (2.0 μM) in the presence of competing ligand (5.0 μM), bound to 96 h αSyn aggregates that have been incubated with **AAR** (20 μM) for 25 days at 37 °C. Data are normalised to the fluorescence emission measured in the absence of any competing ligands. Datapoints are the average of at least three measurements, with 95% confidence intervals shown.

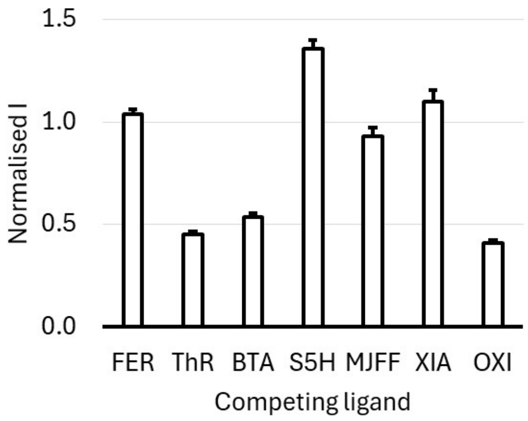

**Figure S68**. Fluorescence emission of **TPHN-1** (2.0 μM) in the presence of competing ligand (5.0 μM), bound to 96 h αSyn aggregates that have been incubated with **TPHN-1** (20 μM) for 25 days at 37 °C. Data are normalised to the fluorescence emission measured in the absence of any competing ligands. Datapoints are the average of at least three measurements, with 95% confidence intervals shown.

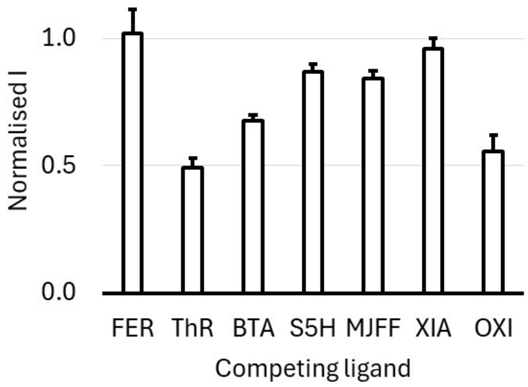

**Figure S69**. Fluorescence emission of **IND** (2.0 μM) in the presence of competing ligand (5.0 μM), bound to 96 h αSyn aggregates that have been incubated with **IND** (20 μM) for 25 days at 37 °C. Data are normalised to the fluorescence emission measured in the absence of any competing ligands. Datapoints are the average of at least three measurements, with 95% confidence intervals shown.

*Ligand Profiling after Long-Term Storage*

Aliquots of the stored αSyn aggregates (1.0 μM) were combined with reporting ligand (2.0 μM) and optionally competing ligand (5.0 μM) in 1xPBS (pH 7.4). The fluorescence of the resultant solution was measured on the plate reader. These data were normalised to the fluorescence intensity measured for the reporting ligand bound to the incubated αSyn aggregates in the absence of any competing ligands.

The fluorescence measurements for fibrils incubated with **ThX** are shown in Figure S70, with **CR** are shown in Figure S66, with **AAR** are shown in Figure S72, and with **TPHN-1** are shown in Figure S73.

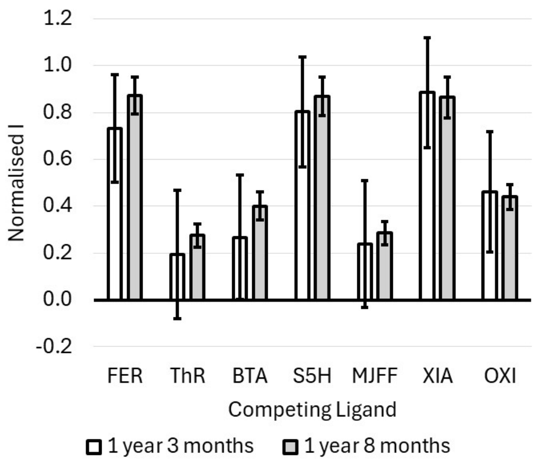

**Figure S70**. Fluorescence emission of **ThX** (2.0 μM) in the presence of competing ligand (5.0 μM), bound to αSyn aggregates that have been stored for either 1 year and 3 months, or 1 year and 8 months. Data are normalised to the fluorescence emission measured in the absence of any competing ligands. Datapoints are the average of at least three measurements, with 95% confidence intervals shown.

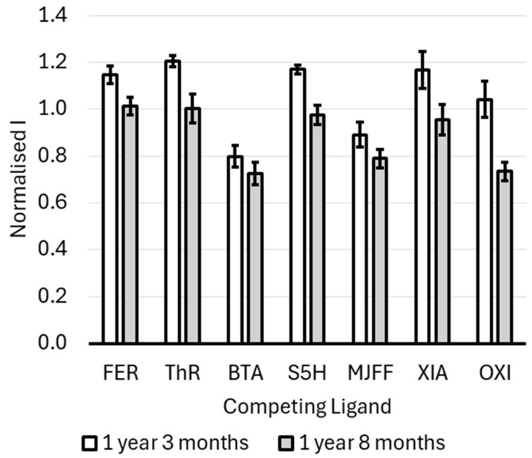

**Figure S71**. Fluorescence emission of **CR** (2.0 μM) in the presence of competing ligand (5.0 μM), bound to αSyn aggregates that have been stored for either 1 year and 3 months, or 1 year and 8 months. Data are normalised to the fluorescence emission measured in the absence of any competing ligands. Datapoints are the average of at least three measurements, with 95% confidence intervals shown.

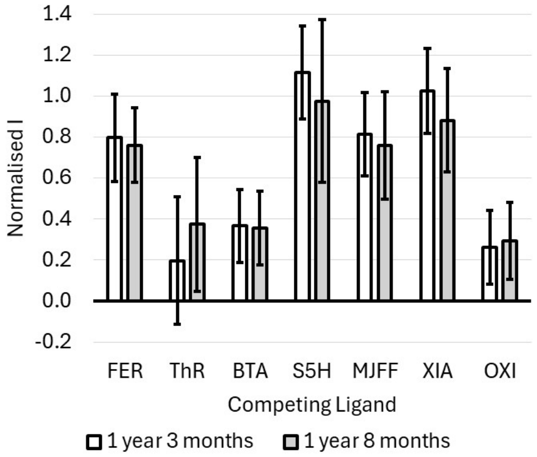

**Figure S72**. Fluorescence emission of **AAR** (2.0 μM) in the presence of competing ligand (5.0 μM), bound to αSyn aggregates that have been stored for either 1 year and 3 months, or 1 year and 8 months. Data are normalised to the fluorescence emission measured in the absence of any competing ligands. Datapoints are the average of at least three measurements, with 95% confidence intervals shown.

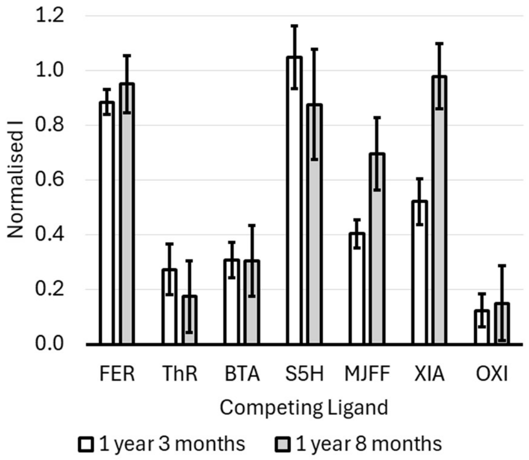

**Figure S73**. Fluorescence emission of **TPHN-1** (2.0 μM) in the presence of competing ligand (5.0 μM), bound to αSyn aggregates that have been stored for either 1 year and 3 months, or 1 year and 8 months. Data are normalised to the fluorescence emission measured in the absence of any competing ligands. Datapoints are the average of at least three measurements, with 95% confidence intervals shown.

*Ligand Profiling after Freeze-Thaw Cycles*

Aliquots of αSyn aggregates (1.0 μM) after a given number of freeze-thaw cycles were combined with reporting ligand (2.0 μM) and optionally competing ligand (5.0 μM) in 1xPBS (pH 7.4). The fluorescence of the resultant solution was measured on the plate reader. These data were normalised to the fluorescence intensity measured for the reporting ligand bound to the incubated αSyn aggregates in the absence of any competing ligands.

The fluorescence measurements for fibrils incubated with **ThX** are shown in Figure S74, with **CR** are shown in Figure S75 , with **AAR** are shown in Figure S76, and with **TPHN-1** are shown in Figure S77.

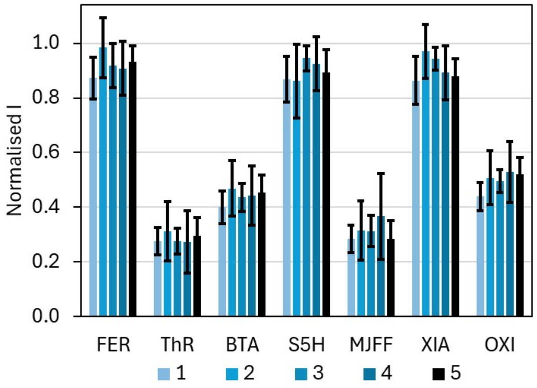

**Figure S74**. Fluorescence emission of **ThX** (2.0 μM) in the presence of competing ligand (5.0 μM), bound to αSyn aggregates that have been stored for 1 year and 8 months and subjected to one to five freeze-thaw cycles. Data are normalised to the fluorescence emission measured in the absence of any competing ligands. Datapoints are the average of at least three measurements, with 95% confidence intervals shown.

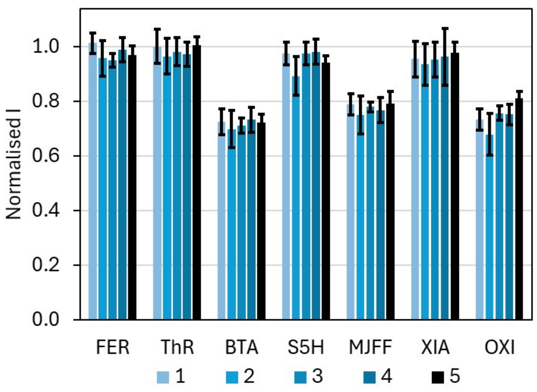

**Figure S75**. Fluorescence emission of **CR** (2.0 μM) in the presence of competing ligand (5.0 μM), bound to αSyn aggregates that have been stored for 1 year and 8 months and subjected to one to five freeze-thaw cycles. Data are normalised to the fluorescence emission measured in the absence of any competing ligands. Datapoints are the average of at least three measurements, with 95% confidence intervals shown.

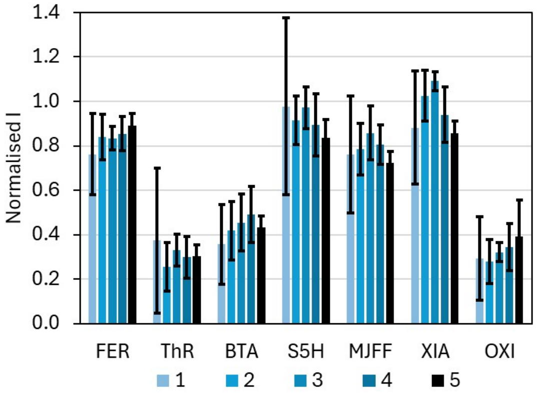

**Figure S76**. Fluorescence emission of **AAR** (2.0 μM) in the presence of competing ligand (5.0 μM), bound to αSyn aggregates that have been stored for 1 year and 8 months and subjected to one to five freeze-thaw cycles. Data are normalised to the fluorescence emission measured in the absence of any competing ligands. Datapoints are the average of at least three measurements, with 95% confidence intervals shown.

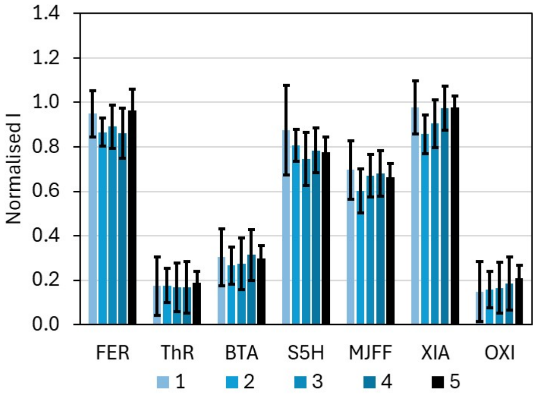

**Figure S77**. Fluorescence emission of **TPHN-1** (2.0 μM) in the presence of competing ligand (5.0 μM), bound to αSyn aggregates that have been stored for 1 year and 8 months and subjected to one to five freeze-thaw cycles. Data are normalised to the fluorescence emission measured in the absence of any competing ligands. Datapoints are the average of at least three measurements, with 95% confidence intervals shown.
