## Supporting code and .csv files for "The Morphology of α-Synuclein Fibrils Changes during Formation, Storage, and upon Exposure to Ligands": Chemical Space.ipynb

```
# %% [markdown]
## VISUALISATION OF $\alpha$SYN LIGAND CHEMICAL SPACE

This code extracts $\alpha$Syn ligands from an input database and viualises the chemical space.
1. Importants and filters an input ligand database, selecting only $\alpha$Syn ligands with dissociation constants < 1.0 uM
2. Generates Morgan fingerprints from the ligand SMILES strings
3. Reduces dimensionality using PCA followed by t-SNE
4. Interactively plots the chemical space, and highlights the compounds selected for the panel

# %% [code]
### === IMPORT PACKAGES ===
import pandas as pd
import matplotlib.pyplot as plt
import numpy as np
import math
from math import log
import plotly.express as px
from rdkit import Chem
from rdkit.Chem import AllChem
from rdkit.Chem import rdFingerprintGenerator
from sklearn.manifold import TSNE
from sklearn.decomposition import PCA
import csv

# %% [code]
### === IMPORT DATABASE AS .CSV ===
file_path = r'...//Ligand Database.csv'
df_in = pd.read_csv(file_path)

### === GLOBAL VARIABLES ===
RAND_SEED = 42

### === GET DESIRED DATA ===
# We want only data for ligands with dissociation constants below 1.0 uM
df_aS = df_in[df_in['Target fibril'] == 'aS'] # get aSyn data
df_aS_unique = df_aS.drop_duplicates(subset='InChIiso', keep='last') # remove duplicates
df_aS_K = df_aS_unique[df_aS_unique['log(Kd/M)'].notna()].reset_index(drop=True) # keep only rows with valid dissociation constants
df_aS_nM = df_aS_K[df_aS_K['log(Kd/M)'].astype(float)<-6].reset_index(drop=True) # get only data with dissociation constants < 1.0 uM


# %% [code]
def fingerprints_from_smiles(smiles):
    """
    Generate a 2048-bit Morgan fingerprint (radius = 2) from a SMILES string.

    Parameters:
        smiles (str): SMILES string of a compound

    Returns:
        fp: Morgan Fingerprint of the SMILES string
    """
    mol = Chem.MolFromSmiles(smiles)
    if mol is None:
        raise ValueError(f"Invalid SMILES: {smiles}")
    fp = AllChem.GetMorganFingerprintAsBitVect(mol, 2, nBits=2048) 
    return fp

def tsne_df(mol_list):
    """
    Reduce a list of chemical fingerprints to 2D using PCA (50D) followed by t-SNE.
    
    Parameters:
        mol_list (array-like): List/array of molecular representations.

    Returns:
        pd.DataFrame: DataFrame with two t-SNE components.
    """
    pca = PCA(n_components=50,random_state=RAND_SEED)
    pca_result = pca.fit_transform(mol_list)
    tsne = TSNE(n_components=2, verbose=0, perplexity=50, n_iter=300, random_state=RAND_SEED)
    tsne_result = tsne.fit_transform(pca_result)
    return pd.DataFrame(tsne_result, columns=["Component 1", "Component 2"])

# %% [code]
def graph_chemical_space(df_in, smiles_col="SMILES", id_col=None, graph_title=None):
    """
    Visualizes chemical space using 2D t-SNE projection of Morgan fingerprints.

    Parameters:
        df_in (pd.DataFrame): Input DataFrame with SMILES and optional metadata.
        smiles_col (str): Column name with SMILES strings.
        id_col (str or None): Optional column for hover info.
        graph_title (str or None): Optional title of the output graph

    Returns:
        fig (plotly.graph_objects.Figure): Scatter plot of chemical space.
        rd_df (pd.DataFrame): Reduced-dimension DataFrame used for plotting.
    """
    
    df = df_in.reset_index(drop=True)

    mol_list = [fingerprints_from_smiles(smiles) for smiles in df[smiles_col]] # Compute fingerprints
    
    ### === DIMENSION REDUCTION ===
    rd_df = tsne_df(mol_list)
    rd_df = rd_df.join(df[[smiles_col]])

    ### === PANEL IDENTIFICATION ===
    # colour the panel ligands, if available (panel ligands are identified in the input .csv file)
    rd_df = rd_df.join(df['Panel'].astype(str))
    if id_col is not None:
        rd_df = rd_df.join(df[id_col])
    
    # Plot with panel ligands
    fig = px.scatter(
        rd_df, x='Component 1', y='Component 2', color='Panel', 
        title=graph_title or "Morgan Fingerprints with t-SNE",
        hover_data=[id_col] if id_col is not None else [smiles_col],
        width=800, height=800, template='simple_white'
    )
    
    fig.update_layout(
        font=dict(size=24),
        xaxis=dict(range=[-20, 20], dtick=10),
        yaxis=dict(range=[-20, 20], dtick=10),
        shapes=[dict(
            type="rect", xref="paper", yref="paper",
            x0=0, y0=0, x1=1, y1=1,
            line=dict(color="black", width=2),
            fillcolor="rgba(0,0,0,0)"
        )],
        paper_bgcolor="white",
        plot_bgcolor="white"
    )
    
    fig.update_traces(marker=dict(size=10, opacity=0.8))
    
    return fig, rd_df

# %% [code]
### === CREATE PLOT ===
fig_panel, rd_df = graph_chemical_space(df_in=df_aS_nM, smiles_col='SMILES',  id_col = "Compound")
fig_panel.show()

# %% [code]
```
