## Supporting code and .csv files for "The Morphology of α-Synuclein Fibrils Changes during Formation, Storage, and upon Exposure to Ligands": Tanimoto Heatmap.ipynb

```
# %% [markdown]
## TANIMOTO SIMILARITY COEFFICIENTS

Calculates the Tanimoto similarity coefficients between panel ligands

# %% [code]
### === IMPORT PACKAGES ===
import pandas as pd
import numpy as np
import seaborn as sns
import matplotlib.pyplot as plt
from rdkit import Chem
from rdkit.Chem import AllChem, DataStructs

# %% [code]
### === IMPORT PANEL LIGANDS IN .CSV ===
file_path = r'...//Panel Ligands.csv'
df_in = pd.read_csv(file_path)


# %% [code]
def calculate_tanimoto_similarity(smiles_list):
    """
    Compute the Tanimoto similarity matrix for a list of SMILES strings.
    
    Parameters:
        smiles_list (list of str): List of SMILES strings.

    Returns:
        np.ndarray: Symmetric matrix of Tanimoto similarity scores.
    """
    mols = [Chem.MolFromSmiles(SMILES) for SMILES in smiles_list] # generate RDKit molecule objects
    # check for invalid SMILES    
    if any(m is None for m in mols):
        raise ValueError("Invalid SMILES found in input.")
        
    fingerprints = [AllChem.GetMorganFingerprintAsBitVect(mol, radius=2, nBits=2048) for mol in mols] # Generate fingerprints
    
    ### === CALCULATE TANIMOTO SIMILARITY MATRIX ===
    n = len(fingerprints)
    similarity_matrix = np.zeros((n, n))
    for i in range(n):
        for j in range(n):
            similarity_matrix[i, j] = DataStructs.TanimotoSimilarity(fingerprints[i], fingerprints[j])
    return similarity_matrix

# Compute similarity matrix
similarity_matrix = calculate_tanimoto_similarity(df_in['SMILES'])

# Plot heatmap
plt.figure(figsize=(8, 6))
sns.heatmap(similarity_matrix, annot=False, cmap='Reds', xticklabels=df_in['Ligand'], yticklabels=df_in['Ligand'],
           cbar_kws={'label': 'Tanimoto similarity coefficient', 'shrink': 0.8})

# Format axes
cbar = plt.gca().collections[0].colorbar  
cbar.ax.tick_params(labelsize=20)
cbar.ax.set_ylabel('Tanimoto similarity coefficient', fontsize=20)  
plt.xticks(fontsize=20) 
plt.xticks(rotation=90)
plt.yticks(fontsize=20) 

plt.show()

# %% [code]
```
